# Transcriptomic plasticity is a hallmark of metastatic pancreatic cancer

**DOI:** 10.1101/2025.02.28.640922

**Authors:** Alejandro Jiménez-Sánchez, Sitara Persad, Akimasa Hayashi, Shigeaki Umeda, Roshan Sharma, Yubin Xie, Arnav Mehta, Wungki Park, Ignas Masilionis, Tinyi Chu, Feiyang Zhu, Jungeui Hong, Ronan Chaligne, Eileen M. O’Reilly, Linas Mazutis, Tal Nawy, Itsik Pe’er, Christine A. Iacobuzio-Donahue, Dana Pe’er

## Abstract

Metastasis is the leading cause of cancer deaths; nonetheless, how tumor cells adapt to vastly different organ contexts is largely unknown. To investigate this question, we generated a transcriptomic atlas of primary tumor and diverse metastatic samples from a patient with pancreatic ductal adenocarcinoma who underwent rapid autopsy. Unsupervised archetype analysis identified both shared and site-specific gene programs, including lipid metabolism and gastrointestinal programs prevalent in peritoneum and stomach wall lesions, respectively. We developed a probabilistic approach for inferring clonal phylogeny from single-cell and matched whole-exome data. Distantly related genetic clones in the peritoneum express the lipid metabolism program, likely due to signaling by the adipocyte-rich peritoneum environment, and cells in most clones express multiple programs, suggesting that transcriptomic plasticity is a prevalent feature of metastatic cells. These deeply annotated analyses using a patient-centric platform provide a model for investigating metastatic mechanisms and plasticity in advanced cancer.

## Introduction

Metastasis is a systemic disease responsible for the majority of cancer-related deaths^1,2^, yet our understanding of how tumor cells disseminate and thrive in distant tissues remains limited. To metastasize, cancer cells must overcome many hurdles, including the need to escape from the tissue of origin, migrate, evade immune surveillance and invade distant tissue^3,4^. The microenvironments of different organs each pose additional adaptive challenges for cancer cell colonization. For some tumor types, selection may act on intratumor genetic heterogeneity to shape these adaptive processes^5,6^, whereas for others, genomic studies have uncovered few recurrent mutations associated with specific metastatic behaviors or organotropism^7–9^. More recently, epigenetic plasticity has emerged as a hallmark of cancer, which confers the ability to reinvent cellular phenotypes and drive phenotypic heterogeneity in the service of adaptation^10–13^. How this plasticity manifests at the molecular level, the extent to which it shapes tumor progression^14^, and its relevance to treatment^15^ are major open questions.

Pancreatic ductal adenocarcinoma (PDAC) exhibits particularly low heterogeneity in driver mutations, which tend to be shared across primary and metastatic sites^9,16^, underscoring the need to identify alternate adaptive mechanisms. Advanced tumors are not commonly resected and metastases are rarely biopsied sequentially, making it difficult to reconstruct tumor progression and providing scant information on metastases in some organs. Rapid autopsy offers a critical opportunity for systematically investigating shared and organ-specific metastatic programs in multiple lesions derived from a single germline^17^. The ability to collect multiple independent metastases from a single organ also provides an unparalleled approximation of a controlled biological replicate in human cancer. Such post-mortem sampling, coupled with genotyping and lineage reconstruction, recently provided insights into modes of evolution and metastatic seeding in PDAC^18^.

To gain insights into the molecular mechanisms of adaptation in this patient-centric view, however, requires a combination of clonal lineage information and deep phenotypic profiling. Single-cell gene expression data provides rich phenotypic information at the cellular level, but it is problematic for clonal and phylogenetic reconstruction, whereas simultaneously sequencing single-cell DNA can provide genotype information but does not scale sufficiently. Typical phenotypic analyses are also not designed to find adaptive gene programs. Computational approaches are thus needed to overcome these challenges, and to enable the comparison of clonal lineage and molecular phenotypes in a single cancer across multiple lesions and organs.

We collected two primary and nine metastatic tumors from a patient with PDAC who underwent a rapid autopsy, and subjected the samples to single-nucleus RNA sequencing (snRNA-seq), recovering the transcriptomes of over 45,000 cancer epithelial cells. Using archetypal analysis, we identified adaptive gene programs that are missed by standard clustering. To investigate the evolutionary dynamics of metastatic PDAC, we developed IntegrateCNV, an approach to robustly infer copy number alterations (CNAs) from snRNA-seq and matching bulk whole exome sequencing data, and PICASSO, a method to identify cell clones and generate clonal phylogenies using potentially noisy single-cell CNA profiles. We find evidence of strong adaptation to local organ microenvironment, including metabolic rewiring of peritoneal lesions—a very common but little-studied site of metastasis in PDAC—as well as multiple different shared epithelial–mesenchymal transition programs. Our work identifies plasticity as the major force in PDAC metastatic adaptation, and provides approaches for deep phenotypic and phylogenetic analysis from single-cell expression data.

## Introduction

### A patient-specific atlas of PDAC metastasis

Using rapid autopsy specimens from a single patient with PDAC and thoroughly optimized specimen dissociation and scRNA-seq protocols, we constructed a comprehensive atlas spanning primary and metastatic sites, enabling the study of how cancer evolves and adapts across diverse tissue environments. We integrated snRNA-seq and matched WES data from each specimen to uncover both clonal architecture and adaptive transcriptional programs driving metastatic progression.

The patient was diagnosed at age 35 with PDAC and extensive synchronous liver metastases, as evidenced by computed tomography, which was used in addition to CA19-9 tumor marker levels to follow disease status over the 9 months that the patient survived (**Fig. 1a,b**). Despite initial robust response to standard-of-care modified FOLFIRINOX (5-fluorouracil, leucovorin, irinotecan, and oxaliplatin), the rapid emergence of refractory disease, unresponsive to second-line gemcitabine + nab-paclitaxel, highlighted the cancer’s remarkable adaptive capacity within months of treatment.

**Figure 1.**
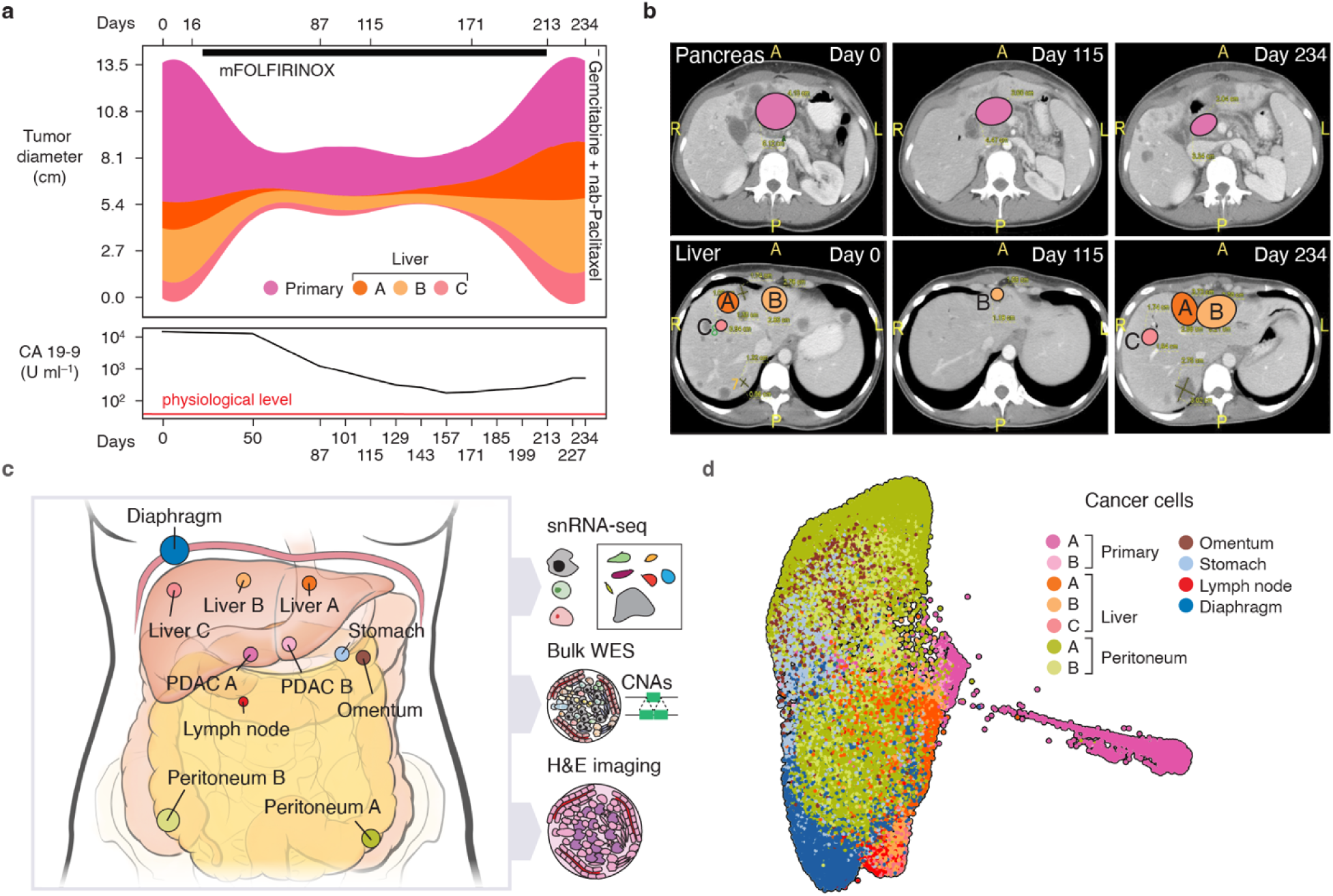
Profile of a cancer ecosystem from a single PDAC patient. **a**, Top, maximal diameter of primary and liver tumors, based on CT measurements at the indicated time points from diagnosis (day 0). Black bar marks the period of mFOLFIRINOX treatment. Bottom, levels of CA 19-9 tumor marker in blood, based on indicated measurement days. Baseline at diagnosis (day 0) is 13,000 U ml^−1^ and upper physiological limit is 37 U ml^−1^ (red line). **b**, Representative CT scans. Primary and liver metastatic tumors are overdrawn with colored ellipses. L, left; R, right; A, anterior; P, posterior. **c**, Anatomical location of collected biospecimens used to generate matched snRNA-seq, WES, and hematoxylin and eosin (H&E) data. Circle diameter indicates relative tumor size. **d**, Force-directed layout (FDL) of cancer cell transcriptomes (45,134 nuclei), colored by sample (Methods). Stomach refers to stomach wall metastasis.

Through the rapid autopsy, we collected 11 tumor specimens representing diverse tissue microenvironments, including the pancreas and six distal organs. The sampling included anatomically separate lesions from the same organ (the best approximation of biological replicates in human cancer) where possible: two peritoneal and three liver metastatic samples, in addition to two regions of the primary tumor (**Fig. 1c**). These 11 samples, collected from 7 distinct organ sites, exhibit diverse cell-type compositions and tissue morphologies (**Extended Data Fig. 1**).

We recovered 73,142 high-quality snRNA-seq profiles from all samples, organized into 39 clusters by PhenoGraph^19^, which we annotated based on known marker genes (**Extended Data Fig. 1b–d**, **Supplementary Table 1** and Methods). To distinguish cancer cells from non-cancer, we identified cells with high *KRAS* signaling^20,21^ and detected clusters with accumulated CNAs using inferCNV^22^ (**Extended Data Fig. 2a,b** and Methods). In total, we recovered 45,134 cancer epithelial nuclei across all lesions, bearing multiple cancer-related mutations (**Fig. 1d** and **Extended Data Fig. 2c,d**).

### PICASSO resolves single-cell phylogenies

The availability of both primary and metastatic cells from the same patient provides a unique opportunity to study how cancer cells evolve and adapt to different tissue environments. To dissect the relative roles of genetic mutations and epigenetic plasticity in metastatic adaptation, it is essential to reconstruct the evolutionary history of cancer cells and compare their genotypic and phenotypic characteristics within a shared phylogenetic framework. However, current approaches face significant limitations.

Bulk whole exome sequencing offers a coarse view of phylogenetic relationships across lesions; however, it lacks single-cell resolution and cannot link genetic mutations to cellular phenotypes^23^. Combined DNA-RNA single-cell assays^24,25^ are limited by cost and throughput—published studies consist of too few cells (typically <1000)^24–27^ to capture the full phenotypic heterogeneity typically observed within lesions^28–31^. Although copy number inference from single-nucleus or single-cell RNA-seq (scRNA-seq) data^22,32^ can inform clonal relationships, current methods are extremely noisy and strongly impacted by confounding factors such as the influence of tumor cell state and its related gene expression patterns^33–37^. In addition, many phylogenetic algorithms assume that mutations occur only once (“perfect phylogeny”), whereas in cancer, CNAs are highly recurrent^38–41^. For example, over 50% of CNA regions violate the perfection assumption in our data, complicating traditional phylogenetic approaches (**Extended Data Fig. 3a** and Methods). Finally, classic algorithms for phylogenetic analysis assume evolutionary characters are reliable, whereas CNAs called from single-cell expression data are uncertain and noisy. Uncovering genotype–phenotype relationships and the role of epigenetic plasticity during cancer progression thus requires new approaches that can (1) reliably infer CNAs from scRNA-seq data, and (2) construct a robust phylogeny of cancer clones, taking into account noise, uncertainty and possible CNA recurrence, as well as the large scale of single-cell data.

To address these challenges, we instigated a two-step approach. First, we developed IntegrateCNV, a statistical framework that leverages matched bulk WES and snRNA-seq profiles to infer CNAs at single-cell resolution (**Extended Data Fig. 3b** and Methods). Unlike existing methods that infer CNAs genome-wide from scRNA-seq alone^32,42,43^, IntegrateCNV uses bulk WES data to identify regions harboring CNAs before performing targeted inference from scRNA-seq data for individual cells in these candidate regions. This focused strategy increases signal-to-noise by limiting analysis to regions with strong evidence of copy number variation. Specifically, for each cell and candidate region, it determines whether an alteration is likely to be present based on gene expression relative to a copy-neutral reference (**Extended Data Fig. 3b** and Methods). IntegrateCNV achieves higher or equal correlation with sample-level CNAs derived from bulk WES data compared to widely used tools such as inferCNV^22^ and CopyKat^32^ (**Extended Data Fig. 3c**).

While IntegrateCNV improves CNA detection accuracy, the profiles it generates still contain many errors (**Extended Data Fig. 3c**), especially false negatives. Unfortunately, even phylogenetic reconstruction methods that allow errors in the character matrix typically assume them to be minimal. Moreover, the few algorithms that infer phylogenies from single-cell CNA profiles are designed for small-scale single-cell DNA sequencing experiments and assume error-free input^44,45^. To overcome these challenges and construct robust phylogenies, we developed PICASSO (Phylogenetic Inference from Copy number Alterations in Single-cell Sequencing Observations), a maximum-likelihood method tailored to large-scale, noisy CNA profiles (**Fig. 2a**, **Extended Data Fig. 3d** and Methods). PICASSO employs a tree-recursive algorithm that starts with a single leaf node containing all cells, then iteratively decides whether to split each leaf into two subclones. Each decision to split is based on maximizing shared information in consensus CNA patterns, corrected for noise and missing values, using expectation–maximization. When there is insufficient evidence for further splitting, a leaf is marked terminal. The output of PICASSO is a probabilistic assignment of cells to clones, and a likelihood-optimized final tree describing clonal phylogenetic relationships and associated uncertainties. This top-down recursive approach only reconstructs major evolutionary relationships with good evidential support, and is substantially more robust to noisy data than standard bottom-up approaches.

**Figure 2.**
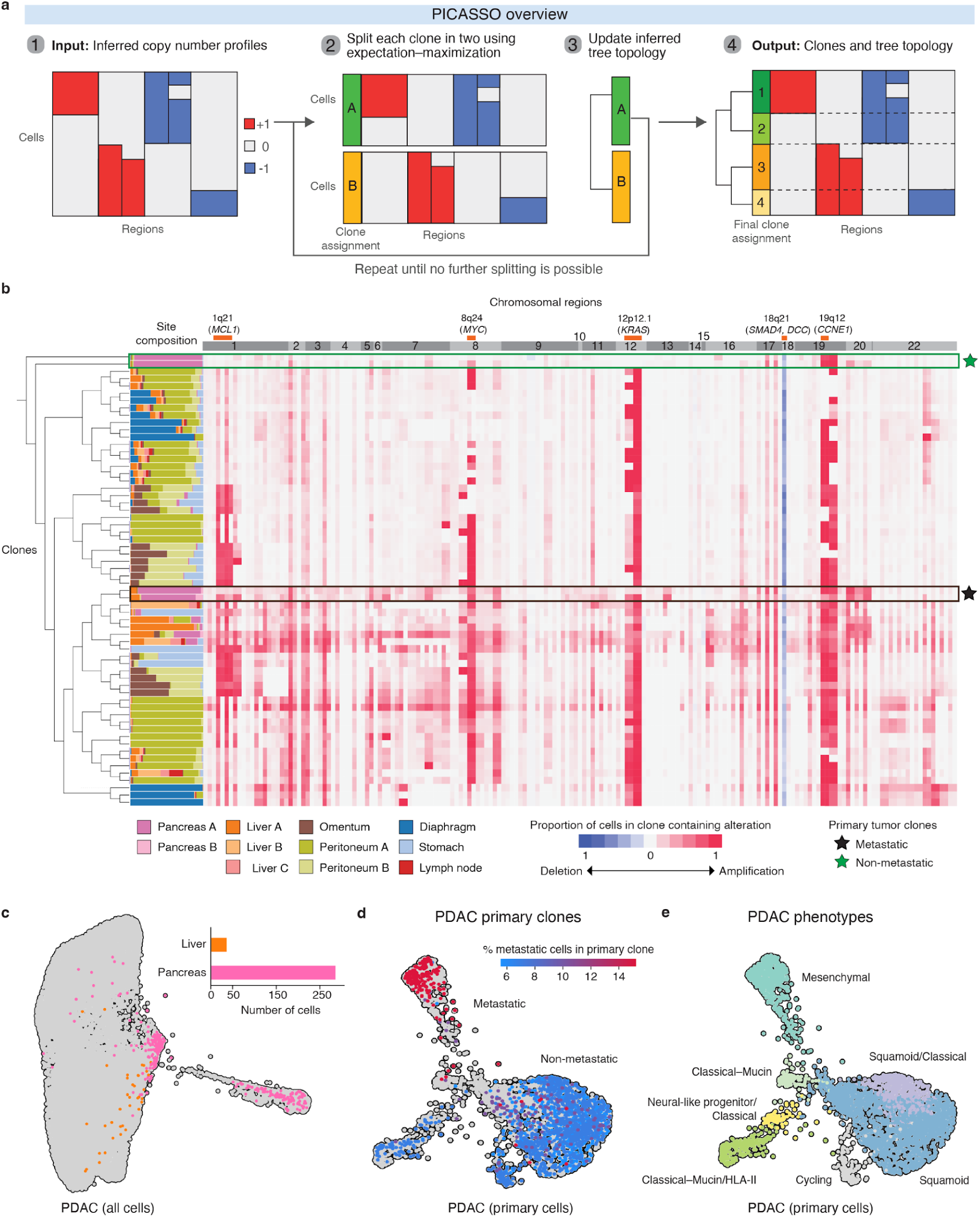
PICASSO generates a CNA-derived single-cell phylogeny. **a**, PICASSO takes CNA profiles from scRNA-seq data (inferred by IntegrateCNV, for example) as input and encodes them in a probabilistic manner, then iteratively splits clones into subclones based on clustering shared patterns by expectation–maximization (Methods). The algorithm proceeds in a top-down fashion until it reaches terminal leaves, which lack evidence for further splits. PICASSO output is the probabilistic assignment of cells to subclones and a maximum-likelihood-optimized tree. **b**, Phylogenetic relationships between clones (rows) derived from single-cell CNA profiles, for all cancer cells in the rapid autopsy dataset. Each stacked bar plot indicates the clone’s site composition (fraction of cells from each metastatic site), and the heatmap at right shows the modal copy numbers inferred by IntegrateCNV for that clone. The four clones that are predominantly from primary tumor (stars) are distinguished by whether they also contain cells in metastatic lesions. **c**, FDL of all cancer cells, colored by sample of origin for cells from the primary tumor clones (>50% cells from primary tumor) that also contain metastatic cells. Inset indicates the number of cells from each sample in these two clones. **d**,**e**, FDL of PDAC primary cells showing cancer clones colored by proportion of primary cells within the clones (**d**) and PDAC phenotypes (**e**). Gray cells in **d** were removed from phylogenetic analysis due to low transcript counts (Methods).

We validated performance using simulated data, which demonstrated that PICASSO produces more parsimonious phylogenies and outperforms agglomerative clustering in both speed and accuracy under varying levels of noise (**Extended Data Fig. 4a,b**). By providing a probabilistic assignment of cells to clones and a likelihood-optimized tree describing clonal relationships and uncertainties, PICASSO is thus an effective tool for dissecting the relationship between genotype and phenotype during cancer progression.

### Evolutionary reconstruction of metastatic PDAC

We applied IntegrateCNV to cancer cells in our metastatic PDAC dataset and used PICASSO on the resulting 45,134 single-cell copy number profiles in this large-scale data (**Fig. 2b** and Methods). Based on CNA calls in 116 candidate regions, PICASSO resolved 62 clones with a clear phylogenetic structure following noise removal. The resulting phylogeny is highly stable; despite the probabilistic nature of the algorithm, most evolutionary relationships are conserved across repeated runs (**Extended Data Fig. 4c**). Furthermore, bootstrapping analysis reveals that the tree structure remains stable even when removing a fraction of cells for each region (**Extended Data Fig. 4d**).

We used the inferred phylogeny to investigate patterns of metastasis, first asking which clones in the primary tumor spread, and why. Four primary clones, defined as containing at least 50% of cells from the primary tumor, were identified—two that metastasized, and two that did not (**Fig. 2b**). A subset of tumor cells from liver metastases were found to be closely related to the metastasizing clones from the primary tumor. Notably, liver-dominant clones are the most closely related metastatic clones to those found in the primary tumor, suggesting that the liver was the initial site of metastasis in this patient, consistent with the observation that PDAC typically spreads to the liver first^46–50^ (**Fig. 2b,c**).

Analysis of the metastasizing primary clones revealed distinct genomic and transcriptional features associated with metastatic potential. Metastatic clones from the primary had many more CNAs than their non-metastatic counterparts. Notably, amplification of the oncogenic *KRAS^G12V^* locus is a hallmark of nearly all metastatic clones. We recently showed that oncogenic *KRAS* enhances plasticity during PDAC premalignancy, partly by remodeling the communication between cancer cells and their environment^51^. Our findings suggest that oncogenic *KRAS* continues to drive plasticity in advanced disease, and that its amplification provides an additional boost that promotes metastatic competence.

In addition to genetic alterations, our dataset provides a rare opportunity to examine the transcriptional states of metastasizing clones. Mapping known PDAC tumor phenotypes revealed that most primary tumor cells from metastatic clones display a mesenchymal phenotype, indicating an epithelial-to-mesenchymal transition (EMT), which has been strongly associated with metastasis^52,53^ (**Fig. 2d,e** and Methods). In contrast, non-metastatic clones are enriched for epithelial phenotypes, suggesting that metastatic clones are already transcriptionally poised for dissemination while in the primary tumor. The observation of mesenchymal phenotypes in cells from non-metastatic clones signifies that EMT alone is insufficient for successful metastasis. This level of resolution is uniquely enabled by our approach, as it allows us to connect transcriptional phenotypes to phylogenetic patterns.

The phylogenetic tree provides insights into metastatic seeding and spread. While some clones map to a dominant metastatic site, most are found in multiple organs, suggesting that metastatic clones can adapt to diverse tissue environments. Conversely, each metastatic site contains cells from multiple clones, some separated by large phylogenetic distances (**Fig. 2b**), implying that metastatic sites were seeded by multiple clones in independent events. This and similar findings in other contexts^54,55^ support the idea that once tumor cells establish themselves at distal sites, they remodel the local microenvironment to create a favorable “soil” for further seeding by the primary tumor or other metastases^56–60^.

### Archetype analysis identifies metastatic gene programs

The observation that most clones metastasized to multiple organs raises the question of how tumor cells adapt to these distinct environments. While metastatic cells must overcome universal hurdles such as migration and extravasation to establish at distal locations^7^, each organ presents unique challenges requiring site-specific adaptations for successful colonization and growth. We reasoned that these adaptations should manifest as highly optimized transcriptional phenotypes, and that examining multiple metastatic sites from the same primary tumor would make it possible to uncover both shared and organ-specific mechanisms.

To systematically identify these adaptive programs, we applied archetype analysis^61–65^, which identifies boundary phenotypes known to represent optimized tasks, using a two-tiered approach (**Extended Data Fig. 5a,b** and Methods). Our strategy was to first analyze each organ separately, identifying four to six archetypes per tissue in a highly robust manner (**Extended Data Figs. 5c and 6a**). Archetype neighborhoods did not associate with cell-state density (**Extended Data Fig. 5c**), suggesting that archetype neighborhoods may represent both major cancer cell state phenotypes (high-density) and rare (low-density) cancer phenotype states^66^. Next, to find archetypes and programs that are potentially shared across organs, we integrated all 14,513 archetype-labeled cells (32% of all cancer cells) into a single matrix and applied graph-based clustering, yielding 19 archetype clusters (**Fig. 3a** and Methods). Finally, each archetype cluster was annotated using differentially expressed genes (DEGs) and Hotspot^67^ analysis (**Fig. 3b** and Methods).

**Figure 3.**
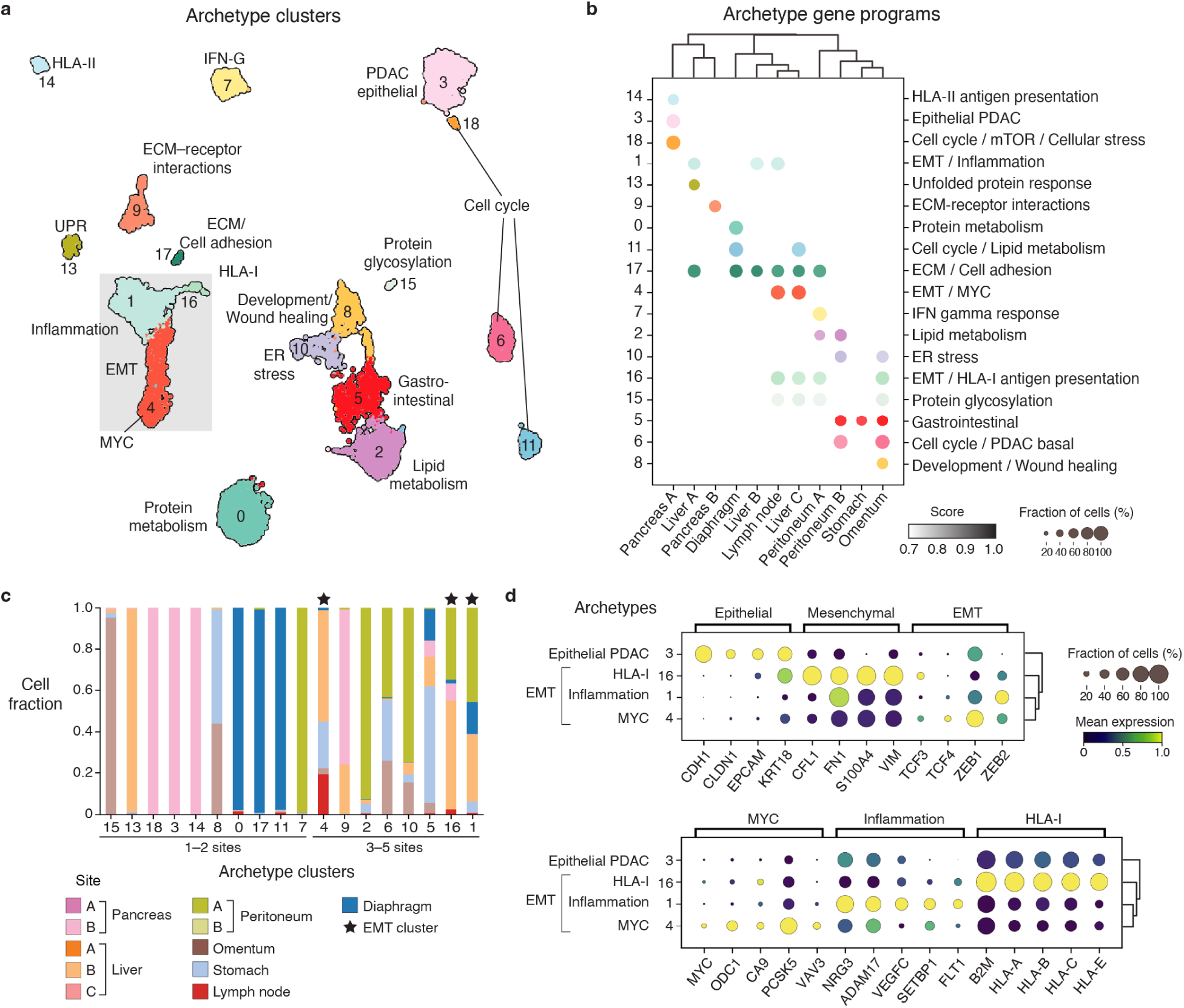
Archetype gene programs of primary and metastatic PDAC. **a**, UMAP of clustered archetypal cells from primary and metastatic sites, colored by cluster. Gray box encompasses three distinct archetype clusters related to EMT. **b**, Archetype gene program expression in each tumor sample. **c**, Fraction of cancer cells per archetype cluster, colored by sample. **d**, Expression of individual markers in archetypes 1, 3, 4 and 16. Archetype 3 corresponds to classical–squamous cells that are more epithelial, and is included for comparison. Canonical markers are indicated for epithelium, mesenchymal, and EMT, *MYC*, *MYC* targets and modules downstream of *MYC* signaling, inflammation, and HLA-I antigen processing and presentation.

Our analysis generated well-defined archetype clusters, including some that are unique to one organ and others that appear in multiple organs (**Fig. 3b,c**, **Extended Data Fig. 6b** and **Supplementary Tables 2–4**). For example, cells in archetype clusters 3, 9, 14 and 18 are only found in primary PDAC; cluster 13 (unfolded protein response: *HSP90AA1*, *HSPH1*, *HSPD1*, *DNAJA1*) is unique to the liver; and cluster 8 (development: *PBX1*, *HES1*, *PDGFB* and wound healing: *FOS*, *JUNB*, *NR4A1*, *ANGPTL4*) is specific to omentum. Additionally, cluster 5 (gastrointestinal: *MUC13*, *FABP1*, *FCGBP*) is found mostly in the stomach wall and cluster 2 (lipid metabolism: *HMGCS1*, *SQLE*, *FDPS*) is mainly in the peritoneum.

In contrast, we found that archetypes related to core cellular processes, such as cell cycle, migration, EMT, and cell–environment interactions, such as extracellular matrix (ECM) interactions and inflammation, are typically shared across multiple organs (**Fig. 3b,c**). To gain insight into biological functions that broadly contribute to metastatic capacity, we focused on archetype clusters 1, 4 and 16, which are present in multiple organs that together comprise all seven organ sites (**Fig. 3c**). These three clusters express mesenchymal genes^68–72^ and transcription factors related to EMT programs^73–76^ associated with metastatic spread^77,78^ (**Fig. 3d**). However, further analysis revealed that these apparently similar EMT states are distinguished by distinct gene and regulatory programs (**Supplementary Tables 3 and 4**).

Cluster 1 is enriched for programs associated with cytokine and chemokine secretion as well as TNF-α/NF-κB^79^ and IL-17 signaling, suggesting inflammatory activation. Cluster 16 is enriched for focal adhesion and ECM interactions. ECM remodeling is required for cancer cell growth and can recruit immune cells^56,80,81^, suggesting a potential role in establishing the metastatic niche. In contrast, cluster 4 is enriched for glucagon signaling, a common liver-expressed pathway^82^ that may reflect the influence of the liver microenvironment on cluster 4 cells, most of which originate in this organ (**Fig. 3c**). We found that cluster 4 has the highest expression of *MYC*, *MYC* target *ODC1*^83^, and *CA9,* which can be regulated by *MYC* under hypoxic conditions^84^, as well as genes downstream of *MYC* signaling (**Fig. 3d**), and is most enriched for MYC-expressing cells (**Extended Data Fig. 6c,d**). PDAC patient data and mouse models have linked *MYC* hyperactivation to more aggressive metastatic disease^85,86^ and chemoresistance^87–89^. Moreover, both *MYC* and EMT pathways are enriched in metastases compared to primary tumors in PDAC patients^86,90^. Further distinguishing these states, archetype cluster 1 expresses inflammatory genes, while cluster 16 expresses HLA-I antigen processing genes (**Fig. 3d**). Together, these findings reveal three distinct EMT phenotypes: archetypes 1 and 16 have a mesenchymal profile associated with inflammatory response, while archetype 4 has an EMT program that co-occurs with *MYC* signaling.

Unlike archetype analysis, traditional clustering approaches are not designed to identify gene programs optimized for specific biological tasks. Rather, they aim to define groups of cells that have more similar average expression than cells in other clusters. Direct comparison of these approaches in our dataset reveals substantial differences in cell groupings, DEGs between groups, and biological annotations (**Extended Data Fig. 6e–g**). While clustering detects broad processes such as EMT, proliferation, and stress (**Supplementary Table 5**), more specific adaptations to metastatic sites, such as lipid metabolism and gastrointestinal gene programs, are only identified by archetypes (**Supplementary Table 3**). The ability to identify adaptive programs in archetype analysis stems from the focus on boundary states that represent specialized cellular functions, rather than average behaviors captured by clustering. The combination of comprehensive sampling across metastatic sites and archetype-based analysis thus provides a powerful framework for discovering key metastatic phenotypes.

### Stomach wall metastases express gastrointestinal gene programs

While liver metastasis in PDAC is well-studied, metastasis to the stomach wall is rare and poorly characterized, despite often leading to severe gastrointestinal complications including pain, ascites, bowel obstruction and other morbidity. Our analysis revealed evidence of organ-specific adaptation: tumor cells from the stomach wall are enriched in archetype cluster 5 (AC5), and the vast majority of AC5 cells originate from this site (**Figs. 3b,c** and **4a**). Hotspot analysis identified three distinct gene modules expressed by AC5 cells that correspond to intestinal, stomach, and gallbladder epithelial cells based on healthy human single-cell reference data^91^ (**Fig. 4b**). These gene modules are minimally expressed in normal pancreatic tissue, indicating that while these metastatic cells are of pancreatic origin, they have acquired transcriptional programs resembling other gastrointestinal epithelia (**Fig. 4c** and **Supplementary Tables 6 and 7**).

**Figure 4.**
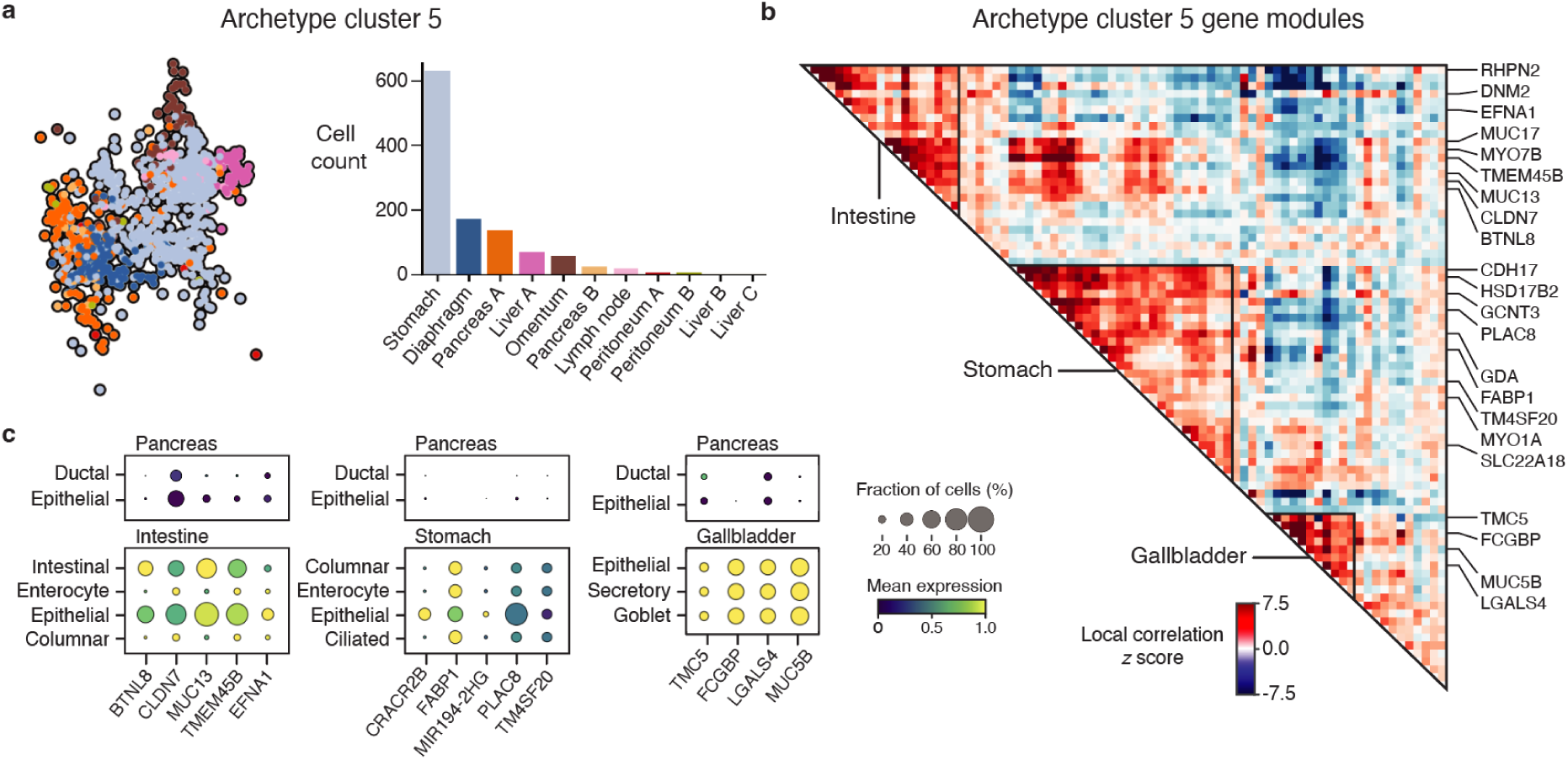
A gastrointestinal archetype indicates PDAC adaptation to the stomach environment. **a**, UMAP embedding (left) and distribution by sample (right) of archetype cluster 5 cells. **b**, Hotspot modules in all archetype cluster 5 cells, based on 78 highly variable genes with significant autocorrelation (FDR < 0.05). Highlighted genes were used to annotate intestine, stomach and gallbladder modules. **c**, Expression of archetype cluster 5 genes in normal pancreas and gastrointestinal tissues based on the CZ CELLxGENE Discover database ^91^.

Archetype cluster 5 genes reflect diverse functions of the gastrointestinal tract, including digestion, nutrient absorption, protective barrier maintenance, and bile production (a gallbladder function), which are distinct from physiological pancreatic capabilities. Although the pancreas is a gastrointestinal tissue, it is considered an accessory organ whose primary function is to secrete digestive enzymes and bicarbonate to neutralize stomach acid. We found AC5 gastrointestinal genes related to cell adhesion and structural integrity (*CDH17, RHPN2, CLDN7, MYO1A, MYO7B*), mucus production and protection (*MUC17*, *MUC13*, *MUC5B*, *FCGBP, GCNT3*), metabolism and transport (*HSD17B2*, *FABP1*, *SLC22A18*, *GDA*), and epithelial cell differentiation (*PLAC8*). Our analysis thus demonstrates that PDAC metastatic cells acquire extensive new gastrointestinal features in the stomach, likely as an adaptation or response to its unique signaling milieu.

Interestingly, a small group of cells from the primary tumor also predominantly express the AC5 gene program. Mapping archetype clusters to the primary tumor revealed that AC5 cells correspond to a classical–mucin phenotype (**Fig. 2e** and **Extended Data Fig. 7a**), which has been observed in human primary PDAC tumors, as well as primary lung, colorectal, gastric^92^, liver, and head and neck cancers^93^. Consistent with this phenotype, AC5 cells in primary tumor express high levels of mucin (*MUC13*, *MUC5AC*, *MUC5B*), mucin production (*GCNT3*, *TFF1–TFF3*)^94,95^, and mucus-producing goblet cell differentiation (*CREB3L1*)^96^ genes, compared to other primary tumor cells (**Extended Data Fig. 7b**). We find that the classical–mucin phenotype is more similar to metastatic states than to other primary phenotypes, as classical–mucin cells co-embed near metastatic AC5 cells and are separated by shorter diffusion distances, reflecting greater transcriptional similarity (**Extended Data Fig. 7c** and Methods).

We examined clonal membership to understand the origins of AC5 classical–mucin cells in the primary tumor, finding that they belong to advanced clones composed mainly of metastatic liver and stomach wall cells, which suggests that the primary tumor was reseeded by stomach wall metastases (**Extended Data Fig. 7d**). These clones appear to express more advanced classical–mucin phenotypes, and not the earlier classical–mucin–HLA-II phenotypes (**Extended Data Fig. 7e**). To confirm the clone assignments of primary cells expressing AC5, we examined the CNA profiles of cells from the two advanced AC5 clones harboring the most primary cells (clones I and J, **Extended Data Fig. 7d**) and found that they are more similar to the profiles of their assigned clones than those of non-metastasizing primary clones (**Extended Data Fig. 7f** and Methods). The primary cells in these clones exhibit similar assignment confidence values as the other cells (primarily stomach and liver) in their assigned clones (**Extended Data Fig. 7f**). Thus, although few primary cells express the AC5 program, they are assigned to the advanced clones with high confidence and provide strong evidence for reseeding.

Another mucus production program, which includes robust expression of transcription factor *SPDEF* and its targets *AGR2* and *ERN2*, is highly expressed in precancerous lesions and classical tumor subtypes^97^. We found that these genes are enriched in primary archetype cluster 14 (AC14), which also expresses high levels of HLA-II molecules, thus fully capturing the PDAC primary classical–mucin–HLA-II phenotype (**Fig. 3b** and **Extended Data Fig. 7b**). Moreover, the classical–mucin–HLA-II cells belong to the earliest clone in the phylogeny (**Fig. 2b,d,e**), supporting that this program is indeed related to early PDAC stages, as reported in mouse models and laser-capture microdissected epithelium from patients with PDAC^97^. In a phase II first-line chemoimmunotherapy clinical trial in advanced gastroesophageal adenocarcinoma patients, a gene program containing AC5 genes *TFF1* and *MUC5AC* was the program most highly expressed by cancer epithelial cells in fast-progressing patients compared to slow progressors^98^. In contrast, expression of AC14 genes (HLA-II programs) by cancer epithelial cells was significantly higher in slow progressors^98^. Our results suggest that PDAC cells can express at least two different mucin production programs—classical–mucin–HLA-II captured by AC14, representing earlier-stage primary cells with a less aggressive prognosis, and classical–mucin associated with AC5, representing later clones associated with greater metastatic potential or chemotherapy resistance.

### Peritoneal metastases rewire lipid metabolism

Archetype cluster 2 (AC2) consists almost entirely of cells from the peritoneum (**Figs. 3b** and **5a**). The peritoneal cavity is the second most common site of metastasis in pancreatic cancer^46,49^, but the mechanisms of metastatic initiation, progression, and adaptation remain poorly understood^99^. Unlike hematogenous metastases to the liver or lungs, which typically present as discrete nodules or masses, peritoneal dissemination often occurs through trans-coelomic spread, leading to thin, diffuse layers over the omentum that escape detection^48,100^. Peritoneal metastases are typically only diagnosed after reaching an advanced, treatment-refractory state known as peritoneal carcinomatosis, which accelerates cachexia—a syndrome characterized by malabsorption, significant weight loss, malignant ascites, and bowel obstruction—and the subsequent rapid decline limits opportunities for investigation.

**Figure 5.**
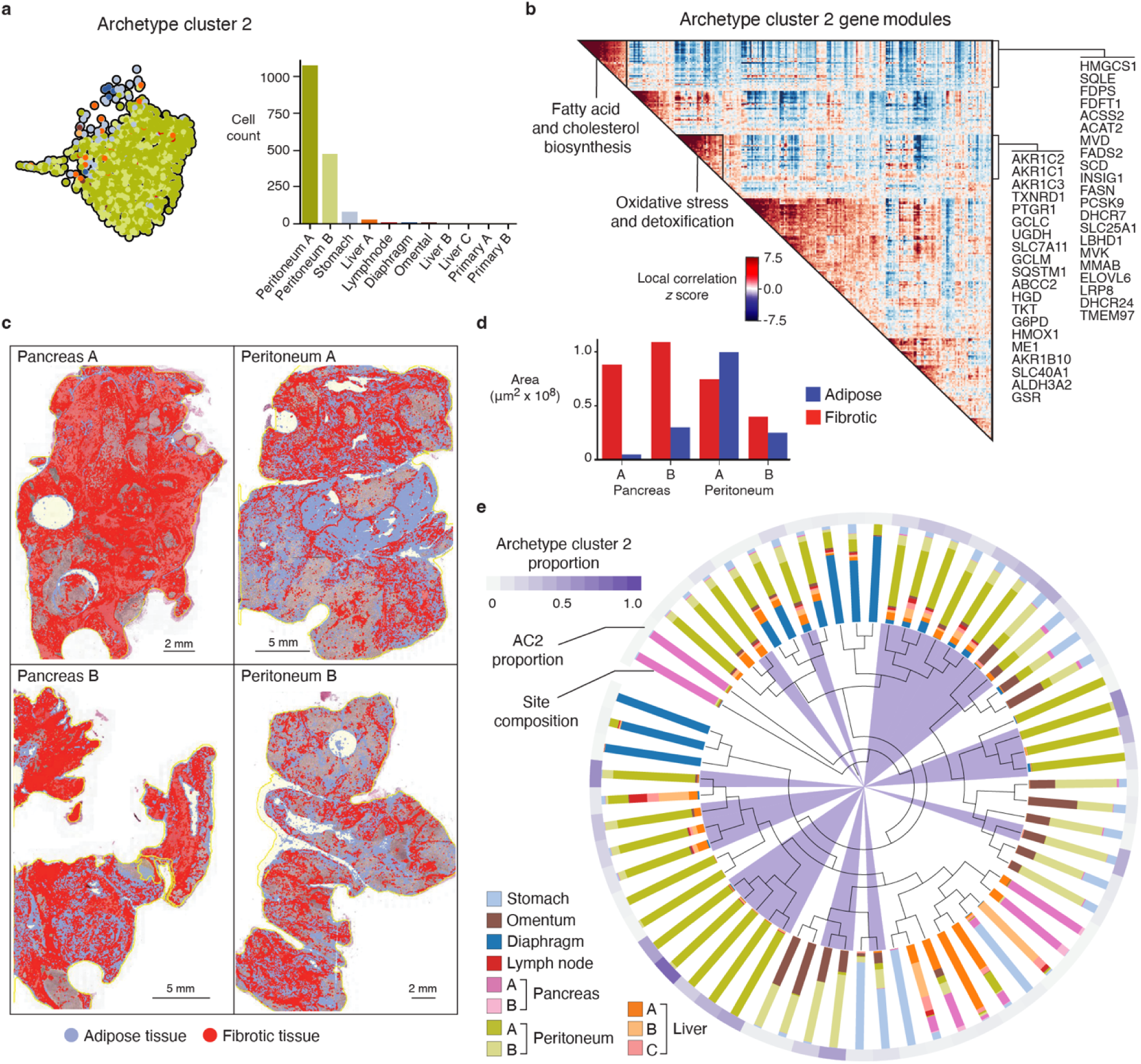
Lipid metabolic rewiring is a prominent feature of peritoneal metastases. **a**, UMAP embedding, colored by tissue site (left), and sample distribution and composition (right), of all archetype 2 cluster cells. **b**, Archetype 2 Hotspot analysis, highlighting lipid metabolism and oxidative stress and detoxification modules. **c**, Digital pathology of H&E-stained primary and peritoneal metastasis tissue, showing expansion of adipose tissue in the peritoneum. **d**, Quantification of adipose and fibrotic tissue in sections in (**c**). **e**, Cancer clone phylogeny, indicating AC2-enriched clones (purple triangles), fractional tumor site composition for each clone (stacked bars) and proportion of cells in each clone assigned to AC2 (outer circle).

We found that the two peritoneal metastases from opposite flanks of the patient both contribute substantially to AC2 (**Fig. 5a**) and have very similar transcriptomic profiles (median 33% of a cell’s kNN graph neighbors are from the other site), including strong upregulation of lipid metabolism genes compared to other archetype clusters (**Extended Data Fig. 8a, Supplementary Table 2** and Methods). Hotspot identified multiple gene modules, including one associated with fatty acid and cholesterol biosynthesis and another with oxidative stress and detoxification (**Fig. 5b**). Genes uniquely upregulated in AC2 include key players in cholesterol (*TM7SF2*) and fatty acid (*ME1, IDH1*) biosynthesis; aldo-ketoreductases (*AKR1B10*, *AKR1C2*, *AKR1C3*); prostaglandin regulators (*PTGIS*, *PTGR1*); and redox balance genes (*GCLM*, *GCLC*, *GPX2*, *GSR*, *PIR*, *SLC7A11*, *TXNRD1, UGDH*) that respond to oxidative stress triggered by lipid production and accumulation (**Fig. 5b** and **Extended Data Fig. 8b**). Genes involved in lipid droplet turnover (*SQSTM1*), lipid transport (*ABCA10, ABCC3*) and adipocyte differentiation (*PLAC8*) are also differentially upregulated in AC2 cells. These genes constitute many components of the lipogenic pathway, by which fatty acids are synthesized for energy storage and cell membrane biosynthesis, primarily in the liver and adipose tissue (**Extended Data Fig. 8c**). Lipid metabolic and oxidative stress genes are not expressed appreciably in tumor immune or stromal cells, confirming that their detection in cancer cells is not due to ambient peritoneal RNA (**Extended Data Fig. 8b**).

The peritoneal cavity is supported by metabolically active adipose tissue that is rich in free fatty acids and signaling molecules, including adipokines and cytokines^101^. Digital pathology of peritoneal and primary tumor sections revealed a greater fraction of adipose tissue in peritoneal metastases than in primary samples, which are dominated by fibrotic stroma (**Fig. 5c,d**). Moreover, whereas cancer cells in primary PDAC tumors typically occur in multiple distinct pockets^102^, they are interspersed among adipocytes in the peritoneal samples (**Extended Data Fig. 8d**). Thus, in contrast to the catabolic processes and patient-level wasting caused by cachexia, our observations suggest that metastatic PDAC cells respond to the adipocyte-rich peritoneal environment by upregulating lipid anabolism and oxidative stress detoxification. A lipogenic phenotype has been reported previously in PDAC cell lines^103^ as well as preclinical models and PDAC patients^104^, but its robust upregulation has not been associated with peritoneal metastasis.

To determine whether the lipid metabolic phenotype generalizes beyond the two independent samples in our patient, we obtained two post-mortem peritoneal metastases from a different patient with PDAC and performed snRNA-seq and similar data analysis (Methods). Importantly, the second subject was 70 years old, succumbed to metastatic disease within three months of diagnosis, and did not receive treatment. Despite the markedly different clinical circumstances in these two cases, we found that fatty acid and cholesterol biosynthesis, as well as cholesterol metabolism and homeostasis, are the most significantly enriched gene programs in the second case (**Extended Data Fig. 9a** and **Supplementary Table 8**).

### Lipid metabolic rewiring is not driven by genotype

We sought to understand whether the highly specialized phenotypes that dominate peritoneal metastases are due to clonal selection of genetically encoded adaptive traits, or were acquired by epigenetically plastic cells in response to a novel environment. To help distinguish between these possibilities, we leveraged the two anatomically separate peritoneal metastases and the cancer phylogeny.

We hypothesized that if clonal selection—under the clonal evolution model^105^—drove the lipid anabolism phenotype, distinct clades of AC2 clones would map to each peritoneal site; after passing through the original selection bottleneck, cells at each site would accumulate unique sets of alterations over time due to genetic drift. On the other hand, if the lipid anabolism phenotype was due to plastic cells responding to the lipid-rich peritoneal environment, there would be no association between clone identity and peritoneal site, and diverse peritoneal clones could contain cells from opposite flanks of the body. We assessed which lipid anabolism-enriched clones (defined as >10% of cells with AC2 phenotype) belong to each peritoneal metastasis in the phylogenetic tree, and found 26 clones spread across all three major clades, including early branches with fewer CNAs as well as late branches (**Fig. 5e** and **Extended Data Fig. 9b**). Both pure and mixed clones are present in the independent peritoneal sites. For example, early clones enriched for lipid metabolism are derived from both peritoneum A (16% to 68%) and B (8% to 40%) sites, and late clones are derived from a mix of sites as well (69% to 83% peritoneum A and 8% to 12% peritoneum B). Intermediate clones are pure for either peritoneum site but still share the same clades, suggesting common ancestors in the primary tumor (**Extended Data Fig. 9b**).

The existence of diverse clones enriched for lipid anabolism over several branches of the tree—some populating both peritoneal sites, some specific to each site but belonging to the same clade—support the hypothesis that cancer cell plasticity drove the lipid anabolism phenotype through phenotypic convergence to local environmental.

### Transcriptomic plasticity is a hallmark of PDAC metastasis

The plasticity we identified in peritoneal tumors involves multiple clones that express a diversity of additional archetypes, motivating a more systematic investigation of whether plasticity is a general feature of PDAC metastasis. Indeed, while each clone represents a shared genetic lineage, clones throughout the phylogeny do not appear to be constrained by their lineage and express a diversity of archetypal phenotypes, corresponding to high per-clone archetype entropy (**Fig. 6a** and Methods). This is in line with a lineage tracing study in a PDAC mouse model, which found that cell cycle and EMT cell states are not correlated with cellular phylogeny^106^. We asked whether the diversity of archetypes that clones exhibit is greater than expected, which would indicate substantial phenotypic plasticity (Methods). The empirical distribution of per-clone archetype entropy (mean *μ* = 1.42) is shifted higher than expected under simulations in which the site is highly predictive of phenotype (*μ* = 0.97), but lower than expected under random assignment of archetypes to cells (*μ* = 2.42, **Fig. 6b** and Methods). This suggests that the variety of archetypes present in each clone is not driven by the diversity of tumor sites within each clone, but rather by the ability of cells to acquire a range of phenotypes even within a single site.

**Figure 6.**
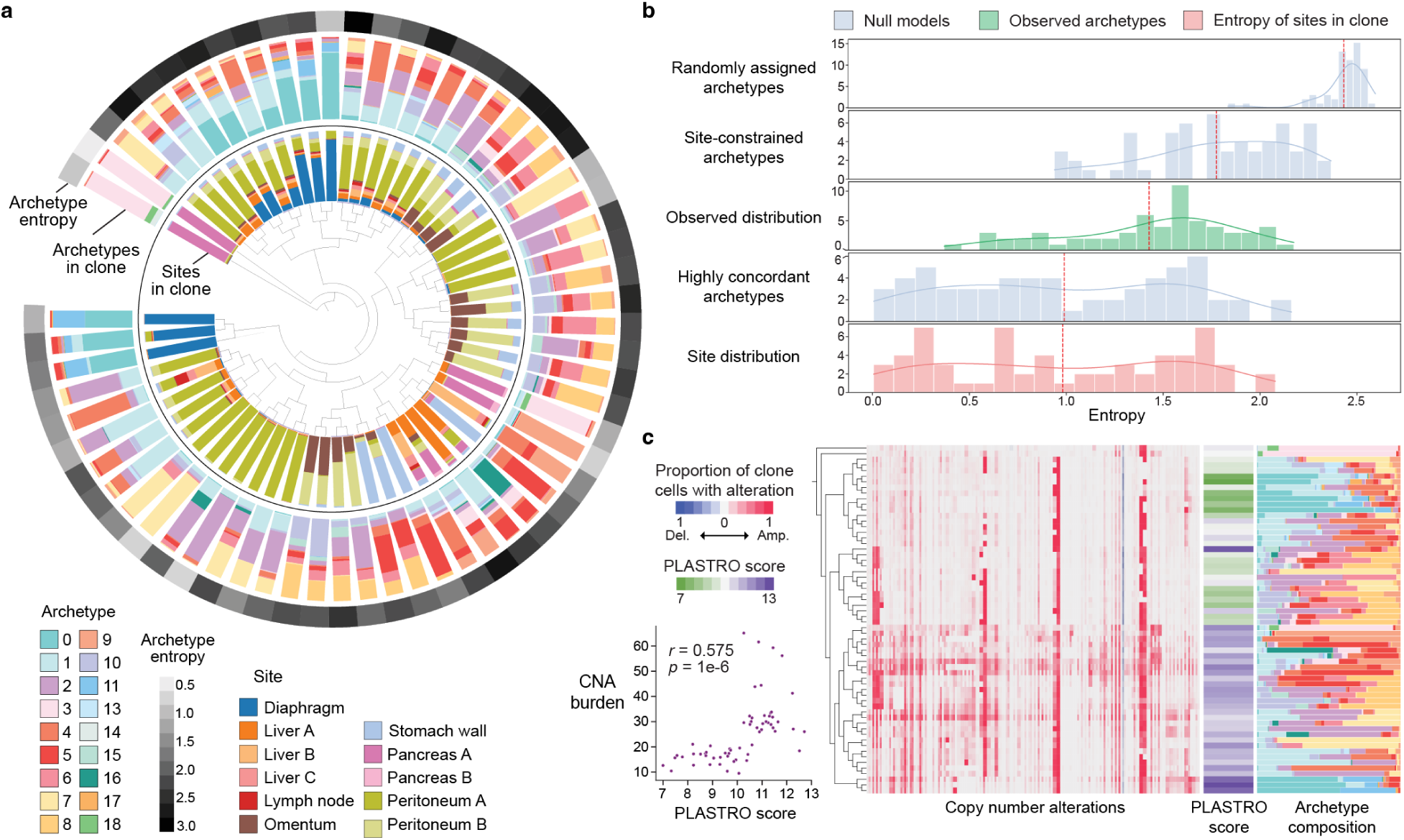
Transcriptomic plasticity is a common feature of metastatic cells. **a**, PDAC tumor cell clonal phylogeny (center), showing fraction of cells from each site, fraction of archetypes, and archetype entropy from inside to outside, for each clone (leaf in the phylogeny). **b**, Entropy distributions for three null models and for data in this study (Methods). Bars indicate the number of clones for each binned entropy value (*n* = 62 clones for each distribution), curves represent smooth trends, and dashed vertical lines correspond to mean entropy. Observed clones have lower archetype entropy than clones with randomly assigned archetypes, but more than models based on strong archetype bias for metastatic site, indicating high cellular plasticity. **c**, PDAC tumor cell phylogeny showing copy number profiles of each clone, PLASTRO score (Methods) and archetype composition. Scatterplot indicates that higher CNA burden is associated with higher plasticity. *r*, Pearson correlation.

To quantify plasticity more rigorously at the clonal level, we developed plasticity analysis from single-cell transcriptional and evolutionary neighborhood overlap (PLASTRO) (**Extended Data Fig. 10** and Methods). PLASTRO compares evolutionary similarity (lineage distance) and phenotypic similarity (phenotypic distance with respect to archetype composition) between cells, based on the assumption that low cellular plasticity should result in a strong overlap between lineage and phenotype. Specifically, for a given clone, it quantifies the degree of discordance between phenotypic and phylogenetic neighborhoods, while remaining insensitive to neighborhood size. Interestingly, we found that cells with few CNAs tend to have low PLASTRO scores, whereas more advanced clones bearing extensive CNAs score high for plasticity (**Fig. 6c**). Given that CNA burden correlates with metastasis in PDAC and other cancers^8^, our finding that CNA burden is strongly associated with plasticity is consistent with a model whereby plasticity enables metastasis. To evaluate this effect more quantitatively, we performed a Mantel test^107^, which assesses the correlation between two distance matrices (Methods). We observed a Mantel test statistic of 0.13 (*p* < 1 ×10^−3^) for matrices of phenotypic distances within distinct clones, suggesting that cells are plastic (the statistic ranges between –1 and 1, with 0 denoting no correlation), as their phenotypes differ significantly from that of their lineage.

## Discussion

Rapid autopsy makes it possible to investigate clonal lineage histories and adaptive phenotypes in a single cancer ecosystem. In our comprehensive analysis of a patient with PDAC who underwent rapid autopsy, we evaluated the phenotypic landscape that a single cancer can occupy and developed computational approaches that bridge single-cell transcriptomics with phylogenetic reconstruction to dissect the relative contributions of clonal evolution and transcriptomic plasticity to metastatic adaptation. Our analysis reveals that transcriptional plasticity, rather than genetic evolution and selection, is the dominant force shaping metastatic phenotypes.

A critical insight from our study is that successful metastatic clones exhibit remarkable phenotypic diversity, even within the same anatomical site. We demonstrate that genetically related metastatic clones can colonize multiple organs while manifesting diverse transcriptional states independent of their anatomical location. Moreover, each organ site harbored multiple phylogenetically distant clones, suggesting extensive parallel seeding. This evidence points to non-genetic plasticity as a key mechanism enabling metastatic cells to transition between different gene programs across metastatic sites, thereby enhancing their adaptability and survival. This plasticity is notably amplified in clones with higher CNA burden, suggesting that genomic instability may facilitate transcriptional adaptation—not through specific mutations, but by creating a permissive state for phenotypic exploration. This observation aligns with recent findings that chromatin accessibility increases with genomic instability in various cancers, potentially enabling broader transcriptional responses to environmental cues^108,109^.

Our profiling of common but understudied metastatic sites in PDAC revealed distinct organ-specific adaptation programs, providing new insight into how cancer cells respond to diverse tissue environments. The acquisition of gastrointestinal programs by stomach wall metastases demonstrates remarkable cellular plasticity, suggesting that tumor cells can co-opt organ-specific transcriptional modules to enhance colonization and acquire fitness in new environments. Similarly, peritoneal metastases upregulate lipid anabolism and oxidative stress response pathways, suggesting that tumor cells adopt metabolic features of adipocytes and adapt their redox response to counteract reactive oxygen species generated by metabolic stress. This is consistent with prior studies showing that lipid metabolism plays a crucial role in PDAC progression^110^ and chemoresistance^111^. Both site-specific gene programs suggest that metastatic cells adapt to their microenvironment, possibly in response to stroma-derived signaling and environment lipid availability^112^. The convergent adaptation of these phenotypes across multiple independent clones strongly supports the role of microenvironmental pressures in shaping cellular phenotypes, independent of genetic evolution.

The methodological advances developed for this study—particularly PICASSO for phylogenetic reconstruction and our approach to archetype analysis—provide a robust framework for similar investigations across cancer types. However, several critical questions emerge from our findings: How do specific tissue environments orchestrate the activation of adaptive programs? What role do stromal cells play in this process? Can we target the mechanisms underlying cellular plasticity with therapies? What do genomic markers such as *RAS* amplification contribute, given the importance of plasticity as an emerging resistance mechanism to *RAS* therapies? Future studies combining spatial transcriptomics with single-cell lineage tracing could help address these questions and further illuminate the complex interplay between genetic inheritance and environmental adaptation in cancer progression.

Our findings emphasize the fundamental roles of cellular plasticity and metabolic adaptation in enabling the successful colonization of diverse organ sites. They suggest that effective therapeutic strategies must account for both genetic and non-genetic mechanisms of adaptation, potentially through approaches that constrain cancer cell plasticity or target site-specific vulnerabilities. These insights may guide the development of more effective treatments for metastatic disease, particularly for challenging sites such as peritoneal metastases that currently lack targeted therapeutic options.

## Supplementary Tables

**Supplementary Table 1** | Marker genes used for cell-type annotation.

**Supplementary Table 2** | Upregulated DEGs of archetype clusters (log-fold change > 1, *p* < 0.05) assessed for intracluster modularity using Hotspot, and informative genes (autocorrelation, FDR < 0.05).

**Supplementary Table 3** | GSEA scores of significantly enriched (FDR < 0.05) archetype cluster genes.

**Supplementary Table 4** | Full GSEA enrichment scores of archetype cluster genes.

**Supplementary Table 5** | GSEA scores of significantly enriched (FDR < 0.05) Leiden cluster genes.

**Supplementary Table 6** | Gene expression of archetype cluster 5 gastrointestinal genes for intestinal, stomach, and gallbladder epithelial cell types in CELLxGENE.

**Supplementary Table 7** | Gene fraction of cells expressing archetype cluster 5 gastrointestinal genes for intestinal, stomach, and gallbladder epithelial cell types in CELLxGENE.

**Supplementary Table 8** | GSEA scores of significantly enriched (FDR < 0.05) archetype neighborhood cells in peritoneal metastases of rapid autopsy RA19_21.

**Supplementary Table 9** | Expert-curated gene sets used in this study.

## Acknowledgements

In deep gratitude, we honor the patient and family, whose courage and contribution to medical research for pancreatic cancer continue to inspire progress and hope for future patients. This work was supported by National Institutes of Health (NIH) cancer center support grant P30 CA08748 and Human Tumor Atlas Network grant U2C CA233284 (D.P., C.I.D., I.P.), as well as U54 CA274492 (D.P), R35 CA220508-03 (C.I.D.), P50 CA25788 (C.I.D., D.P.), and the AACR-CRUK Transatlantic Fellowship (A.J.S). D.P. is a Howard Hughes Medical Institute investigator. We thank Vincent Liu for initial data exploration, and the Last Wish Program, Gerry Metastasis and Tumor Ecosystems Center, David M. Rubenstein Center for Pancreatic Cancer Research and Single-cell Analytics Innovation Lab at MSK for their support.

## Contributions

D.P. and C.I.D. conceived the study and designed and supervised experiments. W.P. and E.M.O. provided patient care, clinical insights and diagnostics. A.H. collected and processed biospecimens. A.H., S.U. and J.H. provided pathological analysis and contributed to data interpretation. L.M. and R.C. oversaw single-cell genomics and I.M. carried out single-cell protocol optimization and data generation. I.P. and D.P. conceived and supervised algorithm development, Y.X. and A.J.S. performed data processing, S.P., T.C., Y.X. and A.J.S. developed IntegrateCNV, S.P. developed PICASSO and plasticity analysis approaches, A.J.S. and R.S. developed the archetype analysis pipeline. A.J.S., S.P., R.S., A.M., T.N. and D.P carried out data analysis and interpretation, and D.P., A.J.S., S.P. and T.N. wrote the manuscript. All authors reviewed and edited the manuscript.

## Competing interests

A.M. has served in consultant or advisory roles for Third Rock Ventures, Asher Biotherapeutics, Abata Therapeutics, Clasp Therapeutics, Flare Therapeutics, venBio Partners, BioNTech, Rheos Medicines and Checkmate Pharmaceuticals. He was previously an Entrepreneur-in-Residence at Third Rock Ventures, is currently a Venture Partner for The Column Group, is a co-founder of Monet Lab, an equity holder in Monet Lab, Clasp Therapeutics, Asher Biotherapeutics and Abata Therapeutics, and has received research funding support from Bristol-Myers Squibb (BMS). C.I.D. received research support from BMS and is on the advisory board of Episteme. D.P. has equity interests and consults for Insitro. All other authors declare no competing interests.

## Methods

### Biospecimen collection

#### Patient information

Warm autopsy samples were collected from a 35-year-old female patient with informed consent to the Last Wish Program and approval by the patient’s family and the Internal Review Board of Memorial Sloan Kettering Cancer Center (IRB protocol 15-021). The patient was diagnosed with metastatic PDAC, exhibiting macroscopic lesions in the pancreas and liver (detected by computed tomography scan) and upregulated CA19-9 tumor biomarker. The patient was treated with standard mFOLFIRINOX therapy and tumors showed clinical response for approximately 6 months before they stopped responding, at which point mFOLFIRINOX was halted and a dose of Gemcitabine + nab-Paclitaxel was given, but no further response was observed. The patient survived for just over 9 months from diagnosis, which is expected in a metastatic PDAC patient treated with standard chemotherapy.

Both primary and metastatic tumors were readily detectable. The primary tumor appeared as a white-gray mass, while liver metastases were white-yellow with extensive necrosis. Multiple peritoneal and omental metastases, along with a single gastric metastasis, were palpable and appeared as white nodules. Prominent diaphragm metastases resembling an “omental cake” were also identified.

#### Biospecimen collection

Samples were obtained using standard autopsy techniques, specifically the Rokitansky method. Following the removal of all organs from the body, more than 50 samples were collected from macroscopically identifiable tumors in both primary and metastatic sites. Autopsies were initiated within two hours of death, and biospecimens were collected within an hour. Multiple lesions collected from the same organ were clearly separate anatomically. The exception is primary tumor, for which two adjacent sections were processed as Pancreas A and B samples (see below for sectioning information) for single-nucleus RNA sequencing. Tumors larger than 1 cm in size were trimmed to 1-cm squares, then divided in half. One half was used to generate a formalin-fixed paraffin-embedded (FFPE) block for detailed histological analysis. The other half was cut into 5–7 mm pieces, placed in cryotubes, and rapidly frozen in liquid nitrogen before storing at −80°C. For particularly large primary tumors, samples were obtained after slicing. The position of each sampling site within the organ was meticulously documented during the autopsy. Approximately 10 samples were taken from normal tissue alongside the tumors.

For whole exome sequencing (WES), a portion of each flash-frozen sample was used to create an optimal cutting temperature (OCT) block. H&E staining of frozen OCT sections was performed to identify tumor regions and confirm inclusion of sufficient tumor tissue before macrodissection to extract DNA for bulk WES, typically from 5–10 sections.

For snRNA-seq, a different portion of the frozen tissue was sectioned and tumor tissue inclusion was confirmed using the frozen H&E slide method before proceeding with single-nucleus suspension and sequencing library preparation.

### Experimental Methods

#### Whole exome sequencing

Bulk whole exome sequencing was performed as described in refs.^18,113^. Briefly, genomic DNA was extracted from each tissue sample using QIAamp DNA Mini Kits (Qiagen). Sequencing was carried out on an Illumina HiSeq 4000 or NovaSeq 6000 platform, with a target coverage of 250x for all samples.

#### Single-nucleus RNA-seq Generation of nucleus suspensions

Single-nucleus suspensions were generated following the Frozen tissue dissociation for single-nucleus RNA-seq protocol^114^. Specifically, frozen rapid autopsy specimens were cut into approx. 2 mm^3^ pieces using a disposable scalpel (Technocut, 10148-882) and transferred to 1 ml of freshly prepared ice-cold lysis solution (250 mM sucrose, 50 mM citric acid, 0.01% DEPC). Next, the entire lysis solution with specimens was transferred to a Dounce homogenizer (Sigma, D8938-1SET). The tissue pieces were grounded by gently moving a large-clearance pestle (Tube A) up and down 10 to 15 times, followed by gentle up-and-down movement 10 times using a small clearance pestle (Tube B). After grinding steps, the homogeneous suspension of minced tissue was strained through the 35-µm snap cap strainer (Fisher Scientific, 352235) and kept on ice for 1 min. Filtered nucleus suspension was transferred into a 2-ml tube and spun down at 4 °C in a swinging bucket centrifuge at 500 *g* for 5 min. The supernatant was discarded, leaving ∼20 µl above the nucleus pellet. Next, the pellet was resuspended in 1 ml ice-cold 1 ml nucleus wash buffer (250 mM sucrose, 50 mM citric acid, 1% (w/v) BSA, 20 mM DTT and 0.2 U µl^-1^ RNase inhibitor (Ambion, AM2682), in DEPC-treated water (Ambion, AM9915G)). The tube was spun down in a swinging bucket centrifuge at 500 *g* for 5 min at 4 °C and the supernatant was aspirated without disrupting the now-smaller pellet. The pellet was then resuspended in 0.5 ml nucleus resuspension buffer (3X SCC (Invitrogen, AM9770), 20 mM DTT, 1% (w/v) BSA, and 0.2 U µl^-1^ RNase inhibitor (Ambion, AM2682), in DEPC-treated water (Ambion, AM9915G)) and passed through a 35-µm snap cap strainer. Nuclei were quantified by staining 10 µl of nucleus suspension with 0.2 µl of 100 μg ml^-1^ DAPI and 10 µl of 0.4% Trypan Blue, and carefully inspected for quality and separation under bright field and fluorescence microscopes. The entire procedure took approximately 1 hr to complete and generated 10^6^ –10^7^ single nuclei per 1 ml.

#### Single-nucleus enrichment

Prior to snRNA-seq, single nuclei were purified by fluorescence-activated cell sorting (FACS) to remove debris and clumps following ref.^114^. In a typical scenario, a 50-µl aliquot of the nucleus suspension was added to 250 µl nucleus resuspension buffer and used as an unstained reference sample for FACS, and the remaining suspension (∼900–950 µl) was stained with 10 µl of 100 μg ml^-1^ DAPI. Nucleus sorting was performed on a BD FACS Aria II instrument equipped with a 100-µm nozzle. Sorting was conducted at 5,000–10,000 events/second, by selecting events based on DAPI signal and particle size. The sorted nuclei were transferred to 1.5-ml Protein LoBind tube (Eppendorf) and centrifuged in a swinging bucket at 600 *g* for 5 min at 4 °C. The nucleus pellet was resuspended in 100 µl of supernatant and manually counted under bright field microscope after mixing 10 µl of nucleus suspension with 10 µl of 0.4% Trypan Blue. The suspension concentration was adjusted to obtain ∼2000 nuclei/µl before proceeding with the v3 chemistry kit on the Chromium instrument (10x Genomics).

All samples were split and processed by the sorting protocol above or without it (unsorted). Both unsorted and sorted samples were submitted for snRNA-seq preparation to ensure no systematic biases were experimentally generated.

#### snRNA-seq library preparation

Single-nucleus RNA library preparation was performed following the Chromium Single Cell 3’ Reagent Kits User Guide, v3.1 Chemistry (10x Genomics), as in ref.^114^. Library sequencing was performed on Illumina NovaSeq 6000 instruments using a paired-end 2 x 150-bp configuration.

### Algorithmic development

#### IntegrateCNV for copy number inference

Copy number inference from scRNA-seq data assumes that changes in gene expression reflect underlying changes in gene dosage. However, epigenetic factors also affect expression and obscure the link between expression and copy number. Furthermore, scRNA-seq data is noisy and sparse, leading to noise in the inferred copy number profiles. To mitigate noise and sparsity, we restrict single-cell copy number inference to regions known with high confidence to harbor copy number alterations (CNAs) based on bulk WES data, thereby greatly reducing false positive CNAs. Sparsity is also mitigated by aggregating expression across genes for greater robustness within these regions.

We developed IntegrateCNV to infer per-cell copy number variation from paired single-cell/nucleus RNA sequencing and bulk WES data (**Extended Data Fig. 3b**). IntegrateCNV first identifies regions likely to harbor CNAs in WES data, then calculates the likelihood of each genomic region being affected by these alterations in each single-cell. IntegrateCNV accepts as input (i) a cell × gene matrix of scRNA-seq count data and “normal” or “tumor” annotation for each cell, and (ii) paired copy number profiles from bulk WES data in matching tissue. Using this information, the algorithm determines (i) a set of chromosomal regions that are copy-neutral across all samples, and (ii) a set of chromosomal regions of sufficient size that are altered in at least one sample. Finally, integrateCNV outputs (iii) a cell × region matrix containing the likelihood of that cell being copy number neutral in that region for each (cell, region) pair.

### IntegrateCNV algorithm

The integrateCNV algorithm performs a two-tailed hypothesis test to determine whether each (cell, region) pair has expression levels that differ significantly from the expression levels in known normal cells. The null distribution of expression in each region is Gaussian, with expression mean and variance taken from matching regions in a set of reference normal cells. The algorithm performs the following steps:

1. Identify chromosomal regions that are copy number neutral across all samples as a normalization factor.
2. Identify chromosomal regions that are copy number altered in at least one sample based on bulk WES data.
3. Aggregate expression across genes within each altered region.
4. Normalize and log-transform the per-region expression.
5. Determine the null distribution based on annotated non-tumor cells.
6. Perform a hypothesis test to indicate the presence or absence of an alteration.

IntegrateCNV allows us to better normalize single-cell expression data against neutral regions without removing the biological signal inherent in library size.

#### Determining neutral and altered regions

The first input to integrateCNV is a set of copy number profiles derived from bulk DNA sequencing. For each sample, we use FACETS^115^ to identify the total copy number in each region. CNAs are centered around 0 so that a neutral region is represented by the copy number ‘0’. The CNA profiles are saved as BED files, containing, for each region, information about the chromosome, start position, end position, and copy number. BED files from all samples are processed to find intersecting genomic regions using the multi_intersect function from pybedtools. The resulting intersections capture the chromosomal regions and CNAs in each sample. Neutral regions are then identified as those with no CNA in any sample. We denote the set of neutral regions by A^0^.

Candidate altered regions are first identified as those in which at least one sample contains an alteration. Of the candidate regions, only those containing sufficient genes (>20 genes by default) are retained for downstream analysis so as to provide sufficient coverage to reliably recover copy numbers without being unduly influenced by the potential outlier effects of few genes. This set of altered regions, A^20+^, is used as the set of regions within which we will infer copy number alterations.

#### Processing count data

We denote the scRNA-seq cell × gene count matrix by *X*, where *X*_*c*,*g*_ represents the expression of gene *g* for cell *c* = 1, ···, *n*. The raw count matrix, *X*, is the second input to the integrateCNV algorithm. Using the set of candidate regions (A^20+^), we aggregate counts over genes within a given region, indexed by *r*, to determine a cell × region matrix, *U*.

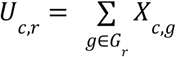

where *g* ∈ *G*_*r*_ are the genes which physically overlap with the genomic region indexed by *r*.

The counts from regions A^0^ that are found to be neutral in all samples are used as a pseudo ‘spike-in’ control in order to normalize count data without removing the biological signal of total library size, which can correlate with copy number burden. The total counts from genes across all neutral regions are summed for each cell, *c*, and the sum is denoted by library size normalization factor, *l*_*c*_.

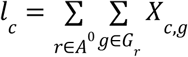

The cell × region matrix, *U*, is then divided by the library size normalization factor and the log of the resulting normalized expression is computed to give data matrix, *V*.

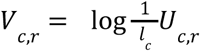

#### Inferring copy number alterations

The log-normalized expression matrix, restricted to normal cells, now defines a null Gaussian distribution on expression levels in unaltered cells for each region. For each cell and region, the *z*-score is computed using this null distribution, and is used to define copy number altered regions.

#### Extracting integer copy number calls

Finally, a two-tailed hypothesis test is performed for each (cell, region) pair to determine whether the cell has expression values significantly higher or significantly lower than expected in a diploid cell. A *p*-value threshold (default 0.05) is used to determine the critical values for two-tailed hypothesis testing. All regions above or below the upper or lower critical values, respectively, are called as alterations. We note that because deleted regions have a small dynamic range (0, 1 or 2), there is less power to detect them and thus the procedure results in many false negatives for deleted regions. For all called alterations, we use the copy number from the corresponding bulk WES sample to insert an integer copy number. This denoising procedure ensures that, for a region to be denoted as altered in a cell, it must be supported by evidence from both snRNA-seq and sample-level bulk DNA data. The final output matrix is an integer copy number profile for each single cell, and can be used for downstream phylogenetic analysis of clonal relationships.

#### Comparison of CNA inference methods

To benchmark integrateCNV against existing approaches that infer CNAs from scRNA-seq data, we first determine single-cell copy number *z*-score profiles, which are computed without any prior knowledge of which sample (or bulk WES data) each cell belongs to. We then aggregate cells within samples to compare against the ‘ground truth’ bulk WES copy number profile.

For each sample, we run inferCNV^22^ and CopyKat^32^, which return per-region and per-gene CNA scores, respectively, for each cell in the sample. For integrateCNV, we compute the *z* score for each altered region harboring an alteration in each cell. We then compute the average score across all cells in the sample to determine a pseudo-bulk CNA score for each method. Since all methods return continuous valued predictions of alterations rather than discrete copy number calls, we compute the correlation between the bulk DNA CNA call and the pseudo-bulked inferred CNA score.

#### Identifying recurrent CNAs

We use the four gamete test^116^ to identify potential violations of the infinite sites assumption that may be due to recurrent alterations. The four gamete test considers mutation states at pairs of sites. We binarize copy number alterations and represent diploid sites as 0, and aneuploid sites as 1. For any two sites in a sequence, there are four possible combinations of mutation states - (1,1), (1,0), (0,1) and (0,0). If all four combinations are observed in a population, this violates the infinite sites model (which assumes each mutation only occurs once).

For each pair of regions for which single-cell copy number profiles were computed by IntegrateCNV, we identify all pairs of mutation states which are observed in our inferred CNA profiles. To account for noise in the copy number inference, we consider only pairs which are represented in at least 100 cells. If all four mutation state pairs are observed, we denote that region pair as violating the infinite sites assumption, potentially due to recurrent copy number alterations.

### Phylogenetic inference from single-cell CNA calls

Most efforts to reconstruct tumor phylogenies rely on single-nucleotide variants (SNVs) derived from DNA sequencing data^117–119^. A few approaches specifically address CNA phylogenies^44,45^, but they are designed for copy number profiles derived from deconvolved bulk DNA sequencing or single-cell DNA sequencing. These methods typically assume that input copy number profiles are reliable and accurately specified for contiguous genomic regions, and most do not scale to large numbers of cells. These assumptions do not hold when considering CNA profiles derived from scRNA-seq experiments, as inferred copy number profiles are very noisy and dataset sizes are significantly larger. Researchers thus often resort to distance-based agglomerative clustering methods such as neighbor joining^120^ to reconstruct cell hierarchies.

To overcome these challenges, we developed PICASSO (phylogenetic inference from copy number alterations in single-cell sequencing observations), to infer cellular clones and their phylogenetic relationships from CNA calls derived from single-cell expression data. The PICASSO algorithm assumes that observed single-cell copy number profiles are noisy measurements of true clonal profiles, such that cells in the same clone share similar CNA patterns. Phylogenetic relationships are unobserved and result from (potentially recurrent) gain and loss of copy number variants from an original parent clone. PICASSO thus aims to group single cells based on membership to inferred clones, and determine the evolutionary relationships between these clones.

As input, PICASSO accepts a character matrix of cells by regions, with each entry consisting of an integer CNA state for the corresponding region and cell. Using this information, the algorithm generates (i) assignments of cells to clones and (ii) a phylogeny describing the relationship between clones.

#### PICASSO algorithm

PICASSO is a tree-recursive algorithm whereby each iteration considers the cells currently assigned to a leaf node of the phylogenetic tree and determines whether to split that leaf into further branches. It comprises the following steps (**Extended Data Fig. 3a**):

1. Encode integer copy numbers into ternary profiles. If the maximum absolute copy number (relative to diploid) is *j*, copy number *k* is encoded as a vector of length *j* with *k* leading 1s so that similar copy number profiles are similar in the encoded space. In practice, we cap the maximum copy number at *j* = 2, distinguishing only between amplified and highly amplified copy numbers. Similarly, negative copy number − *k* is encoded as a vector of length *j* with *k* leading −1s. This allows us to represent the cumulative nature of CNAs, whereby moderate gains or losses may precede more severe alterations, and also account for small mistakes when inferring CNA magnitude.
2. Construct an initial phylogeny comprising a single leaf node containing all cells in the dataset.
3. For each leaf node, split the node into two clones based on shared CNAs using expectation–maximization (EM). Cells are partitioned such that (i) CNAs are allowed to recur independently in distinct clones, and (ii) cells are grouped based on global CNA profile, mitigating the outsize effect of noisy or incorrect calls in a few genomic regions. More explicitly, for each non-terminal leaf in the phylogeny:
  a. If sufficient evidence exists to split cells, assign cells to one of two subclones using EM. These subclones are the new children of the original leaf node.
  b. If insufficient evidence exists to split cells, designate this leaf as a terminal node.
  c. Repeat until all leaf nodes are terminal nodes.
4. Cell groupings identified from this iterative process constitute clone assignments, and relationships between groups constitute the phylogenetic relationships between clones. The tree is re-rooted so that the clone with fewest CNAs is most ancestral, reflecting the fact that CNA burden generally increases during evolutionary progression.
5. As optional post-processing, we may collapse small subclones containing too few cells to draw meaningful statistical conclusions.

#### Encoding the character matrix

We denote the cell × region matrix of integer CNAs by *B*, where *B*_*c*,*r*_ represents the inferred copy number of cell *c* = 1 ··· *n* in region *r* in *A*^20+^, and *A*^20+^ denotes the set of altered regions. To facilitate further analysis, we transform matrix *B* into a matrix *M* using the following encoding scheme:

1. Determine the maximum absolute value. For each column (region) *r* in *B*, determine the maximum absolute value, *P*_*r*_, representing the highest CNA observed in that region. In practice, we cap this value at copy number +2 (two copies more than expected in a diploid cell), as we may not trust or be able to reliably distinguish between very large copy numbers.
2. Encode copy number values. For each cell *c* and region *r*, encode the copy number *k* = *B*_*c*,*r*_ into *P*_*r*_ columns in *M* according to this scheme:
  a. If *k* ≥ 0, the encoding is [1, 1,…, 1, 0, 0,…, 0] with *k* ones followed by *P*_*r*_ − *k* zeros.
  b. If *k* < 0, the encoding is [–1, –1,…, –1, 0, 0,…, 0] with |*k*| negative ones followed by *P*_*r*_ − |*k*| zeros.
3. Construct the matrix *M*. Replace each column *r* in *B* with *P*_*r*_ columns in *M* according to the above encoding scheme, resulting in a ternary matrix where each original region is expanded into multiple columns representing CNA magnitude and direction.

This transformation allows us to enforce similar copy number profiles between CNAs of similar values. The dimension of the resulting matrix *M* is 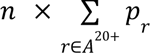.

#### Top-down phylogeny construction

We use an expectation–maximization approach to construct a top-down phylogenetic tree based on shared patterns of copy number breakpoints. The phylogeny is initialized with a single clone containing all the cells in the data set. At each iteration, the depth of the existing tree may be increased by one as each leaf clone may be split into two further subclones if there is sufficient evidence of differences between them. Sufficient evidence of differences between potential subclones exists when the copy number patterns observed cannot be reasonably explained by a single population. Using the Bayesian Information Criterion (BIC), we only create a new branch in the evolutionary tree when the data strongly suggests that two distinct copy number clone populations exist. Alternatively, any given clone may remain intact as a terminal clone.

#### Mixture model for clustering CNA clones

The input to PICASSO is the copy number profile of distinct genomic regions that are likely to harbor CNAs. We therefore assume that CNA occurrences at each genomic region are independent, which allows us to consider each profile as a draw from a multivariate categorical mixture model. We can use an EM algorithm to cluster each existing leaf into two subclones, mimicking the evolutionary process that distinguishes clones by the accumulation of copy number differences.

For each subclone, we learn a probabilistic profile over CNAs, allowing us to capture several essential features. The learned probability associated with the categorical distribution for a given CNA can be less than 1, permitting CNAs to only be present in a subset of cells in an inferred subclone. Further, the subclonal structure can be disentangled by subclone splitting in subsequent iterations. Additionally, the probabilistic profile allows us to model the high degree of false positives and false negatives in inferred CNA data by tolerating small probabilities of a clone missing or containing a specific alteration. Finally, there may be a positive probability of a particular alteration occurring at the same position in both clones, allowing for the independent recurrence of copy number changes in multiple clonal lineages, which has been observed extensively in previous CNA of cancer data^121^.

The EM algorithm is a widely used iterative method to find maximum likelihood estimates of parameters in probabilistic models, particularly for clustering problems. PICASSO uses an EM algorithm for clustering categorical data with states {–1, 0, 1}, which represent different copy number alterations in cells.

The observed copy number profiles, *M* = {*M*_1_, *M*_2_,…, *M*_n_}, contain the encoded CNAs for each cell, *c* = 1 ··· *n*, across regions. Each *M*_*c*_= [*m*_*c*1_, *m*_*c*2_,…, *m*_*cd*_] is a vector of *d* categorical observations for cell *c*. Each observation *m*_*cj*_ can take one of three states: –1, 0, or 1, representing different CNAs.

We assume there are two clusters representing an evolutionary split between subclones, and each cluster *k* is characterized by a set of parameters θ_*k*_ = {π_*k*’_, ϕ_*k*_}, where π_*k*_ parametrizes the prior probability of cluster observation in cluster *k*. θ_*k*_ and ϕ_*k*_ the probability distribution over the states for each observation in cluster *k*.

#### Expectation maximization algorithm

The EM algorithm iterates between the expectation (E) and maximization (M) steps until convergence. The goal is to assign each cell to one of the subclones in a way that maximizes the likelihood of the observed data.

We let ϕ*_k_* ∈ ℝ^3×*d*^ represent the parameters of the categorical distribution for component *k* ∈ {1, 2}, and π*_k_* represent the mixture proportions. We also define the latent variable *z_c_*, which indicates the membership of the *c*-th observation to one of the two components, where *z_c_* ∈ {1, 2}. The responsibility γ*_ck_* = 𝔼[*z_ck_*] is the expectation of *z_ik_*.

The complete data log-likelihood is:

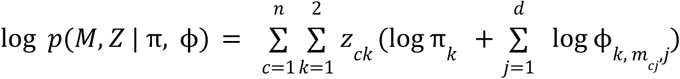

The E-step updates the prediction of which subclone each cell belongs to based on the likelihood of the observed data under the current model. We calculate the posterior probabilities, γ*_ck_*, also known as responsibilities, which represent the probability that each cell *c* belongs to each cluster, *k*.

The M-step uses the assignment probabilities calculated in the E-step to update the model parameters. Specifically, we adjust the subclone priors π*_k_* and the categorical distribution parameters ϕ*_k_* ∈ ℝ^3×^*^d^* to maximize the expected log-likelihood of the observed data, weighted by the assignment probabilities. The categorical distribution parameters ϕ_*k*_ ∈ ℝ^3×^*^d^* for clone *k* represents the probability of observing each (copy number state, encoded region) pair. This step ensures that the parameters better reflect the observed data given the current cluster assignments.

By iteratively updating the assignment probabilities in the E-step and the model parameters in the M-step, the EM algorithm gradually converges to a set of parameters that maximize the likelihood of the data. This iterative process allows the algorithm to find the most probable clustering of the cells based on shared CNA patterns.

##### Initialization

We begin by randomly initializing the subclone assignments, γ*_ck_* of each cell so that cells are distributed randomly between clones. In order to mitigate the effect of local minima when performing this iterative optimization, we perform five random restarts and select the model which has the highest likelihood amongst the five trials.

##### E-step

To determine the optimal assignment of cells to sub-clones, we compute the posterior probabilities (responsibilities) that each cell *M_c_* belongs to sub-clone *k*:

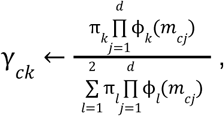

where ϕ*_k_* (*m_cj_*) is the probability of observing *m_cj_* in cluster *k*.

##### M-step

To update the probabilistic sub-clone profiles, we update the parameters π*_k_* and ϕ*_k_* to maximize the expected log-likelihood:

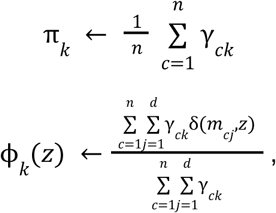

where *z* is a copy number state (–1,0,1) being updated and δ(*a*, *b*) is the Kronecker delta function, which is 1 if *a* = *b* and 0 otherwise.

#### Termination of subclone splitting

We implement two methods to determine whether a clone should be split further. The first (and preferred) option compares the Bayesian information criterion (BIC) score of a model with one clone to that of a model with two clones, and terminates the splitting process if the BIC score does not improve with two clones. Specifically, we calculate

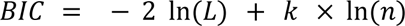

where *L* is the maximum likelihood, *k* is the number of parameters in the model, and *n* is the number of cells. When splitting a clone into two subclones, the model gains additional parameters (new probabilistic profiles and mixing proportions), which incurs a penalty term in the BIC calculation. Only when the improvement in likelihood outweighs this complexity penalty do we proceed with the split. This approach rigorously controls model complexity by requiring substantial evidence that observed variations reflect genuine biological differences rather than stochastic noise^122^.

In cases with limited cell numbers, the statistical power needed for BIC to detect meaningful biological differences may be insufficient. The cell assignment confidence approach provides a complementary criterion that can identify biologically relevant subpopulations even when BIC would prematurely terminate splitting, making it particularly valuable for datasets with fewer cells or more subtle clonal differences.

The second method relies on cell assignment confidence. Using the responsibilities matrix from the EM algorithm, we check the proportion of confidently assigned cells. Specifically, if a cell’s responsibility value exceeds a user-defined threshold (e.g., 0.75), it is considered confidently assigned. If the proportion of confidently assigned cells falls below a user-defined threshold (typically 0.6-0.8), the splitting process is terminated. This ensures further subdivisions are only made when cells show clear membership patterns, avoiding overfitting to noisy data.

#### Post-processing subclones

Inference from scRNA-seq data produces inherently noisy copy number profiles due to technical limitations in the sequencing process. These profiles may contain artifacts and false signals that can lead to the detection of spurious subclones. To ensure the reliability of our phylogenetic analysis, we implement a post-processing step that retains only those clones with sufficient statistical support and biological plausibility, filtering out clusters that likely arise from technical noise rather than true clonal evolution.

In order to mitigate the occurrence of clones derived from noise in the copy number inference process, we require clones to (i) be composed of more than 75 cells and (ii) contain at least one CNA at high frequency. We selected a conservative threshold of 75 cells as a minimum clone size in order to ensure that clones are likely to represent true biological subpopulations rather than technical artifacts arising from the copy number inference process.

For a given clone, we define high frequency CNAs as alterations present in at least 80% of the cells in that clone. The requirement for at least one high-frequency CNA provides additional confidence that the identified clone represents a genuine biological subpopulation with shared genomic alterations.

Clones that do not satisfy these conditions are removed from the phylogenetic analysis, since we do not have sufficient confidence to draw conclusions about the cells they contain.

#### PICASSO benchmarking

To evaluate PICASSO’s phylogenetic reconstruction accuracy, we simulated a series of ground truth CNA trees. Existing single-cell phylogenetic algorithms are not well suited to constructing clone trees from noisy, large scale datasets. For example, CNETML^45^, a maximum likelihood algorithm for deriving phylogenies from copy number profiles, only scales to the low hundreds of cells. We thus compared our ability to recover phylogenetic relationships in these simulations with an agglomerative tree-building algorithm, neighbor joining^120^ (**Extended Data Fig. 4a**).

#### Simulation experiments

We start by generating random binary trees that form the backbone of our CNA simulations, providing a structure on which we can model evolutionary relationships. Each leaf in the tree represents a copy number clone, and branches depict the divergence of clonal lineages over time. Next, we annotate these trees with regions and alterations using a Dirichlet distribution to generate probability vectors for region selection. This distribution allows us to model the relative likelihood of alterations occurring across different genomic regions and capture the biological reality that some regions are more susceptible to CNAs than others.

Each branch is assigned specific alterations (values of −2, −1, +1, or +2) based on a predefined probability distribution [0.5, 0.3, 0.2] that determines only how many alterations will occur per branch (with 0.5 probability of 1 alteration, 0.3 probability of 2 alterations, and 0.2 probability of 3 alterations), reflecting the accumulation of genetic changes as cells evolve. The actual alterations themselves are randomly selected from the set [−2, −1, +1, +2] with equal probability.

To capture the cumulative effect of these alterations, we calculate the aggregated alterations for each leaf node by tracing the path from the root to the leaf. This gives us a comprehensive copy number profile for each clone, accounting for all the genetic changes that occurred along its lineage. Cells are then attached to the leaves of the tree, with the number of cells per clone partially determined by the distribution of clone sizes observed in the data. Specifically, we leverage real-world PDAC data, using half the actual observed clone sizes to balance computational efficiency with biological fidelity while preserving the relative proportions of clonal populations seen in patient samples.

In order to simulate realistic copy number profiles inferred from single-cell data, it is essential to introduce realistic noise, including extensive false positives and false negatives. We introduce noise as follows:

1. **False positives: Add noise to neutral regions**. Simulate false-positive inferred CNAs by randomly selecting a proportion of neutral (no copy number change) regions within the cell profiles and applying random alterations. We perform these simulations across four false positive rate parameter regimes: the false positive rate (0.01, 0.1, 0.2, or 0.3) directly determines the proportion of neutral regions altered—for example, at a rate of 0.1, 10% of neutral regions receive artificial alterations. The magnitude of these alterations follows a distribution derived from observed alterations to ensure realistic noise patterns.
2. **False negatives: Zero-out existing alterations**. Simulate false negatives or loss of signal by zeroing out existing alterations in the cell profiles randomly. The false negative rate directly determines the probability of removing each existing alteration—for example, at a rate of 0.2, each real alteration has a 20% chance of being removed. This stochastic process simulates scenarios where genuine copy number changes go undetected.
3. **Perturb existing alterations**. Simulate CNAs whose presence is correctly inferred, but whose magnitude is not, by slightly increasing or decreasing copy number values. We introduce magnitude perturbations with a probability of 0.1 per alteration, randomly adjusting values by +1 or –1 while preserving the direction (gain or loss). This simulates measurement uncertainty in copy number estimation from sequencing data. These perturbations create a consistent baseline of noise across all experimental conditions, independent of the varying false positive and false negative rates being tested, better reflecting the technical challenges in precise CNA quantification.

We conduct simulation experiments with three replicates across multiple parameter configurations. Each simulation maintains 60 leaves and 110 regions, dimensions comparable to the PDAC tree inferred by PICASSO. By systematically varying false positive and false negative rates (0.01, 0.1, 0.2, and 0.3), we comprehensively evaluate the robustness of both neighbor joining and PICASSO methods under increasingly challenging conditions of data quality.

#### Metric for evaluating PICASSO

We evaluated PICASSO and neighbor joining phylogenies using the triplets-correct metric^123^, which assesses the tree’s ability to reconstruct correct phylogenetic relationships between triplets of cells (**Extended Data Fig. 4a**). For each simulated tree, we sample 10,000 triplets (*a*, *b*, *c*). For each triplet, the ground truth tree induces a phylogenetic ordering on the cells. For example, for triplet (*a*, *b*, *c*), the ground truth phylogenetic relationship of these cells may be ((*a*, *b*), *c*), indicating that cells *a* and *b* share a more recent common ancestor than *a* and *c* or *b* and *c*. In an inferred tree, the triplet is scored as “correct” if the phylogenetic relationship between these cells is accurately recovered.

Since the simulated tree defines leaves as “clones” (groups of cells that cannot be distinguished from each other by CNA profile), some triplets will have no clear phylogenetic relationship; they are siblings in a clone. Unlike PICASSO, neighbor joining computes a fully resolved cell tree. Therefore, when computing the proportion of triplet relationships that are correctly determined, we only consider triplets with clearly defined phylogenetic relationships. By counting the proportion of correctly inferred triplets, the triplets correct metric provides a quantitative measure of the tree’s accuracy, helping to identify discrepancies and assess the overall quality of the inferred phylogenetic tree.

#### PICASSO runtime and memory comparison

Given the large size of scRNA-seq datasets, runtime complexity is a significant concern. The neighbor-joining algorithm, commonly used in phylogenetic analysis, has a theoretical runtime complexity of *O(n*^3^), where *n* is the number of cells, although some implementations of neighbor joining use heuristics to improve performance in practice^124^.

We evaluated run times on simulated datasets with 20,000 cells (**Extended Data Fig. 4b**). Given the large size of the datasets, we used a heuristic implementation of neighbor joining, rapidNJ. We measured the runtimes for both neighbor joining and PICASSO on each dataset across all replicates and found that PICASSO is significantly faster and less memory intensive than neighbor joining.

#### PICASSO robustness testing

To evaluate the robustness and reproducibility of PICASSO, we ran the method five times on the full PDAC dataset of approximately 40,000 single cells and assessed the consistency of the resulting phylogenetic reconstructions (**Extended Data Fig. 4c**). Pairwise comparisons of the clone assignments across runs were quantified using normalized mutual information (NMI) and adjusted Rand index (ARI), widely used metrics for comparing the similarity between two clustering assignments.

NMI measures the mutual information shared between two clustering assignments, normalized to a 0–1 scale:

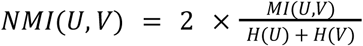

where *MI*(*U*, *V*) s the mutual information between clustering assignments *U* and *V*, and *H*(*U*) and *H*(*V*) are their respective entropies. An NMI score of 1 indicates perfect agreement between clusterings, while 0 indicates completely independent clusterings. The NMI scores we observed were consistently high, averaging around 0.85, indicating strong agreement in the overall clustering structure across runs.

ARI measures the similarity between two clustering assignments by counting pairs of elements that are either assigned to the same cluster or different clusters in both assignments, adjusted for chance:

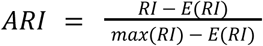

where *RI* is the raw Rand Index, *E*(*RI*) is the expected raw RI and max(RI) represents the theoretical maximum value the Rand Index could achieve for the given clustering problem. The raw Rand Index is defined as:

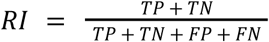

where *TP* is the number of pairs that are in the same cluster in both clusterings, *TN* is the number of pairs in different clusters in both clusterings, *FP* is the number of pairs that are in the same cluster in the first clustering but in different clusters in the second and *FN* is the number of pairs that are in different clusters in the first clustering but in the same cluster in the second.

ARI ranges from –1 to 1, with 1 indicating perfect agreement, 0 indicating random cluster assignments, and negative values indicating worse-than-random agreement. The observed ARI, which is sensitive to both the number and composition of clusters, averaged around 0.6, reflecting a reasonable level of stability given the complexity of the dataset and the stochastic nature of the method.

To further assess consistency at the phylogenetic level, we computed the proportion of triplets (evolutionary relationship between three cells) recovered in each run that matched those identified in a separate run designated as the ground truth. High concordance of triplets across runs demonstrates that PICASSO reliably reconstructs phylogenetic relationships despite inherent variability in clustering.

To further assess the robustness of PICASSO, we conducted a downsampling analysis by randomly subsampling the dataset to 75%, 80%, 85% and 90% of the original dataset (**Extended Data Fig. 4d**). For each downsampled dataset, we ran PICASSO and measured the proportion of triplets in the reconstructed phylogenies that matched the triplets identified in the full dataset, which served as the reference. Across all levels of downsampling, the proportion of correctly recovered triplets remained high, demonstrating the method’s robustness.

### Computational analysis

#### Digital histopathology

Whole slide imaging data were obtained with the assistance of the Molecular Cytology Core Facility at Memorial Sloan Kettering Cancer Center. H&E-stained slides were scanned using a PANNORAMIC scanner (3DHistech, Budapest, Hungary) equipped with a 20x/0.8 NA objective. The resulting data were analyzed using QuPath (version 0.5.1, https://qupath.github.io/).

Adipose and fibrous tissues were annotated by a pathologist using QuPath. Following annotation, the Pixel Classification tool in QuPath was applied with default settings to quantify the areas of adipose and fibrous tissues.

For cell type evaluation, QuPath’s Cell Detection tool was used to identify and analyze tumor cells, stromal cells, and immune cells. Regions containing these three cell types were annotated, and only tumor-cell-containing areas were included in the analysis. A cell classifier was trained using the Object Classification tool in QuPath with default settings, based on pathologist annotations. Features such as nuclear circularity and eccentricity were calculated to characterize the detected cells. The classifications were validated by the annotating pathologist to ensure accuracy.

### WES data analysis

#### WES data preprocessing

Initial processing began with adapter trimming of FASTQ files using cutadapt (v1.9.1) to remove standard Illumina 5’ and 3’ adapter sequences. The trimmed reads were then mapped to the B37 reference genome from the Broad GATK resource bundle using BWA-MEM (v0.7.12). Post-alignment processing included sorting of SAM files and addition of read group tags using PICARD tools (v1.124). The read group information includes sample identifiers, sequencing library identifiers, and Illumina platform information. The sorted BAM files were then processed with PICARD MarkDuplicates to identify PCR duplicates (https://github.com/soccin/BIC-variants_pipeline).

#### Copy number alteration calling

Copy-number alterations in solid tumors were computed from tumor and matched normal tissue WES data using default settings in the FACETS (Fraction and Allele-Specific Copy Number Estimates from Tumour Sequencing) (v0.6.2) algorithm (https://github.com/mskcc/facets-suite)^115^. FACETS provides allele-specific copy number estimates at the level of both gene and chromosome arm.

#### Single nucleotide variant calling

We used the standardized Illumina (HiSeq) Exome Variant Detection Pipeline to detect variants in the output of preprocessed WES data. Following duplicate marking, BAM files are processed according to GATK (v3.4-0) best practices version 3 for tumor–normal pairs. This includes local realignment using ABRA (v2.17) with default parameters, followed by base quality score recalibration using BaseQRecalibrator with known variants from the Broad GATK B37 resource bundle, including dbSNP v138.

Somatic variant calling is performed using muTect (v1.1.7) with default parameters for SNV detection, while somatic indels are identified using GATK HaplotypeCaller with subsequent custom post-processing. A final “fill-out” step computes the complete read depth information at each variant position across all samples using the realigned BAMs. This step applies quality filters requiring mapping quality ≥ 20 and base quality ≥ 0, with no filtering for proper read pairing.

All analyses were performed using a standardized computational environment managed through Singularity (v2.6.0). The complete pipeline source code, including all post-processing scripts, is available at:

- https://github.com/soccin/BIC-variants_pipeline
- https://github.com/soccin/Variant-PostProcess

Additional software versions used in the pipeline include Perl (v5.22.0), Samtools (v1.2), VCF2MAF (v1.6.21), and VEP (v102).

#### SNV and CNA visualization

To visualize the SNV and CNA status of key cancer genes, as well as tumor mutation burden (**Extended Data Fig. 2d**), we used CoMut^125^.

#### snRNA-seq data pre-processing

After quality controls described in the following section below, snRNA-seq generated a total of 73,142 high-quality transcriptomes from 11 samples.

#### Alignment of sequencing reads

All scRNA-seq samples were pre-processed as follows: FASTQ files from the rapid autopsy samples were processed with the SEQC (v.0.2.4) pipeline^126^(https://github.com/dpeerlab/seqc) using the hg38 human genome reference, default parameters and platform set to 10x Genomics v3 3′ scRNA-seq kit. The SEQC (v.0.2.4) pipeline performs read demultiplexing, alignment and unique molecular identifier (UMI) and cell barcode correction, producing a preliminary count matrix of cells by unique transcripts. By default, the pipeline will remove putative empty droplets and poor-quality cells based on (1) the total number of transcripts per cell (cell library size); (2) the average number of reads per molecule (cell coverage); (3) mitochondrial RNA content; and (4) the ratio of the number of unique genes to library size (cell library complexity).

Nuclear transcriptomes from human rapid autopsy samples are expected to have lower RNA content and quality than regular single-cell assays^127^. To obtain a more comprehensive representation of cancer phenotypes we included both FACS and non-sorted samples (see Single-nucleus RNA-seq section), however, non-sorted samples carry a greater degree of low quality nuclei. Therefore, due to the intrinsic lower RNA content and sample quality of flash-frozen snRNA-seq derived transcriptomes, we performed further quality control steps as described in the following sections.

### snRNA-seq data quality control

#### Ambient RNA removal

During nucleus extraction from flash-frozen tissue, cell-free ambient RNA is liberated into the dissociation solution and becomes encapsulated with nuclei during library construction. Ambient RNA contamination can create undesired technical artifacts in single cell data, such as ectopic gene expression and the removal of real biological differences between distinct cell population transcriptomes^128^.

To address this issue, we corrected for ambient RNA expression using CellBender (v.0.1.0)^129^(https://github.com/broadinstitute/CellBender). CellBender is an unsupervised Bayesian model that requires no prior knowledge of cell-type-specific gene expression profiles to identify ambient RNA counts^129^. The approach is based on the principle that ambient RNA contamination will have a more uniform distribution across all cells, whereas cell-specific RNA will display more variable expression patterns. The procedure for removing ambient RNA using CellBlender involved the following steps with default parameters:

##### Quality control

Rapid autopsy snRNA-seq samples (particularly non-sorted samples) have more low-quality droplets with debris and ambient RNA than regular scRNA-seq samples^127^. To increase the signal-to-noise ratio between ambient RNA and real RNA counts, we first performed a lenient QC by removing nuclei with more than 5% mitochondrial genes, and fewer than 127 genes or fewer than 255 reads, and by removing genes present in fewer than 10 cells. The estimated cell number of each batch was inferred with SEQC^126^. We applied CellBender to this initial lenient-filtered snRNA-seq data as follows.

##### Estimation of ambient RNA levels

CellBender estimated levels of ambient RNA for each gene across all nuclei by assessing the distribution of expression levels for each gene and identifying genes with a uniform distribution as candidates for ambient RNA contamination.

##### Subtraction of ambient RNA

Next, CellBender subtracted the estimated ambient RNA contamination from the expression level of each gene in every droplet. This process generated a corrected gene expression matrix with non-transformed integer counts.

##### Evaluation of ambient RNA correction

We selected 5,000 highly variable genes using the variance-stabilizing transformation method^130^. To normalize the data, we scaled each cell to 10,000 reads and applied a log_2_(X+1) transformation. Dimensionality reduction was performed using principal component analysis (PCA) and the top 50 components were utilized for downstream analysis. We constructed a k-nearest neighbor (kNN) graph using *k* = 30 and applied PhenoGraph^19^ to identify distinct coarse cell clusters. Cell-type-specific markers were used post-hoc to evaluate ambient RNA correction. CellBender successfully retained cell-type-specific markers in corresponding clusters, while removing unexpected RNA counts, particularly genes from acinar cells that appeared in other cell types.

#### Filtering low-quality nuclear transcriptomes

Proceeding with the CellBender-corrected count matrix, cells with a low number of detected genes, a low total UMI count (sequencing depth) and a high fraction of mitochondrial counts were designated low-quality cells, as they can represent dying cells with broken membranes^131^. Previous snRNA-seq protocols have also reported that ribosomes can remain attached to the nuclear membrane during nucleus isolation^127,132^; therefore, data were further assessed for library size, total gene counts, mitochondrial and ribosomal RNA content.

##### Library size and gene count thresholds

We removed cells with fewer than 500 RNA counts and fewer than 200 genes.

##### Mitochondrial and ribosomal RNA content thresholds

Since our droplets contained nuclear transcriptomes, we reasoned that mitochondrial and ribosomal RNA should be greatly reduced in high-quality transcriptomes. Hence, we checked for cells with high mitochondrial and ribosomal content. Cells with higher levels of mitochondrial and ribosomal genes primarily belonged to non-sorted samples, suggesting that these droplets contained higher levels of debris, as expected. After manual assessment, we removed droplets with more than 1% of mitochondrial RNA and/or more than 10% ribosomal RNA fractions.

#### Doublet detection

Multiplets (droplets containing more than a single nucleus), predominantly doublets, are an undesired byproduct of library production that create artifactual transcriptomes and confound real biological signal. Homotypic doublets encapsulate two nuclei from the same cell type, and heterotypic doublets capture two different cell types, leading to cell-type mislabeling^131^. Given the challenging task of differentiating single transcriptomes from doublets, using more than one detection approach and comparing results can increase the accuracy of doublet detection^133^. We used DoubletDetection^134^ and Scrublet^135^, two of the top-performing doublet detection algorithms^136^, and further inspected identified doublets to confirm larger library size compared to singlets, as well as expression of conflicting gene markers. For each sample independently, we visually compared putative doublet and singlet total count distributions together, and their clustering distribution in UMAP projections.

##### DoubletDetection

DoubletDetection is a machine-learning algorithm for identifying doublets in scRNA-seq data^134^(https://github.com/JonathanShor/DoubletDetection). It generates synthetic doublets, clusters them together with the original data using PhenoGraph^19^, and assigns a score and *p*-value for clusters with enriched synthetic doublets using a hypergeometric test. We used DoubletDetection separately in each sample raw snRNA-seq count matrix with default parameters.

##### Scrublet

Scrublet^135^(https://github.com/swolock/scrublet) simulates doublets from the observed data and uses a kNN classifier to calculate a continuous doublet_score (between 0 and 1) for each transcriptome. The score is automatically thresholded to generate predicted_doublets, a boolean array that is True for predicted doublets and False otherwise. We used Scrublet independently for each sample’s raw snRNA-seq count matrix with default parameters.

We found the results from both methods to be complementary and removed cells identified as doublets by either method. Transcriptomes passing library size, mitochondrial, ribosomal and doublet detection criteria were retained and the data matrices concatenated into a single matrix for downstream analysis.

### snRNA-seq data analysis

#### Feature selection, normalization, and variance stabilization

Following quality control, we selected 5,000 highly variable genes (HVGs) using the ‘seurat_v3’ flavor in scanpy (v.1.9.8)^137^(https://github.com/scverse/scanpy), which computes a normalized variance for each gene on the raw counts^130^. Other parameters were set as default. To normalize the data we scaled each cell to 10,000 reads. The normalized counts were then log-transformed (base 2).

#### Dimensionality reduction and visualization

PCA of the log-normalized matrix was performed using the ARPACK solver on the selected HVGs. We retained the first 50 principal components (PCs), which explained 33.5% of the variation in the data, and constructed a kNN graph using *k* = 30. To visualize the data, UMAP was applied to the PCA-reduced data and a minimum distance of 0.1.

Since non-cancer cells from different libraries were well integrated (**Extended Data Fig. 1d**) we did not perform any batch correction on our data. Differences between samples from different anatomical locations were regarded as potentially biologically driven.

#### Gene signature scores

To generate all gene signature scores in our study, we used the Scanpy score_genes function^138^, which calculates the mean expression of genes of interest subtracted by the mean expression of a random expression-matched set of reference genes. To control for gene set sizes, we selected the random reference set to be the same size as the gene set of interest. Other parameters were set to default.

#### Cell-type annotation

##### Cancer cell-type annotation

To annotate cell types, we first sought to discern cancer cells from non-cancer cells. The tumors harbor a truncal *KRAS^G12V^* mutation, detected both by MSK-IMPACT^139^ and WES mutation calling; therefore, we used two independent but complementary *KRAS* signatures from the literature to generate a KRAS_signaling score per cell:

*KRAS_PDAC*^20^: This signature of 36 genes is based on differential expression between epithelial cells in wild-type *KRAS* and *KRAS*-knockout mouse tumors. We used the human orthologs provided in the signature.

*KRAS_addiction*^21^: This signature was generated by comparing human lung and pancreatic cancer lines that require *KRAS* to maintain viability with those lines that do not; all lines harbored *KRAS* mutations and were treated with short hairpin RNAs to deplete *KRAS*. The resulting signature is specific to *KRAS*-dependent cells, and is associated with a well-differentiated epithelial phenotype also observed in primary tumors.

We scored these signatures separately, and although high-scoring cells for the two signatures did not overlap fully, both robustly identified the same clusters; thus, we used the union of *KRAS_PDAC* and *KRAS_addiction* to generate the *KRAS_signaling* signature for cancer cell annotation. Positive clusters were confirmed by CNA profiles inferred from the scRNA-seq data using inferCNV^22^, as described in the following section.

##### Non-cancer cell-type annotation

To label non-cancer cells, we clustered all cells using PhenoGraph^19^ with default parameters on the previously obtained PCs, and used literature-curated canonical cell-type-specific markers (**Supplementary Table 1**) to annotate the clusters. For clusters related to smooth muscle cells, MUC1/MUC6 epithelial cells, and adipocytes, no initial cell-type identity could be discerned. Therefore, we ranked the genes underlying each cluster using the Scanpy function scanpy.tl.rank_genes_groups with the sparse matrix and default parameters. Reference clusters were set to ‘rest’ as well as adjacent clusters with known cell-type identity for increased granularity. Genes among the top 20 ranked genes were used to identify the cell identity of those clusters.

#### Inferring copy number alterations from snRNA-seq data

To infer chromosomal CNAs in tumor cells, we ran inferCNV (v1.10.0)^22^(https://github.com/broadinstitute/inferCNV) and copyKAT (v1.1.0)^32^(https://github.com/navinlabcode/copykat) using the Python API of these algorithms implemented in the infercnvpy package (v0.1.0). We ran both packages using default parameter settings, and used non-cancer cell types as the diploid reference. InferCNV was run with a window size of 100 genes and a step size of 1, to balance the detection of focal and broad CNA events.

#### Phylogenetic inference in rapid autopsy data

We used ductal and acinar cells as reference normal cells for IntegrateCNV. The algorithm returned a matrix containing copy numbers for 43,949 cells in 116 genetic regions. We only took the subset of cells annotated as tumor, and expanded this matrix to a ternary matrix, as described above. We then removed features that are highly similar across all cells by filtering out features that are modal with frequency 99% or higher, reasoning that small variations in copy number (frequencies below 1%) are likely noise.

We applied PICASSO to this input data and required that each cell have a UMI count greater than 750 and that each clone contains at least 75 cells, generating 66 clones. As a final filtering step to remove noisy clones data from the phylogeny, we required each clone to have at least one CNA at a prevalence greater than 80% to be considered valid. We reason that clones without highly prevalent CNAs are not likely to be well-supported and may represent ‘noise’ clones with cellular CNA profiles that are inconsistent with more well-defined clones. Removing four such noisy clones left a total of 62 clones (95-1,613 cells per clone, median = 617.5 cells) containing 40,994 cells in the phylogeny (**Figs. 5d, 6a** and **Extended Data Fig. 7d**).

We defined a primary clone as containing at least 50% of cells from the primary tumor, yielding four primary clones in the data (**Fig. 2b**). As a proxy for the metastatic behavior of each primary clone, we calculated the proportion of non-primary cells within each clone, with higher values indicating greater dissemination (**Fig. 2c**).

#### AC5 clone assignment

To confirm that primary cells expressing the archetype cluster 5 (AC5) program were strongly associated with advanced clones (**Extended Data Fig. 7d**), we focused on the two advanced AC5 clones with the most primary cells (clones I and J, bearing 7 and 9 cells, respectively). We compared the CNA profiles of these cells to the clone profiles (CNA change probabilities at each site) of their assigned clones as well as the clone profiles of a representative clone (clone 1-1-0-1-1-0) with a majority of primary cells.

We also computed the log-likelihoods of the primary cells CNA profiles in these clones, compared them with those of all other cells in the clone, and found that they exhibited median levels of clone confidence compared to the other (primarily stomach and liver) cells in the clones.

#### Pairwise diffusion distances of pancreas primary archetype 5 cells

To quantitatively evaluate the similarity of pancreas primary AC5 cells with metastatic cells versus other pancreas primary cells we compared the pairwise diffusion distances from all primary AC5 cells to all metastatic AC5 cells and to all other primary cells separately. We used the ‘scipy.spatial.distance.cdist’ function with the metric = euclidean on the diffusion map coordinates. This computes the distance between each pair of the two collections of inputs.

#### Archetype analysis

We used archetype analysis to identify optimal phenotypes (representing adaptive processes) among cancer cell transcriptomes, which may be shared or specific to one or more tumor sites. Archetype analysis identifies the vertices of a convex polytope—an approximation of a convex hull that encapsulates the data in phenotypic space^140^, which in our case is diffusion space. Archetypes often correspond to the extremes of single diffusion components, thus we identified *m + 1* archetypes, where *m* is the number of diffusion components as computed in the section “diffusion components” below (**Fig. 3a** and **Extended Data Fig. 5a,b**). To understand the gene programs that cancer cells used to adapt to different metastatic sites, which likely pose unique challenges and stresses, we computed archetypes in each tissue independently as described below.

#### Archetype analysis per tumor site

First, we partitioned the data by tumor site (pancreas primary, liver, omentum, peritoneum, diaphragm, stomach, lymph node). Each site was normalized independently by scaling each cell to 10,000 reads and applying a log_2_(X+1) transformation.

The selection of the number of HVGs is crucial for capturing meaningful biological variability while minimizing technical noise. Too few HVGs (<500) risks losing important biological variation, while too many HVGs (>5,000) increases noise without adding significant biological variation. In general, our study and others with large data sets (>50,000 cells) and diverse cell types select around 5,000 HVGs. For medium size datasets (5,000–50,000 cells) and less cell-type diversity, 2,000–3,000 HGVs are recommended. To perform archetype analysis per site, which includes only cancer cells from the same organ (470–23,950 cancer cells per organ, median 4,031), we computed 2,000 HVGs using the ‘seurat_v3’^130^ method in scanpy.

We computed 50 PCs using the svd_solver = ‘arpack’ on the HVGs on each dataset. Sites included PDAC primary (3,479 cells, 40% variance explained by PCA), liver (4,031 cells, 36% variance), peritoneum (23,950 cells, 36% variance), lymph node (470 cells, 44% variance), stomach (4,075 cells, 35% variance), diaphragm (6,137 cells, 37% variance) and omentum (3,305 cells, 34% variance). We then computed the kNN graph with *k* = 30 neighbors on the PC space representation (X_pca). We chose 30 neighbors to balance between adding noise (<20 neighbors) and losing biological variation (>50 neighbors) in the medium size datasets we analyzed.

To visualize each site separately, we computed UMAP (min_dist = 0.1) and FDL with default parameters on the kNN graph (**Extended Data Fig. 5c**). We then clustered each dataset using Leiden clustering in scanpy with default parameters and further assessed cell quality and cancer cell purity in each cluster. We detected some outlier clusters with low library size in lymph node (n = 12 cells) and stomach (n = 226 cells) and non-cancer cell contamination in liver (n = 85 cells) data partitions. Given the objective of archetype analysis in detecting extreme data points in the multidimensional space, we removed those cells from each data partition and from the entire dataset.

##### Diffusion components

Given the presence of different cell-state densities in the data, we used an adaptive anisotropic kernel^63^, which adjusts the local bandwidth (sigma) based on local density, to compute diffusion maps. This can give more flexibility in regions with different densities, improving resolution in sparse areas and reducing over-smoothing in dense areas, compared to the fixed anisotropic Gaussian kernel with a predefined scale (sigma) in scanpy, which is more appropriate for relatively uniform cell-state density datasets.

With the adaptive anisotropic kernel, we computed 10 diffusion components (DCs) on the PC projections of the data and calculated their corresponding eigenvalues and the diffusion operator. We used the eigenvalue knee point to determine the number of DCs for each site. Archetypes were calculated on the DCs using the Python implementation of the PCHA algorithm with *delta* = 0. Archetypes were identified independently 10 times to assess robustness, and the nearest real cell to each archetype was identified using Euclidean distance in the diffusion space.

##### Archetype neighborhoods

We next sought to annotate each archetype based on gene expression. Since each archetype is identified as a single cell, we enhance statistical power by defining archetypal neighborhoods, consisting of each archetype’s most similar cells in diffusion map space. The neighborhoods are defined such that they include enough cells to enhance the robustness of inference, while maintaining the archetypal phenotype and distinction between archetypes. Importantly, different metastatic sites have different numbers of cancer cells, archetypes and the density of cells in the phenotype space varies. To account for all these differences, for a given archetype A in a given tissue, we calculate the diffusion distance (D) to its nearest archetype and define the neighborhood for A as the set of cells which are within a fraction of D. This ensures no overlap between the archetypal neighborhoods, thereby maintaining their distinctions. Parameters used for each site are: PDAC primary DC fraction distance = 1/3 (91–1,571 cells per neighborhood); liver DC fraction distance = 1/3 (63–696 cells); peritoneum DC fraction distance = 1/4 (311–1,098 cells); lymph node DC fraction distance = 1/2 (36–112 cells); stomach DC fraction distance = 1/3 (315–997 cells); diaphragm DC fraction distance = 1/3 (160–2,037 cells); and omentum DC fraction distance = 1/3 (43–417 cells). To visualize archetype neighborhoods, we colored the selected neighborhood cells on the FDL projections (**Extended Data Fig. 5c**).

##### Differential gene expression

For each tumor site, DEGs were calculated for each archetype neighborhood versus all other neighborhoods from the same site, using raw counts. Genes expressed in fewer than 5% of cells in each group were filtered out to reduce noise. Differential expression was performed using diffxpy (https://github.com/theislab/diffxpy) with a Wald test, considering DEGs with log_2_ fold change > 0.05 and *q*-value < 0.01.

#### Robustness analysis of archetype neighbors

We tested the robustness of our archetype analysis and archetype neighborhood selection by downsampling library size to various extents for each organ separately. For this, we downsampled counts from each tumor site raw counts data using ‘sc.pp.downsample_counts’. We set the count_per_cell parameter to be 10% or 20% of the original library size, resulting in a randomly downsampled dataset. For each site and downsampling level, we repeated the analysis 20 times with a different random seed for subsampling.

We repeated the entire archetype analysis process using the same parameters as described above in the subsampled data sets. We then compared the selected archetype neighborhoods using the Jaccard metric, which measures the similarity between two sets of elements by quantifying how many elements (archetype neighbor cells) the sets have in common relative to their total unique elements.

Even at higher synthetic dropout rates, the archetype neighborhoods remained stable (>0.75 Jaccard similarity), confirming that the selected archetype neighborhoods are robust and that highly expressed genes do not disproportionately drive the results.

#### Cell-density estimation

To evaluate if archetype neighborhoods were driven by the cell-state density distribution in the high-dimensional space, we estimated the cell-state density of each tumor site partitioned data using Mellon^66^(https://github.com/settylab/Mellon) with default parameters. Mellon is a non-parametric cell-state density estimator based on a nearest-neighbors-distance distribution. It estimates cell-state densities from high-dimensional representations of single-cell data using a Gaussian process. We preprocessed and calculated cell-state densities for each tumor site separately following the basic tutorial (https://github.com/settylab/Mellon/blob/main/notebooks/basic_tutorial.ipynb).

#### Integrated archetype clusters

To capture possible shared processes, we subsetted the data to include all cells labeled with an archetype, and all genes that were included in any DEGs associated with any archetype in any organ. All 14,826 archetype cells were combined into a single matrix which we median-count normalized, log-transformed counts. PCA (56 PCs, 20% variation explained) was followed by kNN graph construction (*k* = 30 neighbors), Leiden clustering (*resolution* = 1), PAGA^141^, and UMAP visualization (*min_dist* = 0.1, *init_pos* = PAGA) (**Extended Data Fig. 5e,f**).

##### Level 1: Differentially upregulated genes

Differential expression using diffxpy (https://github.com/theislab/diffxpy) (Wald test, DEGs with log_2_ fold change > 1 and *q*-value < 0.05) was calculated for each integrated archetype versus all other archetypes (**Extended Data Fig. 5d**).

##### Level 2: Gene modules

Cancer cells are able to express a variety of gene expression programs that may resemble a distinct modular processes in a physiological setting. To disentangle these gene expression programs we used Hotspot^67^. Hotspot identifies informative genes based on gene-gene autocorrelation in local neighborhoods, using a kNN graph which we generated with weighted_graph = false, n_neighbors = 30, and FDR < 0.05. Gene modules were computed on these genes. Informative genes from Hotspot modules were ranked by local correlation *z*-score. Then pre-ranked GSEA^142,143^ was performed using GSEApy^144^ (https://github.com/zqfang/GSEApy) against selected GSEApy supported gene set libaries (https://maayanlab.cloud/Enrichr/#libraries) and expert-curated gene sets:

*GSEApy libraries:* GO_Biological_Process_2021, MSigDB_Hallmark_2020, Reactome_2016, KEGG_2021_Human, GO_Cellular_Component_2021, GO_Molecular_Function_2021, WikiPathways_2019_Human, and Azimuth_Cell_Types_2021.

*Expert-curated gene sets:* Azimuth_Pancreas_Cells (https://azimuth.hubmapconsortium.org/references/human_pancreas/), PDAC_Subtypes (classical and basal)^94^, PDAC_Signatures^145^, Pancreas_Development^146^, Cancer_Metaprograms^93^, Cell_Cycle^147^, KRAS_signaling^20,21^. Manually-curated gene sets are listed in **Supplementary Table 9**.

The pancreas development^146^ gene set was generated by calculating DEGs (using MAST^148^ with default parameters) between emergent endodermal pancreas (clusters marked by *PRX1*) and other emerging endodermal organs. Then we mapped the gene orthologs between mouse and human genomes.

##### Level 3: Archetype genes

DEGs and genes with modules whose mean expression is highest in a given archetype were used to characterize the archetype. This level of annotation ensures that genes are specifically upregulated in the archetype over other archetypes. Level 3 genes in each archetype were manually inspected to confirm GSEA results and to increase the granularity of the archetype descriptions. Archetypes with low normalized enrichment scores from GSEA were further inspected and labeled according to level 3 genes.

CZ CELLxGENE Discover^149^ was used to annotate archetype 5. Gene expression of archetype cluster 5 genes was evaluated in CZ CELLxGENE. Higher average expression was observed in intestinal, stomach, and gallbladder tissues. Specifically, epithelial cell types were then evaluated for expression of AC5 genes (**Fig. 4c**):

**Intestine:** endocrine cell, columnar/cuboidal epithelial cell, secretory cell, enterocyte, epithelial cell, mesothelial cell, glandular epithelial cell, goblet cell, absorptive cell, brush cell, intestinal crypt stem cell of colon, intestinal epithelial cell, intestinal enteroendocrine cell.

**Stomach:** enterocyte, epithelial cell, ciliated epithelial cell, columnar/cuboidal epithelial cell, glandular epithelial cell, secretory cell, enteroendocrine cell, endocrine cell, peptic cell, mucous cell of stomach, parietal cell, glandular cell of esophagus, epithelial cell of esophagus, intestinal epithelial cell, brush cell, type G enteroendocrine cell, mucus secreting cell, goblet cell, intestine goblet cell.

**Gallbladder:** epithelial cell, secretory cell, goblet cell

**Pancreas:** pancreatic ductal cell, epithelial cell of pancreas To annotate AC2 at a more granular level, we also used the Kyoto Encyclopedia of Genes and Genomes (KEGG) database^150^. Specifically, we used the KEGG Mapper Search Tool (https://www.genome.jp/kegg/mapper/search.html), which searches various KEGG objects, including genes, KOs, EC numbers, metabolites and drugs, against KEGG pathway maps and other network entities. Then the top matching KEGG objects found were used to explore and annotate the biology of AC2 modules:

##### Fatty acid and cholesterol biosynthesis

Metabolic Pathways (hsa01100) and Fatty Acid Metabolism (hsa01212).

##### Oxidative stress and detoxification

Metabolic Pathways (hsa01100), Ferroptosis (hsa04216), Glutathione metabolism (hsa480) and Chemical carcinogenesis - reactive oxygen species (hsa05208).

#### Archetype cluster annotation

Archetype cluster annotation was performed by first considering NES and the specific archetype genes deemed significant by GSEA. The NES g were used as an initial general guide. Higher priority was then given to the specific gene modules and genes to annotate clusters in a granular and specific manner. CZ CELLxGENE Discover^149^ was used to annotate archetype 5 since only the PDAC Adhesive gene program^145^ was significantly enriched.

#### Comparison with Leiden clustering

To compare archetype clusters and Leiden clusters we first clustered the cancer data using ‘sc.tl.leiden’ with default parameters. Then the same level 1 and 2 steps employed for archetypes were used to annotate gene programs associated with Leiden clusters. We compared the archetype and Leiden clusters’ DEGs using Jaccard Similarity.

#### Archetype analysis and annotation of RA19_21 peritoneum metastases

Two PDAC peritoneum metastases were harvested from the rapid autopsy RA19_21 and snRNA-seq data were collected following the same protocol described in snRNA-seq data pre-processing, scRNA-seq data analysis, and Archetype analysis sections above for RA19_10. Data preprocessing and quality control were also performed using the same workflow. The same archetype analysis and gene program annotation workflows were performed for these peritoneal metastatic samples to evaluate expression of the lipid metabolism and oxidative stress programs found in AC2. No integration of archetype clusters was required since only peritoneal metastases were analyzed.

#### Entropy of archetype distributions

We sought to determine whether each clone exhibits a greater diversity of archetypes than expected by chance, which would indicate phenotypic plasticity across the phylogeny (**Fig. 6b**). For each clone, we computed the Shannon entropy of the observed archetype distribution as a measure of phenotypic diversity. Shannon entropy, *H*, is calculated as:

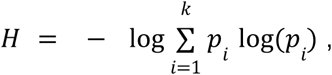

where *p_i_* is the proportion of cells within the clone assigned to archetype *i* and *k* = 18 is the number of unique archetypes. This entropy metric allows us to quantify the spread of archetype diversity within clones, with higher entropy values indicating more even and diverse distributions of archetypes.

#### Null model comparisons

To contextualize the observed entropy and evaluate whether the diversity observed within clones is greater than expected by chance, we compared our results to several null models. Each null model simulates archetype distributions under different assumptions, providing a range of baselines. In order of decreasing expected diversity, they are:

1. **Random assignment.** Archetypes are assigned to cells randomly across all clones, with probabilities matching the global frequencies of each archetype. This model retains the overall prevalence of each archetype, but removes any structure associated with clone or site, simulating a scenario in which cells randomly adopt a phenotype without any constraints.
2. **Site-constrained random shuffle.** Archetypes are assigned to cells randomly *within sites*, preserving each site’s archetype frequency distribution. This model retains the overall presence of each archetype and its prevalence within each site, but removes any structure associated with clones.
3. **High site–archetype concordance assignment.** Archetypes are assigned to cells to minimize the dispersion of archetypes across sites. Greedily, we assign archetype labels to cells within sites in a way that retains the global archetype frequency, but not the per-site frequencies. This model shows the expected diversity if cells were insufficiently plastic to adopt the same phenotype in multiple distinct sites.
4. **Site entropy within clones.** We compute the entropy of site distribution within each clone, ignoring archetype labels, to model the simplistic scenario in which the site drives all phenotypic variation.

PLASTRO quantifies clone plasticity.

The entropy of archetypes within clones provides information about the number of phenotypes a clone can adopt. However, to measure lineage plasticity—which we define as the cells’ inherent capability to flexibly transition between various lineage states or phenotypes—we must examine cellular phenotypes in the context of their phylogenetic relationships. This approach allows us to assess the extent to which cells or cell groups adopt distinct phenotypes compared to their evolutionary ancestors.

We leveraged two complementary data modalities to develop metrics for measuring lineage plasticity: PICASSO, which enables reconstruction of phylogenetic relationships, and archetype analysis, which characterizes the breadth of phenotypes present in the cells. Existing methods for measuring plasticity have key limitations, including the need to discretize continuous cell states, dependence on fully resolved cell phylogenies, and sensitivity to neighborhood size hyperparameters. Our integrated approach specifically addresses these concerns.

Yang and colleagues^123^ defined three metrics for quantifying cellular plasticity: scEffectivePlasticity applies the Fitch-Hartigan algorithm to calculate a normalized parsimony score based on discrete Leiden cluster transitions across a phylogenetic tree; scPlasticityAllelic offers a tree-agnostic alternative by measuring the proportion of cells belonging to different Leiden clusters than their closest genetic relatives (determined by edit distance); and scPlasticityL2 combines phylogenetic information with continuous phenotypic measurements by calculating the Euclidean distance in scVI latent space between cells and their tree-defined neighbors, avoiding the limitations of discrete cluster boundaries.

Schiffman and colleagues^106^ introduce phylogenetic correlations to quantify how cellular measurements are distributed across a phylogenetic tree using Moran’s *I* (a measure of spatial autocorrelation) and its bivariate generalization. This approach measures correlation patterns directly across the phylogeny, facilitating analysis of both continuous expression patterns and discrete cell states within their evolutionary context. The method transforms pairwise phylogenetic distances into a weighted matrix, using carefully selected weighting functions. The choice of weighting function is critical as phylogenetic correlations depend significantly on the structure of the normalized weight matrix, and the function selected by the authors only includes cells that are each other’s nearest phylogenetic neighbor. This choice of weighting function may not be suitable for larger scale datasets on the order of tens of thousands of cells.

To address these concerns, we developed PLASTRO, a metric for quantifying plasticity from jointly profiled lineage and scRNA-seq information, without relying on the inference of complete and exact tree topologies, fixed neighbourhood size hyper-parameters or discretization of cell phenotypes. PLASTRO accepts two distance matrices as input: (i) lineage distance, which reflects how similar clones are to each other in evolutionary space, and (ii) phenotypic distance, which reflects how similar clones are to each other functionally. Given these matrices, we can define a lineage neighbourhood and a phenotypic neighbourhood of radius *r* clones for each clone. Each clone’s neighbourhoods comprise its *r* closest cells in lineage and phenotype space, respectively. The key idea behind this approach is that there will be substantial agreement between the lineage neighbourhood and the phenotypic neighbourhood in non-plastic clones; thus, overlap in these neighbourhoods will be high on average. In contrast, highly plastic clones will exhibit phenotypes distinct from other clones in their lineage, and their neighbourhoods will overlap very little on average.

#### Computation of PLASTRO score

PLASTRO accepts a lineage distance matrix and a phenotypic distance matrix as input. Given these matrices, we can define, for each cell, a lineage neighbourhood and a phenotypic neighbourhood of radius *r* cells. The choice of radius clearly has a strong effect on the degree of overlap between phylogenetic and phenotypic neighborhoods. At very small radii, even non-plastic cells may exhibit low overlap by random chance. Conversely, at very large radii, plastic cells will exhibit strong overlap as well, given that each neighbourhood contains nearly all the cells in the dataset. In addition, different radii provide varying signals that help differentiate plastic and non-plastic cells depending on the parameters of the dataset. To circumvent this issue and avoid reliance on neighbourhood size as a parameter of our approach, we measure neighbourhood overlap at varying scales and combine the signal present at each scale.

PLASTRO consists of four main steps:

1. Compute the lineage and phenotypic distance matrices.
2. For each cell, rank all other cells in terms of the distance from that cell in both (a) lineage space and (b) phenotypic space.
3. For a given cell at overlap radius *r*, compute the overlap in their *r* closest cells as defined by phenotypic distance and by lineage distance. This is the number of cells that lie in both the phenotypic neighbourhood of size *r* and the lineage radius of size *r*.
4. Aggregate signal across radii by computing the area under the overlap versus radius graph.

Phenotypic distance matrix.

We calculate the pairwise phenotypic distances between clones using Bray–Curtis dissimilarity, a metric that captures differences in relative abundances and is commonly used in ecological and compositional analyses. Bray–Curtis is particularly suited to compositional data as it accounts for the proportional structure of the data, measuring dissimilarity on a scale from 0 (identical composition) to 1 (completely dissimilar).

The archetype composition for clone *A* is denoted by *a* ∈ *R^k^* where *k* is the number of archetypes and satisfies

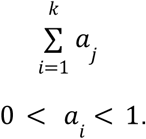

The Bray–Curtis distance between two clones *A* and *B* is then given by

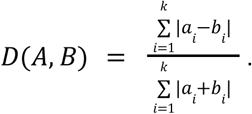

The Bray–Curtis distance ranges from 0 to 1, where 0 indicates that the two samples have identical compositions and 1 that the two samples have completely disjoint compositions (no shared components).

#### Phylogenetic/ lineage distance matrix

We use the phylogeny inferred by PICASSO to construct a pairwise distance matrix between clones; the distance between two clones is given by the number of edges separating them in the phylogeny.

#### Overlap computation

Given a lineage distance matrix *D_L_* and a phenotypic distance matrix *D_P_* constructed on a set of cells, *X*, we compute the overlap for the cell of interest *c* at radius *r* as follows. Denote by *D_L_*(*c*, *r*) the distance in lineage space between cell *c* and its *r^th^* nearest neighbour. Similarly, *D_P_*(*c*, *r*) is the distance in phenotypic space between cell *c* and its *r^th^* nearest neighbour.

We define the lineage neighbourhood of cell *c* at radius *r* as:

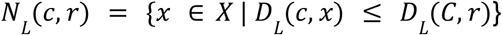

and the phenotypic neighbourhood of cell *c* at radius *r* as:

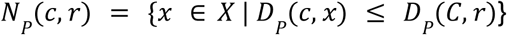

Then, the overlap for cell *c* at radius *r* is defined as the Jaccard similarity of its phenotypic neighbourhood and its lineage neighbourhood:

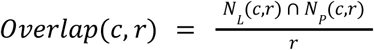

Plastic cells will have lower agreement between lineage and phenotypic neighbourhoods, particularly at lower radii, and thus a lower overlap at that radius on average, compared to less plastic cells.

#### Aggregating signal across radii

To avoid hard-coding a radius which may have a strong effect on the measured plasticity, we aggregate signals across radii by considering overlap size as a function of radius, which is an increasing function bounded by the line *y* = *x*. We compute plasticity as the difference between the area under the line *y* = *x* and the area under the overlap-radius curve.

For more plastic clones, the number of cells in the overlap is lower for smaller radii since the phenotypic neighborhood is highly distinct from the phylogenetic neighborhood, and grows to include all cells as the neighborhood size grows, resulting in a higher plasticity score. For less plastic cells, the overlap proportion is expected to be higher overall, and the overlap-radius curve more closely resembles the *y* = *x* line and thus yields a lower plasticity score.

### Application of PLASTRO to PDAC clones

We apply PLASTRO to compute the plasticity of each clone in our data (**Fig. 6c**). The lineage distance matrix is computed based on the topology of the phylogenetic tree, where clones *A* and *B* have a phylogenetic distance *D_L_*(*A*, *B*) = *n* if there are *n* branches on the shortest tree path between them. The phenotypic distance was computed as described above using the Bray–Curtis dissimilarity between archetype composition of clones.

#### Calculation of global plasticity

We used PLASTRO to calculate plasticity at the clonal level, but to assess global plasticity across the entire biological system, we turned to the Mantel test ^107^, which assesses the correlation between two distance matrices (the phylogenetic and phenotypic compositional distance matrices). The Mantel test is a non-parametric test for assessing matrix correlations and is well-suited for evaluating the phylogenetic signal in data without assuming a specific model of evolution. Mathematically, the Mantel test statistic is computed as

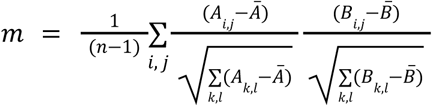

where *A*, *B* ∈ *R^n×n^* are the distance matrices being compared, and 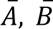 are their respective means. The statistic ranges from –1 to +1, with +1 indicating a perfect positive correlation (as distances in one matrix increase, distances in the other matrix increase proportionally).

A significant positive correlation between the compositional and phylogenetic distance matrices would indicate that clones with closer evolutionary relationships also have more similar compositions. A value of –1 represents a perfect negative correlation (increasing distances in one matrix correspond to decreasing distances in the other, reflecting a complete inverse relationship). A Mantel test statistic near 0 indicates no correlation between the two matrices, such that distances in one matrix do not predict distances in the other, implying that compositional differences are more likely to be driven by factors other than shared ancestry. We used Spearman correlation to measure the association between matrices and performed 1000 permutations to test the significance of the observed correlation.

## Data and software availability

The IntegrateCNV algorithm along with documentation, notebooks and tutorials is available at https://github.com/dpeerlab/integrateCNV.

The PICASSO algorithm, as well as documentation and tutorials for inferring CNA phylogenies and visualizing transcriptional and phenotypic information alongside the tree, is available at https://github.com/dpeerlab/picasso.

---

**Extended Data Figure 1.**
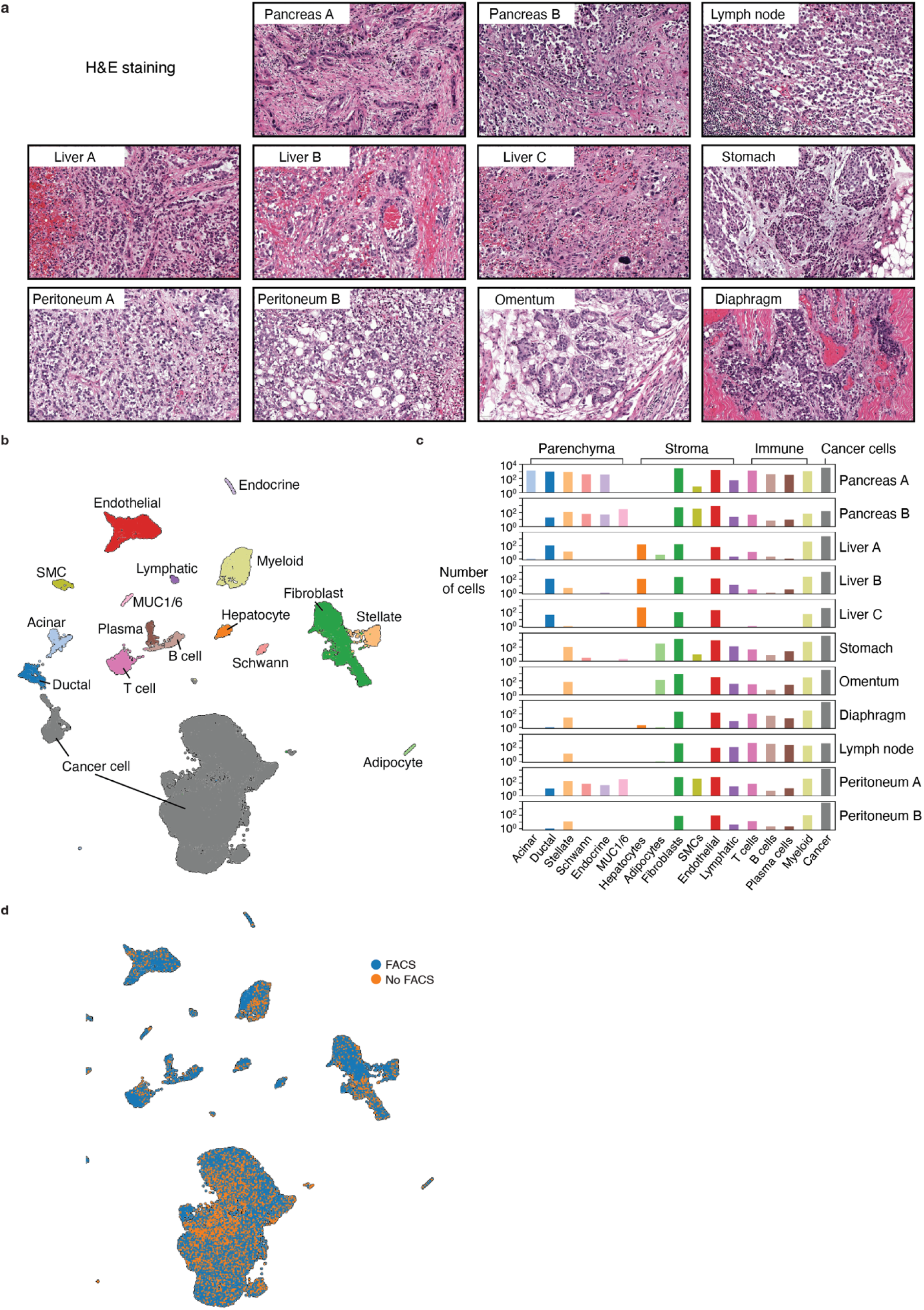
Tumor pathology and gross cellular phenotypes in rapid autopsy data. **a**, Representative images of H&E-stained tumor samples, showing distinct histopathological morphologies. **b**, Uniform manifold approximation and projection (UMAP) of snRNA-seq transcriptomes collected from primary and metastatic tumors (73,142 nuclei), colored by cell type. SMC, smooth muscle cells; MUC1/6, mucinous cells. **c**, Cell type proportions by biospecimen. **d**, UMAP of snRNA-seq transcriptomes colored by FACS protocol (Methods).

**Extended Data Figure 2.**
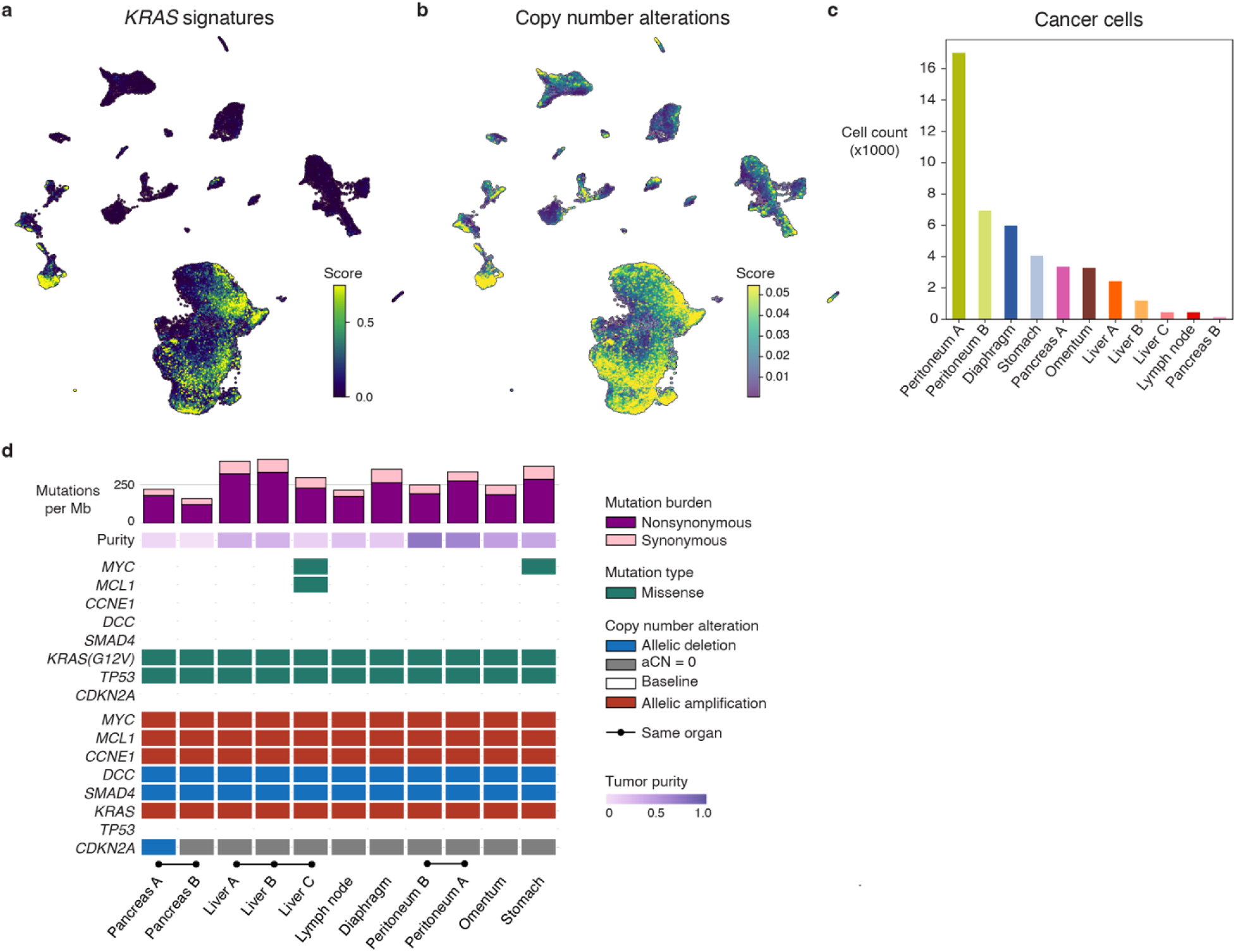
Cancer epithelial cells are distinguished by mutation signatures. **a**, UMAP of all cells, colored by enrichment score of *KRAS* activation signature genes. The score was generated by combining signatures from human cell lines with oncogenic *KRAS^21^* and *KRAS* wild-type PDAC mouse models^20^(**Supplementary Table 1** and Methods). **b**, UMAP of all cells, colored by total CNA burden per cell as measured by inferCNV (Methods). **c**, Cancer cell counts per sample. **d**, Mutational profile of cancer cells from each tumor site, based on WES data. aCN, allelic copy number.

**Extended Data Figure 3.**
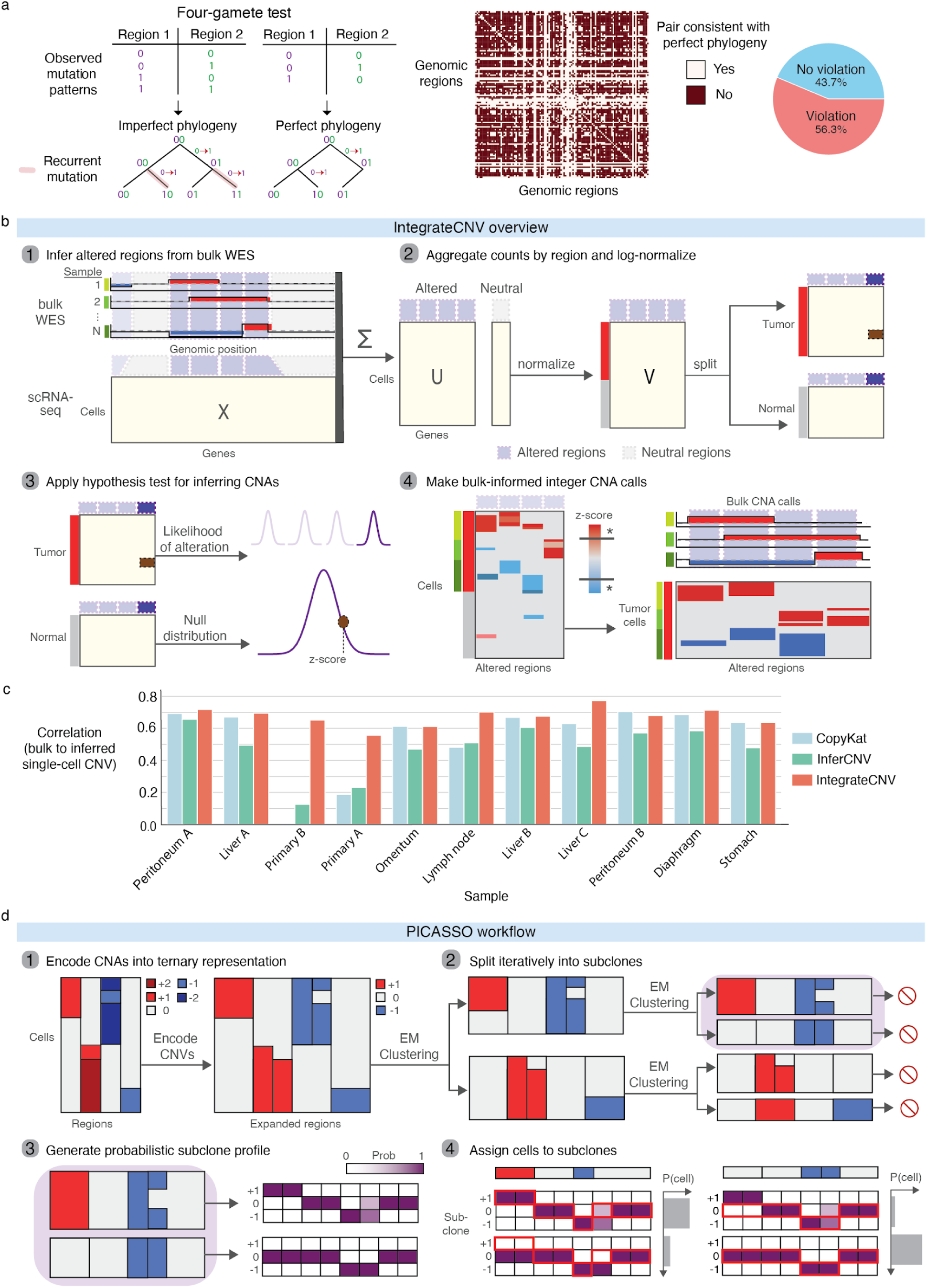
IntegrateCNV identifies copy number variants in single-cell expression data. **a**, Given the absence of recombination between clones, the four-gamete test predicts that it is impossible to achieve a perfect phylogeny if evolutionary characters (mutations) can independently appear more than once (Methods). In cancer, CNAs often recur in different clones. Heatmap shows pairs of genomic regions from inferred single-cell copy number profiles in which all four copy number mutation states are observed and pie chart indicates the overall proportion of pairs that cannot satisfy a perfect phylogeny. **b**, IntegrateCNV identifies regions of the genome that harbor CNAs in bulk WES, then uses a likelihood test on snRNA-seq data to determine whether these regions contain CNAs in each individual cell (Methods). **c**, Benchmarking of single-cell CNA inference methods, based on the correlation between bulk WES (ground truth) data and pseudo-bulk CNA calls. **d**, Detailed schematic of PICASSO, showing probabilistic encoding of integer CNA calls to ternary states (–1, 0, +1 relative to diploid) and iterative cell assignment to subclones (Methods).

**Extended Data Figure 4.**
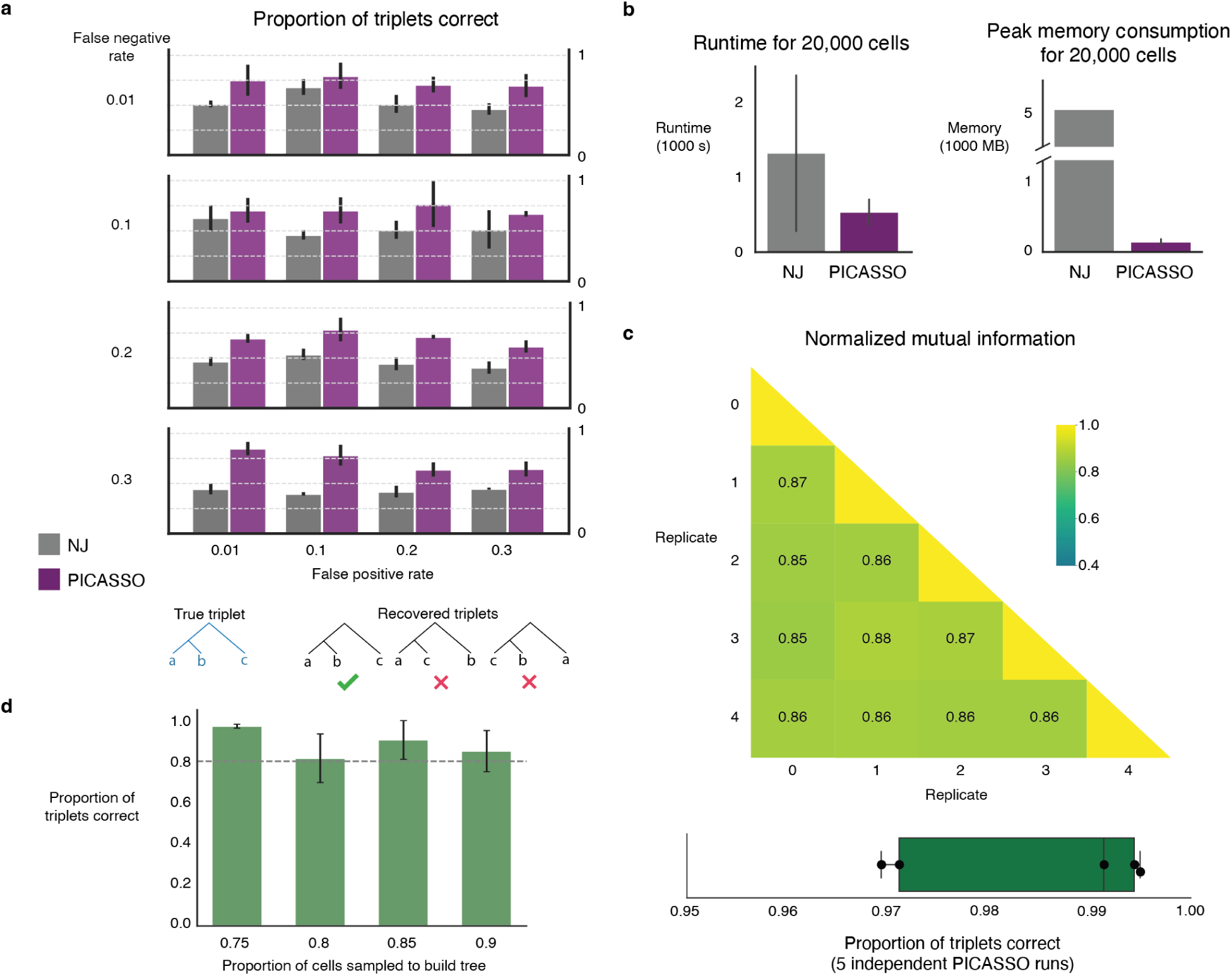
PICASSO enables phylogenetic reconstruction from single-cell expression data. **a**, PICASSO recovers a higher fraction of correct evolutionary triplets than neighbor joining (NJ) in simulated data (Methods). **b**, PICASSO is faster and requires less memory than neighbor joining. **c**, Multiple PICASSO runs generate highly similar clone assignments and tree topologies, as indicated by high normalized mutual information scores between replicates (top) and high proportions of triplets correctly and consistently identified across replicates (bottom). Line, median; boxes, interquartile range; whiskers, full data range. **d**, PICASSO is robust to the downsampling of cells in the dataset, recovering high proportions of triplets correct, compared to the tree inferred from the full dataset. Error bars in **a**,**b**,**d** indicate s.e.

**Extended Data Figure 5.**
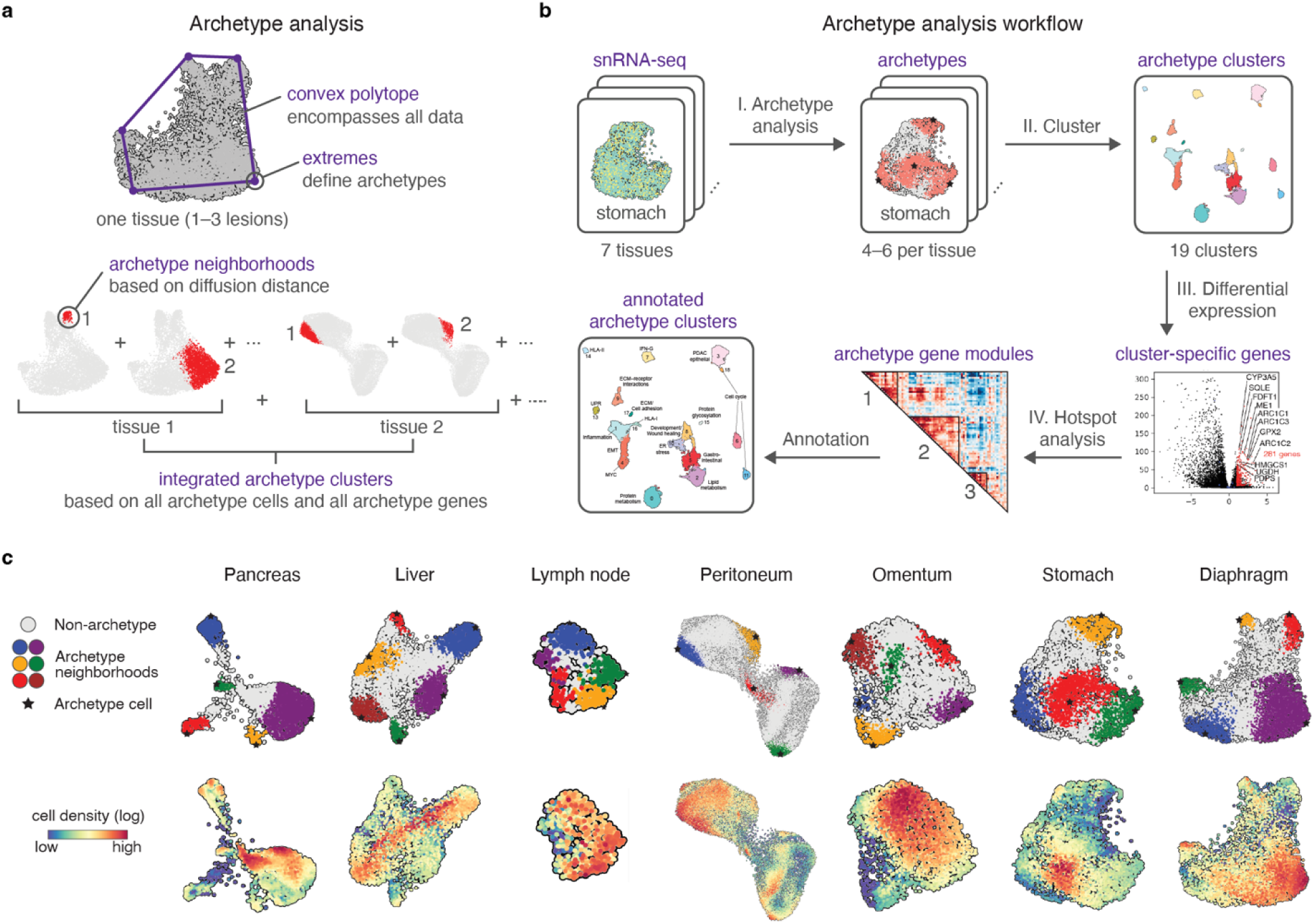
Archetype analysis framework and application to rapid autopsy cancer cells. **a**, Archetype analysis strategy. Archetypes are identified in individual organ sites before integrating into archetype clusters. **b**, Archetype analysis workflow. **c**, FDLs of all cells for each organ, colored by archetypes identified for that site (top) or by cell density (bottom, Methods).

**Extended Data Figure 6.**
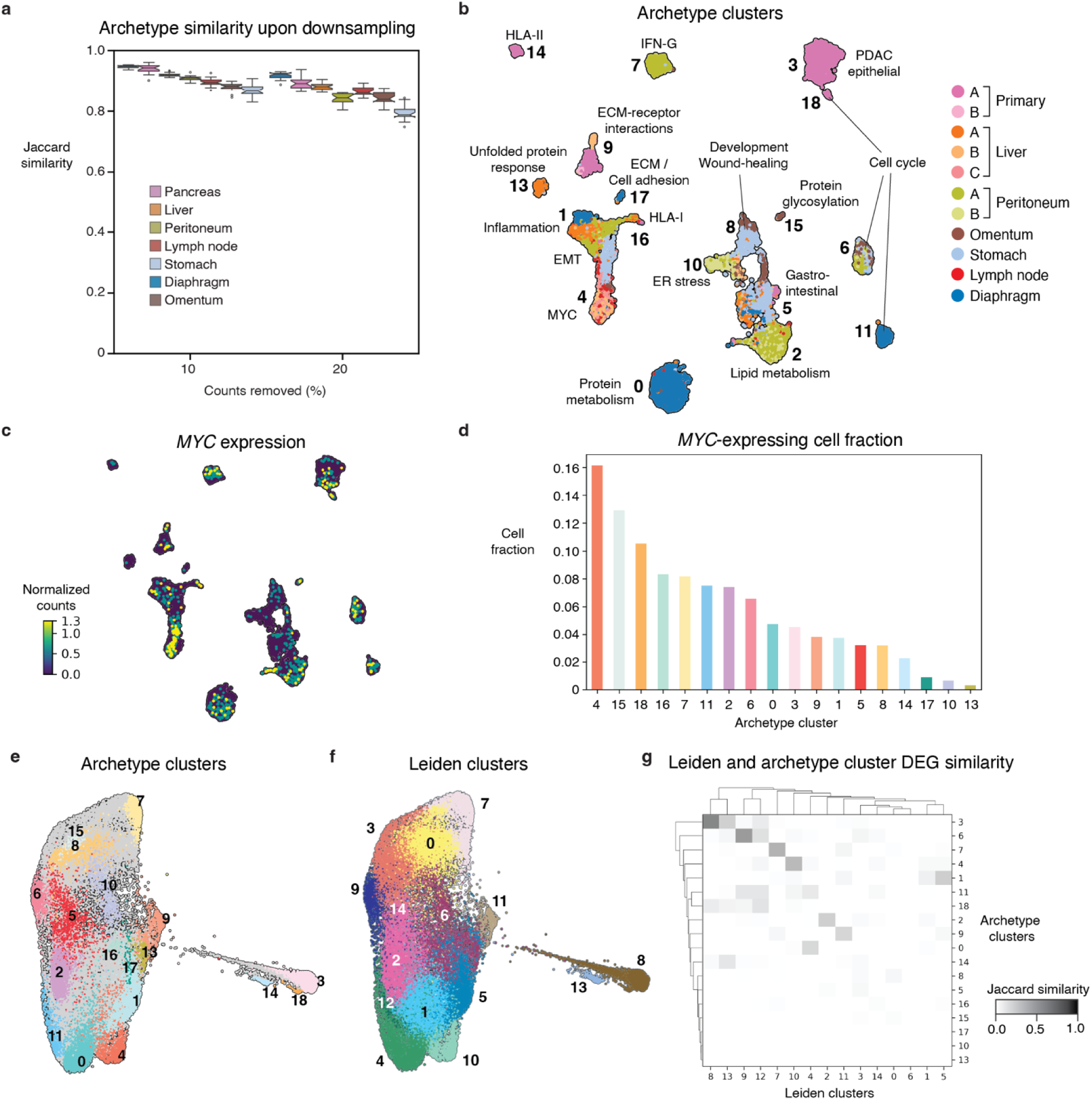
Archetype phenotypes are robust and correspond to biological processes. **a**, High similarity between archetypes before and after downsampling transcript counts (20 repetitions per tissue) demonstrates the robustness of archetype definitions. Line, median; boxes, interquartile range; whiskers, 1.5x interquartile range. **b**, UMAP of clustered archetypal cells, colored by tissue of origin. **c**, UMAP of archetype cells colored by *MYC* expression. **d**, Fraction of cells expressing *MYC* per archetype cluster. **e,f**, FDLs of all cancer cells colored by archetype clusters (**e**) or Leiden clusters (**f**) show distinct non-redundant partial overlap between identified clusters. **g**, Differentially expressed genes for archetypes and Leiden clusters exhibit minimal overlap.

**Extended Data Figure 7.**
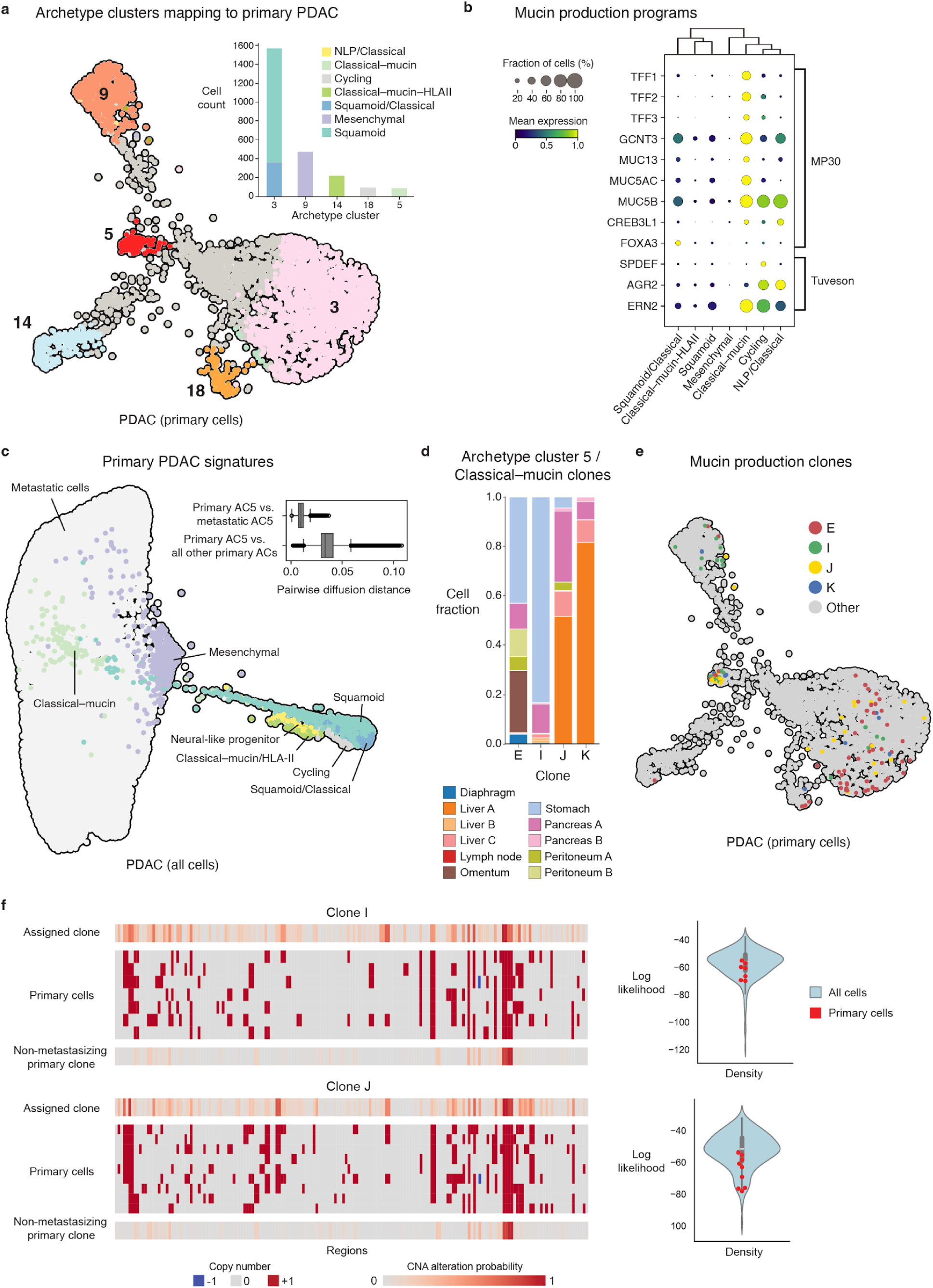
Archetype 5 cells harbor evidence of reseeding the primary tumor from the stomach. **a**, FDL of PDAC primary tumor cells colored by archetype clusters (as in Fig. 3a). Inset, cell count of each archetype with >10 cells from the primary tumor, colored by PDAC phenotype (as in Fig. 2c). NLP, neural-like progenitor. **b**, Expression of mucin production programs across primary PDAC phenotypes. MP30, aberrant induction of mucin production program common in PDAC^93^; Tuveson, mucus production program highly active in PDAC precancerous lesions and the classical subtype^97^. **c**, FDL of all cancer cells colored by primary PDAC signatures. Inset, pairwise diffusion distance (Euclidean distance) between primary AC5 and all other primary cells (mean = 0.04) is greater than between primary AC5 and metastatic AC5 cells (mean = 0.01). **d**, Fraction of cells from each tumor site, for each of the four AC5-annotated clones (E,I,J,K) that contain at least 3 cells derived from the primary tumor. **e**, FDL of PDAC primary tumor cells colored by clones in **d**. **f**, Copy number profiles of primary AC5 cells compared to CNA probability profiles of their assigned clone and a representative non-metastasizing primary clone, for the two AC5 annotated clones with the most primary cells (left). The primary AC5 cells exhibit clone assignment likelihoods typical of all cells in the clone (right).

**Extended Data Figure 8.**
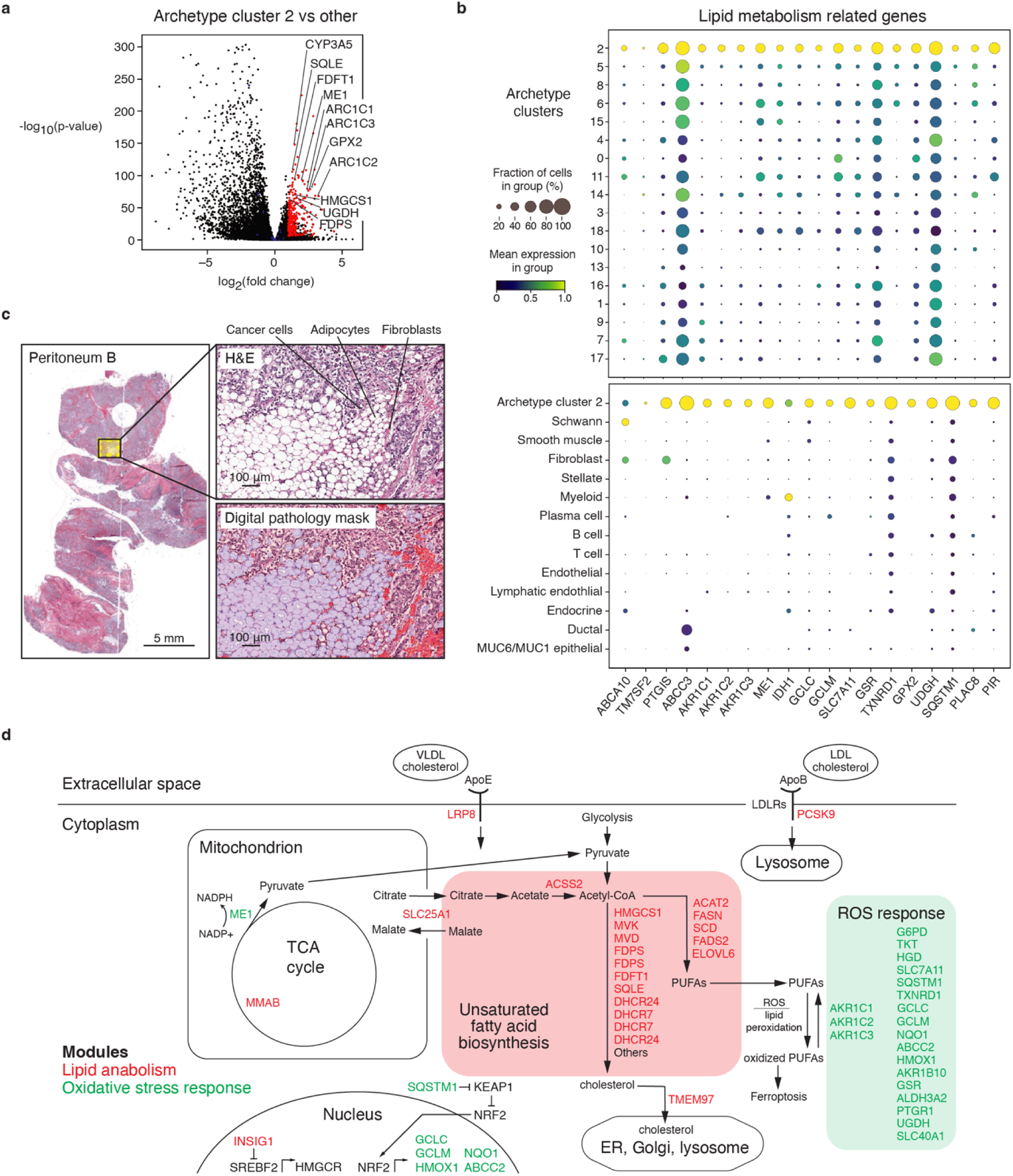
Lipid anabolism is detected in cancer cells in the peritoneum metastases. **a,** Differential upregulation of genes in archetype cluster 2 over all other archetype clusters. **b**, Lipid metabolism genes overexpressed in AC2 metastatic cells compared to other archetype cells (top) and normal cells detected in peritoneal samples (bottom). **c**, H&E showing cancer cells intermingling with adipose cells in peritoneal metastasis. **d,** Lipid anabolism and oxidative stress response pathways, highlighting genes differentially upregulated in AC2 cancer cells in the peritoneum metastases.

**Extended Data Figure 9.**
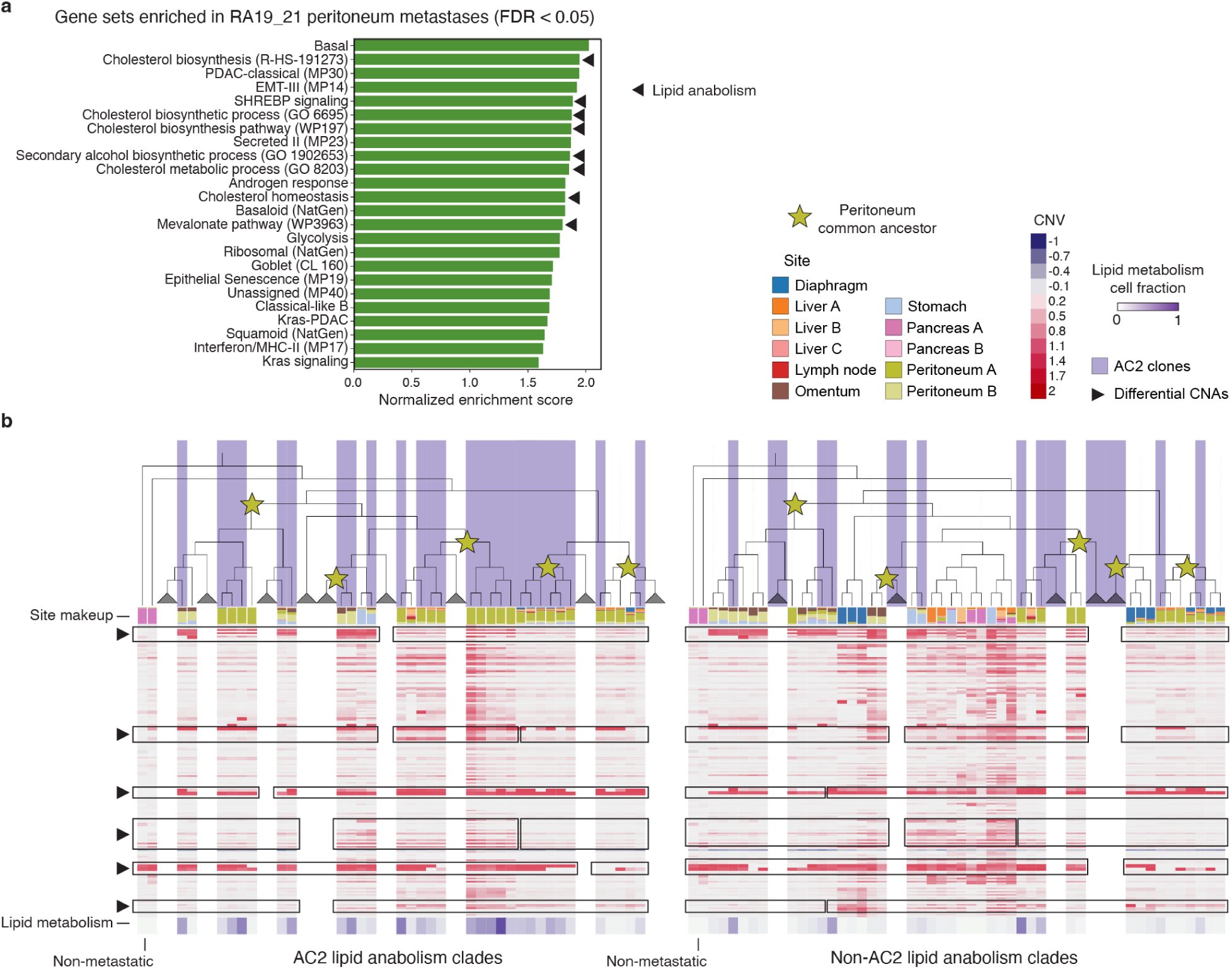
Lipid metabolic rewiring is a prominent feature of peritoneal metastases. **a**, Single sample GSEA normalized enrichment scores of lipid anabolism and oxidative stress AC2 gene modules from snRNA-seq of PDAC peritoneal metastases from an additional rapid autopsy patient. **b**, Cancer cell phylogeny split between AC2 lipid anabolism clades (left) and other clades (right), indicating that the lipid metabolism phenotype is found among clones containing mixtures of peritoneum A and peritoneum B site cells, as well as near-pure peritoneum A or B clones that nonetheless share common ancestors. Stars indicate a most recent common ancestor between lineages that include cancer cells in peritoneum samples.

**Extended Data Figure 10.**
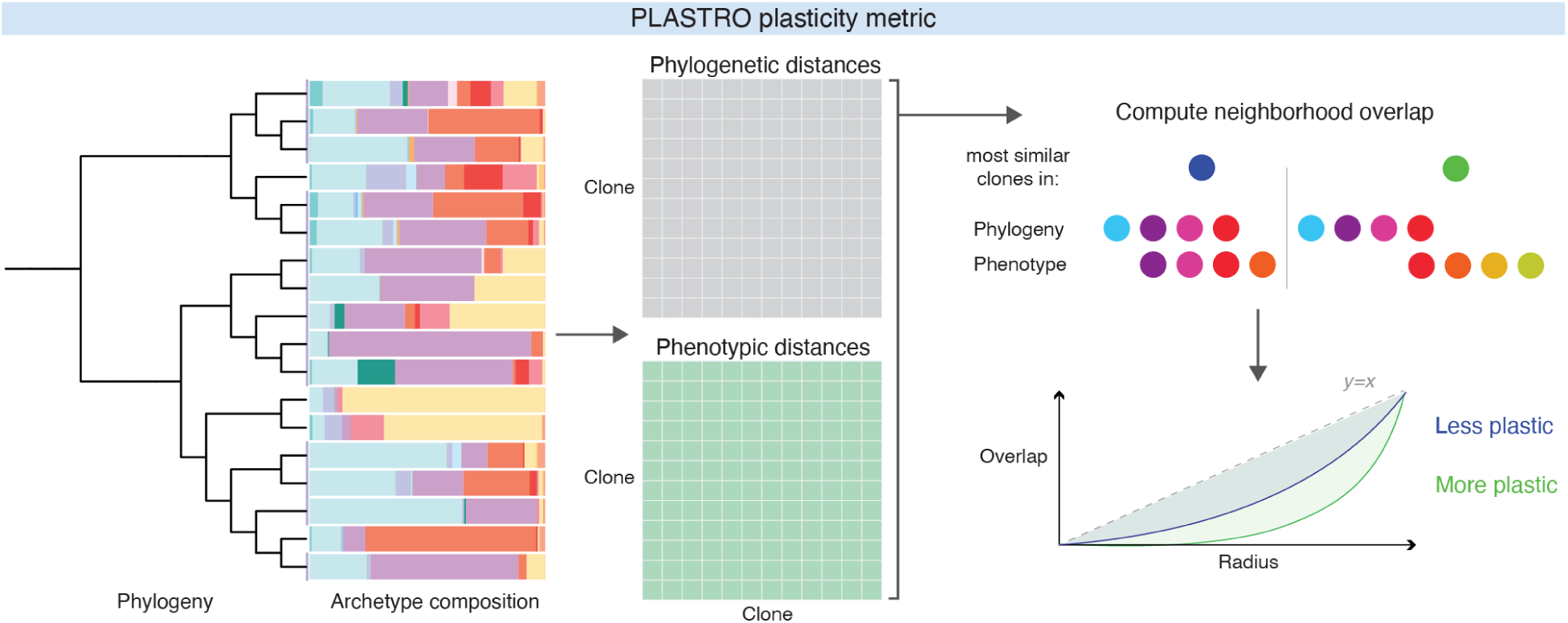
Comparison of phenotype and phylogeny to quantify plasticity. The PLASTRO metric corresponds to the neighborhood-free agreement of phenotype and phylogeny (Methods).

## References

1. Boire, A. et al. Why do patients with cancer die? Nature Reviews Cancer 24, 578–589 (2024).

2. Dillekås, H., Rogers, M. S. & Straume, O. Are 90% of deaths from cancer caused by metastases? Cancer Medicine 8, 5574–5576 (2019).

3. Gerstberger, S., Jiang, Q. & Ganesh, K. Metastasis. Cell 186, (2023).

4. Massagué, J. & Obenauf, A. C. Metastatic colonization by circulating tumour cells. Nature 529, (2016).

5. Hasan, A. M. M. et al. Copy number architectures define treatment-mediated selection of lethal prostate cancer clones. Nature communications 14, (2023).

6. Turajlic, S. et al. Tracking Cancer Evolution Reveals Constrained Routes to Metastases: TRACERx Renal. Cell 173, (2018).

7. Ganesh, K. & Massagué, J. Targeting metastatic cancer. Nature Medicine 27, 34–44 (2021).

8. Nguyen, B. et al. Genomic characterization of metastatic patterns from prospective clinical sequencing of 25,000 patients. Cell 185, (2022).

9. Makohon-Moore, A. P. et al. Limited heterogeneity of known driver gene mutations among the metastases of individual patients with pancreatic cancer. Nature genetics 49, (2017).

10. Flavahan, W. A., Gaskell, E. & Bernstein, B. E. Epigenetic plasticity and the hallmarks of cancer. Science (New York, N.Y.) 357, (2017).

11. Mehta, A. & Stanger, B. Z. Lineage Plasticity: The New Cancer Hallmark on the Block. Cancer research 84, (2024).

12. Moorman, A. et al. Progressive plasticity during colorectal cancer metastasis. Nature 637, 947–954 (2024).

13. Laughney, A. M. et al. Regenerative lineages and immune-mediated pruning in lung cancer metastasis. Nat. Med. 26, 259–269 (2020).

14. Househam, J. et al. Phenotypic plasticity and genetic control in colorectal cancer evolution. Nature 611, (2022).

15. Goyette, M. A., Lipsyc-Sharf, M. & Polyak, K. Clinical and translational relevance of intratumor heterogeneity. Trends in cancer 9, (2023).

16. Iacobuzio-Donahue, C. A., Litchfield, K. & Swanton, C. Intratumor heterogeneity reflects clinical disease course. Nature cancer 1, (2020).

17. Iacobuzio-Donahue, C. A. et al. Cancer biology as revealed by the research autopsy. Nature reviews. Cancer 19, (2019).

18. Mullen, K. M. et al. The Evolutionary Forest of Pancreatic Cancer. Cancer discovery 15, (2025).

19. Data-Driven Phenotypic Dissection of AML Reveals Progenitor-like Cells that Correlate with Prognosis. Cell 162, 184–197 (2015).

20. Ischenko, I. et al. KRAS drives immune evasion in a genetic model of pancreatic cancer. Nature Communications 12, 1–15 (2021).

21. Singh, A. et al. A gene expression signature associated with ‘K-Ras addiction’ reveals regulators of EMT and tumor cell survival. Cancer Cell 15, 489–500 (2009).

22. Patel, A. P. et al. Single-cell RNA-seq highlights intratumoral heterogeneity in primary glioblastoma. Science (2014) doi:10.1126/science.1254257.

23. Resolving tumor evolution: a phylogenetic approach. Journal of the National Cancer Center 4, 97–106 (2024).

24. Dey, S. S., Kester, L., Spanjaard, B., Bienko, M. & van Oudenaarden, A. Integrated genome and transcriptome sequencing of the same cell. Nature Biotechnology 33, 285–289 (2015).

25. Macaulay, I. C. et al. G&T-seq: parallel sequencing of single-cell genomes and transcriptomes. Nature Methods 12, 519–522 (2015).

26. Rodriguez-Meira, A., O’Sullivan, J., Rahman, H. & Mead, A. J. TARGET-Seq: A Protocol for High-Sensitivity Single-Cell Mutational Analysis and Parallel RNA Sequencing. STAR Protocols 1, 100125 (2020).

27. Baysoy, A., Bai, Z., Satija, R. & Fan, R. The technological landscape and applications of single-cell multi-omics. Nature Reviews Molecular Cell Biology 24, 695–713 (2023).

28. Puram, S. V. et al. Single-Cell Transcriptomic Analysis of Primary and Metastatic Tumor Ecosystems in Head and Neck Cancer. Cell 171, 1611–1624.e24 (2017).

29. Peng, J. et al. Single-cell RNA-seq highlights intra-tumoral heterogeneity and malignant progression in pancreatic ductal adenocarcinoma. Cell Research 29, 725–738 (2019).

30. Wu, F. et al. Single-cell profiling of tumor heterogeneity and the microenvironment in advanced non-small cell lung cancer. Nature Communications 12, 1–11 (2021).

31. Xu, K. et al. Single-cell RNA sequencing reveals cell heterogeneity and transcriptome profile of breast cancer lymph node metastasis. Oncogenesis 10, 1–12 (2021).

32. Gao, R. et al. Delineating copy number and clonal substructure in human tumors from single-cell transcriptomes. Nature Biotechnology 39, 599–608 (2021).

33. The hypoxic tumor microenvironment and gene expression. Seminars in Radiation Oncology 14, 207–214 (2004).

34. Lin, W. et al. Single-cell transcriptome analysis of tumor and stromal compartments of pancreatic ductal adenocarcinoma primary tumors and metastatic lesions. Genome Medicine 12, 1–14 (2020).

35. Spatially confined sub-tumor microenvironments in pancreatic cancer. Cell 184, 5577–5592.e18 (2021).

36. Spatially resolved multi-omics deciphers bidirectional tumor-host interdependence in glioblastoma. Cancer Cell 40, 639–655.e13 (2022).

37. Khaliq, A. M. et al. Spatial transcriptomic analysis of primary and metastatic pancreatic cancers highlights tumor microenvironmental heterogeneity. Nature Genetics 56, 2455–2465 (2024).

38. Taylor, B. S. et al. Integrative genomic profiling of human prostate cancer. Cancer cell 18, (2010).

39. Integrated genomic analyses of ovarian carcinoma. Nature 474, 609–615 (2011).

40. Curtis, C. et al. The genomic and transcriptomic architecture of 2,000 breast tumours reveals novel subgroups. Nature 486, 346–352 (2012).

41. Imielinski, M. et al. Mapping the hallmarks of lung adenocarcinoma with massively parallel sequencing. Cell 150, 1107–1120 (2012).

42. Fan, J. et al. Linking transcriptional and genetic tumor heterogeneity through allele analysis of single-cell RNA-seq data. Genome research 28, (2018).

43. Wu, H. et al. Single-cell RNA sequencing reveals tumor heterogeneity, microenvironment, and drug-resistance mechanisms of recurrent glioblastoma. Cancer science 114, (2023).

44. Kaufmann, T. L. et al. MEDICC2: whole-genome doubling aware copy-number phylogenies for cancer evolution. Genome Biology 23, 1–27 (2022).

45. Lu, B., Curtius, K., Graham, T. A., Yang, Z. & Barnes, C. P. CNETML: maximum likelihood inference of phylogeny from copy number profiles of multiple samples. Genome Biology 24, 1–28 (2023).

46. Iacobuzio-Donahue, C. A. et al. DPC4 Gene Status of the Primary Carcinoma Correlates With Patterns of Failure in Patients With Pancreatic Cancer. Journal of Clinical Oncology (2009) doi:10.1200/JCO.2008.17.7188.

47. Yachida, S. et al. Clinical significance of the genetic landscape of pancreatic cancer and implications for identification of potential long-term survivors. Clin Cancer Res 18, 6339–6347 (2012).

48. Thomassen, I. et al. Incidence, prognosis, and possible treatment strategies of peritoneal carcinomatosis of pancreatic origin: a population-based study. Pancreas 42, 72–75 (2013).

49. Wu, L. et al. Clinical significance of site-specific metastases in pancreatic cancer: a study based on both clinical trial and real-world data. Journal of Cancer 12, 1715 (2021).

50. Varghese, A. M. et al. Clinicogenomic landscape of pancreatic adenocarcinoma identifies KRAS mutant dosage as prognostic of overall survival. Nat. Med. 31, 466–477 (2025).

51. Burdziak, C. et al. Epigenetic plasticity cooperates with cell-cell interactions to direct pancreatic tumorigenesis. Science 380, eadd5327 (2023).

52. Dongre, A. & Weinberg, R. A. New insights into the mechanisms of epithelial–mesenchymal transition and implications for cancer. Nature Reviews Molecular Cell Biology 20, 69–84 (2018).

53. Haerinck, J., Goossens, S. & Berx, G. The epithelial–mesenchymal plasticity landscape: principles of design and mechanisms of regulation. Nature Reviews Genetics 24, 590–609 (2023).

54. Gundem, G. et al. The evolutionary history of lethal metastatic prostate cancer. Nature 520, (2015).

55. Maddipati, R. & Stanger, B. Z. Pancreatic Cancer Metastases Harbor Evidence of Polyclonality. Cancer discovery 5, (2015).

56. Oskarsson, T. et al. Breast cancer cells produce tenascin C as a metastatic niche component to colonize the lungs. Nature Medicine 17, 867–874 (2011).

57. Spranger, S., Bao, R. & Gajewski, T. F. Melanoma-intrinsic β-catenin signalling prevents anti-tumour immunity. Nature 523, 231–235 (2015).

58. Hoshino, A. et al. Tumour exosome integrins determine organotropic metastasis. Nature 527, 329–335 (2015).

59. Myc Cooperates with Ras by Programming Inflammation and Immune Suppression. Cell 171, 1301–1315.e14 (2017).

60. Sui, L. et al. PRSS2 remodels the tumor microenvironment via repression of Tsp1 to stimulate tumor growth and progression. Nature Communications 13, 1–19 (2022).

61. Cutler, A. & Breiman, L. Archetypal Analysis. Technometrics 36, 338 (1994).

62. Korem, Y. et al. Geometry of the Gene Expression Space of Individual Cells. PLOS Computational Biology 11, e1004224 (2015).

63. Recovering Gene Interactions from Single-Cell Data Using Data Diffusion. Cell 174, 716–729.e27 (2018).

64. Adler, M., Korem Kohanim, Y., Tendler, A., Mayo, A. & Alon, U. Continuum of Gene-Expression Profiles Provides Spatial Division of Labor within a Differentiated Cell Type. Cell Syst 8, 43–52.e5 (2019).

65. Emergence of division of labor in tissues through cell interactions and spatial cues. Cell Reports 42, 112412 (2023).

66. Otto, D. J., Jordan, C., Dury, B., Dien, C. & Setty, M. Quantifying cell-state densities in single-cell phenotypic landscapes using Mellon. Nature Methods 21, 1185–1195 (2024).

67. Hotspot identifies informative gene modules across modalities of single-cell genomics. Cell Systems 12, 446–456.e9 (2021).

68. Stein, U. et al. The metastasis-associated gene S100A4 is a novel target of beta-catenin/T-cell factor signaling in colon cancer. Gastroenterology 131, (2006).

69. Lo, J.-F. et al. The epithelial-mesenchymal transition mediator S100A4 maintains cancer-initiating cells in head and neck cancers. Cancer Res 71, 1912–1923 (2011).

70. Wang, H. et al. Cofilin 1 induces the epithelial-mesenchymal transition of gastric cancer cells by promoting cytoskeletal rearrangement. Oncotarget 8, 39131 (2017).

71. Li, B. et al. Fibronectin 1 promotes melanoma proliferation and metastasis by inhibiting apoptosis and regulating EMT. OncoTargets and therapy 12, 3207 (2019).

72. Berr, A. L. et al. Vimentin is required for tumor progression and metastasis in a mouse model of non–small cell lung cancer. Oncogene 42, 2074–2087 (2023).

73. Grooteclaes, M. L. & Frisch, S. M. Evidence for a function of CtBP in epithelial gene regulation and anoikis. Oncogene 19, 3823–3828 (2000).

74. Comijn, J. et al. The two-handed E box binding zinc finger protein SIP1 downregulates E-cadherin and induces invasion. Mol Cell 7, 1267–1278 (2001).

75. Sobrado, V. R. et al. The class I bHLH factors E2-2A and E2-2B regulate EMT. Journal of cell science 122, (2009).

76. Cubillo, E. et al. E47 and Id1 Interplay in Epithelial-Mesenchymal Transition. PLOS ONE 8, e59948 (2013).

77. Stemmler, M. P., Eccles, R. L., Brabletz, S. & Brabletz, T. Non-redundant functions of EMT transcription factors. Nature Cell Biology 21, 102–112 (2019).

78. Yang, J. et al. Guidelines and definitions for research on epithelial–mesenchymal transition. Nature Reviews Molecular Cell Biology 21, 341–352 (2020).

79. Kinker, G. S. et al. Pan-cancer single-cell RNA-seq identifies recurring programs of cellular heterogeneity. Nat. Genet. 52, 1208–1218 (2020).

80. Malanchi, I. et al. Interactions between cancer stem cells and their niche govern metastatic colonization. Nature 481, 85–89 (2011).

81. Seubert, B. et al. Tissue inhibitor of metalloproteinases (TIMP)-1 creates a premetastatic niche in the liver through SDF-1/CXCR4-dependent neutrophil recruitment in mice: Seubert, Grünwald et al. Hepatology 61, 238–248 (2015).

82. Nason, S. R. et al. Glucagon-Receptor Signaling Reverses Hepatic Steatosis Independent of Leptin Receptor Expression. Endocrinology 161, bqz013 (2019).

83. Bello-Fernandez, C., Packham, G. & Cleveland, J. L. The ornithine decarboxylase gene is a transcriptional target of c-Myc. Proceedings of the National Academy of Sciences 90, 7804–7808 (1993).

84. Mongiardi, M. P. et al. c-MYC inhibition impairs hypoxia response in glioblastoma multiforme. Oncotarget 7, 33257–33271 (2016).

85. Witkiewicz, A. K. et al. Whole-exome sequencing of pancreatic cancer defines genetic diversity and therapeutic targets. Nature Communications 6, 1–11 (2015).

86. Maddipati, R. et al. Levels Regulate Metastatic Heterogeneity in Pancreatic Adenocarcinoma. Cancer Discov 12, 542–561 (2022).

87. Sancho, P. et al. MYC/PGC-1α balance determines the metabolic phenotype and plasticity of pancreatic cancer stem cells. Cell Metab. 22, 590–605 (2015).

88. Farrell, A. S. et al. MYC regulates ductal-neuroendocrine lineage plasticity in pancreatic ductal adenocarcinoma associated with poor outcome and chemoresistance. Nat Commun 8, 1728 (2017).

89. Silvis, M. R. et al. MYC-mediated resistance to trametinib and HCQ in PDAC is overcome by CDK4/6 and lysosomal inhibition. J. Exp. Med. 220, (2023).

90. Bulle, A. et al. Gemcitabine induces Epithelial-to-Mesenchymal Transition in patient-derived pancreatic ductal adenocarcinoma xenografts. American Journal of Translational Research 11, 765 (2019).

91. CZI Single-Cell Biology Program et al. CZ CELL×GENE Discover: A single-cell data platform for scalable exploration, analysis and modeling of aggregated data. bioRxiv 2023.10.30.563174 (2023) doi:10.1101/2023.10.30.563174.

92. An, M. et al. Early Immune Remodeling Steers Clinical Response to First-Line Chemoimmunotherapy in Advanced Gastric Cancer. Cancer Discov 14, 766–785 (2024).

93. Gavish, A. et al. Hallmarks of transcriptional intratumour heterogeneity across a thousand tumours. Nature 618, 598–606 (2023).

94. Moffitt, R. A. et al. Virtual microdissection identifies distinct tumor- and stroma-specific subtypes of pancreatic ductal adenocarcinoma. Nature Genetics 47, 1168–1178 (2015).

95. Raghavan, S. et al. Microenvironment drives cell state, plasticity, and drug response in pancreatic cancer. Cell 184, 6119–6137.e26 (2021).

96. The Endoplasmic Reticulum Stress Transducer OASIS Is involved in the Terminal Differentiation of Goblet Cells in the Large Intestine. Journal of Biological Chemistry 287, 8144–8153 (2012).

97. Tonelli, C. et al. A mucus production programme promotes classical pancreatic ductal adenocarcinoma. Gut 73, 941–954 (2024).

98. Mehta, A. et al. Immunotherapy Resistance by Inflammation-Induced Dedifferentiation. Cancer Discov 8, 935–943 (2018).

99. Ren, K. et al. Development of the Peritoneal Metastasis: A Review of Back-Grounds, Mechanisms, Treatments and Prospects. Journal of Clinical Medicine 12, 103 (2022).

100. Blastik, M., Plavecz, É. & Zalatnai, A. Pancreatic Carcinomas in a 60-Year, Institute-Based Autopsy Material With Special Emphasis of Metastatic Pattern. Pancreas 40, 478 (2011).

101. Chait, A. & den Hartigh, L. J. Adipose Tissue Distribution, Inflammation and Its Metabolic Consequences, Including Diabetes and Cardiovascular Disease. Front. Cardiovasc. Med. 7, 522637 (2020).

102. Makohon-Moore, A. P. et al. Precancerous neoplastic cells can move through the pancreatic ductal system. Nature 561, 201–205 (2018).

103. Daemen, A. et al. Metabolite profiling stratifies pancreatic ductal adenocarcinomas into subtypes with distinct sensitivities to metabolic inhibitors. Proc Natl Acad Sci U S A 112, E4410–7 (2015).

104. Sunami, Y., Rebelo, A. & Kleeff, J. Lipid Metabolism and Lipid Droplets in Pancreatic Cancer and Stellate Cells. Cancers 10, 3 (2017).

105. Nowell, P. C. The clonal evolution of tumor cell populations. Science 194, 23–28 (1976).

106. Schiffman, J. S. et al. Defining heritability, plasticity, and transition dynamics of cellular phenotypes in somatic evolution. Nat. Genet. 56, 2174–2184 (2024).

107. Mantel, N. The detection of disease clustering and a generalized regression approach. Cancer Res. 27, 209–220 (1967).

108. Corces, M. R. et al. The chromatin accessibility landscape of primary human cancers. Science 362, (2018).

109. D’Ambrosio, A. et al. Increased genomic instability and reshaping of tissue microenvironment underlie oncogenic properties of mutations. Sci Adv 10, eadh4435 (2024).

110. Rozeveld, C. N., Johnson, K. M., Zhang, L. & Razidlo, G. L. KRAS Controls Pancreatic Cancer Cell Lipid Metabolism and Invasive Potential through the Lipase HSL. Cancer Res 80, 4932–4945 (2020).

111. Tadros, S. et al. Lipid Synthesis Facilitates Gemcitabine Resistance through Endoplasmic Reticulum Stress in Pancreatic Cancer. Cancer Res 77, 5503–5517 (2017).

112. Williams, M. J., Werner, B., Barnes, C. P., Graham, T. A. & Sottoriva, A. Identification of neutral tumor evolution across cancer types. Nature Genetics 48, 238–244 (2016).

113. Hayashi, A. et al. A unifying paradigm for transcriptional heterogeneity and squamous features in pancreatic ductal adenocarcinoma. Nat. Cancer 1, 59–74 (2020).

114. Mazutis, L., Masilionis, I. & Chaudhary, O. Frozen tissue dissociation for single-nucleus RNA-Seq protocol metadata. (2020).

115. Shen, R. & Seshan, V. E. FACETS: allele-specific copy number and clonal heterogeneity analysis tool for high-throughput DNA sequencing. Nucleic Acids Res. 44, e131 (2016).

116. Hudson, R. R. & Kaplan, N. L. Statistical properties of the number of recombination events in the history of a sample of DNA sequences. Genetics 111, 147–164 (1985).

117. Kim, K. I. & Simon, R. Using single cell sequencing data to model the evolutionary history of a tumor. BMC Bioinformatics 15, 27 (2014).

118. Malikic, S., Jahn, K., Kuipers, J., Sahinalp, S. C. & Beerenwinkel, N. Integrative inference of subclonal tumour evolution from single-cell and bulk sequencing data. Nat. Commun. 10, 2750 (2019).

119. Satas, G., Zaccaria, S., Mon, G. & Raphael, B. J. SCARLET: Single-cell tumor phylogeny inference with copy-number constrained mutation losses. Cell Syst. 10, 323–332.e8 (2020).

120. Saitou, N. & Nei, M. The neighbor-joining method: a new method for reconstructing phylogenetic trees. Mol. Biol. Evol. 4, 406–425 (1987).

121. Beroukhim, R. et al. The landscape of somatic copy-number alteration across human cancers. Nature 463, 899–905 (2010).

122. Schwarz, G. Estimating the Dimension of a Model. Ann. Statist. 6, (1978).

123. Yang, D. et al. Lineage tracing reveals the phylodynamics, plasticity, and paths of tumor evolution. Cell 185, 1905–1923.e25 (2022).

124. Simonsen, M., Mailund, T. & Pedersen, C. N. S. Rapid neighbour-joining. in Lecture Notes in Computer Science 113–122 (Springer Berlin Heidelberg, Berlin, Heidelberg, 2008).

125. Crowdis, J., He, M. X., Reardon, B. & Van Allen, E. M. CoMut: visualizing integrated molecular information with comutation plots. Bioinformatics 36, 4348–4349 (2020).

126. Azizi, E. et al. Single-cell map of diverse immune phenotypes in the breast tumor microenvironment. Cell 174, 1293–1308.e36 (2018).

127. Slyper, M. et al. A single-cell and single-nucleus RNA-Seq toolbox for fresh and frozen human tumors. Nat. Med. 26, 792–802 (2020).

128. Young, M. D. & Behjati, S. SoupX removes ambient RNA contamination from droplet-based single-cell RNA sequencing data. Gigascience 9, giaa151 (2020).

129. Fleming, S. J. et al. Unsupervised removal of systematic background noise from droplet-based single-cell experiments using CellBender. Nat. Methods 20, 1323–1335 (2023).

130. Stuart, T. et al. Comprehensive integration of single-cell data. Cell 177, 1888–1902.e21 (2019).

131. Heumos, L. et al. Best practices for single-cell analysis across modalities. Nat. Rev. Genet. 24, 550–572 (2023).

132. Drokhlyansky, E. et al. The human and mouse Enteric nervous system at single-cell resolution. Cell 182, 1606–1622.e23 (2020).

133. Neavin, D. et al. Demuxafy: improvement in droplet assignment by integrating multiple single-cell demultiplexing and doublet detection methods. Genome Biol. 25, 94 (2024).

134. Gayoso, A., Shor, J., Carr, A. J., Sharma, R. & Pe’er, D. GitHub: DoubletDetection. (Zenodo, 2019). doi:10.5281/ZENODO.2678042.

135. Wolock, S. L., Lopez, R. & Klein, A. M. Scrublet: Computational identification of cell Doublets in Single-cell transcriptomic data. Cell Syst. 8, 281–291.e9 (2019).

136. Xi, N. M. & Li, J. J. Benchmarking computational doublet-detection methods for single-cell RNA sequencing data. Cell Syst. 12, 176–194.e6 (2021).

137. Wolf, F. A., Angerer, P. & Theis, F. J. SCANPY: large-scale single-cell gene expression data analysis. Genome Biol. 19, 15 (2018).

138. Satija, R., Farrell, J. A., Gennert, D., Schier, A. F. & Regev, A. Spatial reconstruction of single-cell gene expression data. Nat. Biotechnol. 33, 495–502 (2015).

139. Cheng, D. T. et al. Memorial Sloan Kettering-Integrated Mutation Profiling of Actionable Cancer Targets (MSK-IMPACT): A hybridization capture-based next-generation sequencing clinical assay for solid tumor molecular oncology. J. Mol. Diagn. 17, 251–264 (2015).

140. Gaveau, B. & Schulman, L. S. Multiple phases in stochastic dynamics: geometry and probabilities. Phys. Rev. E Stat. Nonlin. Soft Matter Phys. 73, 036124 (2006).

141. Wolf, F. A. et al. PAGA: graph abstraction reconciles clustering with trajectory inference through a topology preserving map of single cells. Genome Biol. 20, 59 (2019).

142. Mootha, V. K. et al. PGC-1alpha-responsive genes involved in oxidative phosphorylation are coordinately downregulated in human diabetes. Nat. Genet. 34, 267–273 (2003).

143. Subramanian, A. et al. Gene set enrichment analysis: a knowledge-based approach for interpreting genome-wide expression profiles. Proc. Natl. Acad. Sci. U. S. A. 102, 15545–15550 (2005).

144. Fang, Z., Liu, X. & Peltz, G. GSEApy: a comprehensive package for performing gene set enrichment analysis in Python. Bioinformatics 39, btac757 (2023).

145. Hwang, W. L. et al. Single-nucleus and spatial transcriptome profiling of pancreatic cancer identifies multicellular dynamics associated with neoadjuvant treatment. Nat. Genet. 54, 1178–1191 (2022).

146. Nowotschin, S. et al. The emergent landscape of the mouse gut endoderm at single-cell resolution. Nature 569, 361–367 (2019).

147. Tirosh, I. et al. Dissecting the multicellular ecosystem of metastatic melanoma by single-cell RNA-seq. Science 352, 189–196 (2016).

148. Finak, G. et al. MAST: a flexible statistical framework for assessing transcriptional changes and characterizing heterogeneity in single-cell RNA sequencing data. Genome Biol. 16, 278 (2015).

149. Abdulla, S. et al. CZ CELL×GENE Discover: A single-cell data platform for scalable exploration, analysis and modeling of aggregated data. Cell Biology (2023).

150. Ogata, H., Goto, S., Fujibuchi, W. & Kanehisa, M. Computation with the KEGG pathway database. Biosystems. 47, 119–128 (1998).

